# Spatial Topography of Individual-Specific Cortical Networks Predicts Human Cognition, Personality and Emotion

**DOI:** 10.1101/213041

**Authors:** Ru Kong, Jingwei Li, Csaba Orban, Mert R Sabuncu, Hesheng Liu, Alexander Schaefer, Nanbo Sun, Xi-Nian Zuo, Avram J. Holmes, Simon B. Eickhoff, B.T. Thomas Yeo

## Abstract

Resting-state functional magnetic resonance imaging (rs-fMRI) offers the opportunity to delineate individual-specific brain networks. A major question is whether individual-specific network topography (i.e., location and spatial arrangement) is behaviorally relevant. Here, we propose a multi-session hierarchical Bayesian model (MS-HBM) for estimating individual-specific cortical networks and investigate whether individual-specific network topography can predict human behavior. The multiple layers of the MS-HBM explicitly differentiate intra-subject (within-subject) from inter-subject (between-subject) network variability. By ignoring intra-subject variability, previous network mappings might confuse intra-subject variability for inter-subject differences. Compared with other approaches, MS-HBM parcellations generalized better to new rs-fMRI and task-fMRI data from the same subjects. More specifically, MS-HBM parcellations estimated from a single rs-fMRI session (10 minutes) showed comparable generalizability as parcellations estimated by two state-of-the-art methods using five sessions (50 minutes). We also showed that behavioral phenotypes across cognition, personality and emotion could be predicted by individual-specific network topography with modest accuracy, comparable to previous reports predicting phenotypes based on connectivity strength. Network topography estimated by MS-HBM was more effective for behavioral prediction than network size, as well as network topography estimated by other parcellation approaches. Thus, similar to connectivity *strength*, individual-specific network *topography* might also serve as a fingerprint of human behavior.

## Introduction

One prominent tool for identifying large-scale human brain networks is resting-state functional connectivity (RSFC), which reflects the synchrony of fMRI signals between brain regions, while a subject is lying at rest without any goal-directed task (Biswal et al., 1995; Greicius et al. 2003; Fox and Raichle, 2007). RSFC brain networks correspond well to task-evoked activation patterns (Seeley et al. 2007; Smith et al., 2009; Cole et al., 2014; Yeo et al., 2015a). RSFC is also heritable (Glahn et al. 2010; Yang et al. 2016; Ge et al., 2017), correlates with gene expression across the cortical mantle (Hawrylycz et al. 2015; Richiardi et al. 2015; Krienen et al. 2016), and predicts individual differences in behavior (Hampson et al., 2006; van den Heuvel et al., 2009; Finn et al., 2015; Smith et al., 2015). Consequently, RSFC has been widely utilized to estimate population-average functional brain networks by averaging data across multiple subjects (Beckmann et al. 2005; Damoiseaux et al. 2006; Fox et al. 2006; Dosenbach et al. 2007; Margulies et al. 2007; Power et al. 2011; Yeo et al., 2011; Lee et al. 2012).

Population-average networks have provided important insights into the large-scale functional organization of the human brain (Buckner et al., 2013; Wig et al., 2017). However, since population-average networks might obscure individual-specific network organization, there is significant interest in estimating individual-specific brain networks (Beckmann et al., 2009; Bellec et al., 2010; Zuo et al., 2010; Varoquaux et al., 2011; Hacker et al., 2013; Wig et al., 2014; Chong et al., 2017). Indeed, many studies have documented that the size, location and spatial arrangement of individual-specific brain networks vary substantially across participants (Harrison et al., 2015; Laumann et al., 2015; Wang et al., 2015; Glasser et al., 2016; Gordon et al., 2017a, 2017b, 2017c; Braga and Buckner, 2017). Yet, the possible behavioral relevance of individual differences in network size and network topography (location and spatial arrangement) remains largely unclear.

We proposed a multi-session hierarchical Bayesian model (MS-HBM) for estimating individual-specific network parcellations of the cerebral cortex and investigated whether individual-specific network topography and size were associated with human behavior. The multiple layers of the MS-HBM allowed explicit separation of inter-subject (between-subject) and intra-subject (within-session) functional connectivity variability. Previous individual-specific network mappings only accounted for inter-subject variability, but not intra-subject variability. However, inter-subject and intra-subject RSFC variability can be markedly different across regions (Mueller et al., 2013; Chen et al., 2015; Laumann et al., 2015). For example, the motor cortex exhibits high intra-subject functional connectivity variability, but low inter-subject functional connectivity variability (Laumann et al., 2015). Therefore, observed RSFC variability in the motor cortex might be incorrectly attributed to inter-subject variability of brain networks, rather than just intra-subject sampling variability, resulting in sub-optimal network mapping. In this paper, we showed that compared with four other approaches, MS-HBM individual-specific parcellations generalized better to new resting and task fMRI data from the same individuals.

Having established that the MS-HBM generated high-quality individual-specific parcellations, we investigated whether individual differences in network topography (i.e., location and spatial arrangement) and size could predict behavioral measures across cognition, personality and emotion. While there is a plethora of studies associating regional brain volumes and anatomical patterns with behavior (e.g., Erickson et al., 2011; Holmes et al., 2016; Sabuncu et al., 2016; Cachia et al., 2017), there are relatively few studies relating topography and size of functional areas (or networks) with behavior or other traits (Dehaene et al., 2010; Bijsterbosch et al., 2018; Salehi et al., 2018). Using kernel regression, we showed that multiple behavioral measures could be predicted with modest accuracy. Furthermore, network topography estimated by MS-HBM achieved better prediction accuracies than topography estimated by other parcellation approaches. Lastly, we found that at least at the resolution of large-scale networks, network topography was more useful than network size in predicting behavior.

The contributions of this work are two-fold. First, by estimating inter-subject variability, intra-subject variability and individual-specific networks within a unified statistical framework, the estimation of individual-specific networks were greatly improved. For example, MS-HBM parcellations estimated from a *single* rs-fMRI session were comparable to those generated by two prominent algorithms using five times the amount of data (Wang et al., 2015; Gordon et al., 2017a; 2017b), as evaluated by generalizability to new rs-fMRI data from the same subjects. Second, our results suggest that individual-specific cortical network topography might serve as a fingerprint of human behavior, which might complement the usage of functional connectivity strength in the vast majority of previous literature (Hampson et al., 2006; Finn et al., 2015; Rosenberg et al., 2016; Smith et al., 2015; Yeo et al., 2015b, Nostro et al., 2018). This highlights the importance of considering both network topography and functional connectivity strength for behavioral prediction.

## Methods

### Overview

We proposed a multi-session hierarchical Bayesian model (MS-HBM) to estimate functional network parcellations of the cerebral cortex in individual subjects. The model distinguished between inter-subject and intra-subject network variability. Subsequent analyses proceeded in four stages. First, to examine whether inter-subject and intra-subject variability could be reliably estimated across datasets, the MS-HBM was applied to three multi-session rs-fMRI datasets. Second, the MS-HBM was compared with four other approaches using new rs-fMRI and task-fMRI data from the same subjects. Third, we examined the reproducibility of the MS-HBM parcellations and how well the parcellations captured inter-subject differences. Finally, we investigated whether individual differences in cortical parcellations reflected individual differences in behavior.

### Multi-session rs-fMRI datasets

The Genomic Superstruct Project (GSP) test-retest dataset (Holmes et al., 2015) consisted of structural MRI and rs-fMRI from 69 healthy young adults (ages 18 to 35). All imaging data were collected on matched 3T Tim Trio scanners (Siemens Healthcare, Erlangen, Germany) at Harvard University and Massachusetts General Hospital using the vendor-supplied 12-channel phased-array head coil. Each participant has two sessions, acquired on two different days separated by less than 6 months. One or two rs-fMRI runs were acquired per session. Each BOLD run was acquired in 3mm isotropic resolution with a TR of 3.0s and lasted for 6min and 12s. The structural data consisted of one 1.2mm isotropic scan for each session. Details of the data collection can be found elsewhere (Holmes et al., 2015).

The Hangzhou Normal University of the Consortium for Reliability and Reproducibility (CoRR-HNU) multi-session dataset (Zuo et al., 2014; Chen et al., 2015) consisted of structural MRI and rs-fMRI from 30 young healthy adults (ages 20 to 30). All imaging data were collected on a 3T GE Discovery MR750 scanner using an 8-channel head coil. Each participant was scanned for a total of 10 sessions across one month (one session every three days). One rs-fMRI run was collected in each session. Each fMRI run was acquired in 3.4mm isotropic resolution with a TR of 2.0s and lasted for 10min. The structural data consisted of one 1mm isotropic scan for each session. Details of the data collection can be found elsewhere (Zuo et al., 2014; Chen et al., 2015).

The Human Connectome Project (HCP) S900 release (Van Essen et al., 2012b; Smith et al., 2013) consisted of structural MRI, rs-fMRI and task fMRI of 881 subjects. All imaging data were collected on a custom-made Siemens 3T Skyra scanner using a multiband sequence. Each participant went through two fMRI sessions on two consecutive days. Two rs-fMRI runs were collected in each session. Each fMRI run was acquired in 2mm isotropic resolution with a TR of 0.72s and lasted for 14min and 33s. The structural data consisted of one 0.7mm isotropic scan for each subject. Details of the data collection can be found elsewhere (Van Essen et al., 2012b; Smith et al., 2013).

It is worth noting the significant acquisition differences among the three datasets, including scanner type (e.g., GE versus Siemens) and acquisition sequence (e.g., multiband versus non-multiband). The interval between repeated visits were also very different, ranging from one day in the HCP dataset and up to six months in the GSP dataset. These differences allowed us to test the robustness of our individual-specific network estimation model.

### Preprocessing

Processing of GSP and CoRR-HNU data followed the surface-based pipeline of Yeo and colleagues (Yeo et al., 2011; Holmes et al., 2015) using a combination of FreeSurfer (Dale et al., 1999; Fischl et al. 1999a; 1999b; Fischl et al., 2001; Ségonne et al., 2007; Greve and Fischl 2009) and FSL (Jenkinson et al., 2002; Smith et al., 2004), with additional censoring steps pioneered by the Petersen’s group (Power et al., 2014; Gordon et al., 2016). The final preprocessed data were on the FreeSurfer fsaverage5 surface space (4mm vertex spacing). More details can be found in Supplementary Methods S1.

In the case of the HCP data, we utilized the MSMAll ICA-FIX data on fs_LR32K surface space (HCP S900 manual; Van Essen et al. 2012a; Van Essen et al. 2012b; Glasser et al. 2013; Smith et al. 2013; Griffanti et al., 2014; Salimi-Khorshidi et al. 2014) with additional nuisance regression, censoring (Burgess et al., 2016; Siegel et al., 2016) and spatial smoothing. More details can be found in Supplementary Methods S2.

### Population-level parcellation and functional connectivity profiles

We have previously developed an approach to derive a population-level parcellation of the cerebral cortex into large-scale resting-state networks (Yeo et al., 2011). The term “parcellation” is used to indicate that every cortical location is assigned a label (Fischl et al., 2004; Yeo et al., 2011), rather than delineating the locations of a specific brain function or network. The cortical networks were defined as sets of cortical regions with similar corticocortical functional connectivity profiles. Here we applied the same approach to the GSP, CoRR-HNU and HCP datasets. Our previous analyses (Yeo et al., 2011) identified 7 and 17 networks to be particularly stable. For simplicity, we will only consider 17 networks throughout this paper. Details of this approach have been previously described (Yeo et al., 2011). For completeness, we will briefly describe its application to the current datasets.

Recall that the preprocessed fMRI data from the CoRR-HNU and GSP subjects have been projected onto the fsaverage5 surface meshes. The fsaverage5 surface meshes consisted of 18715 cortical vertices. Following previous work (Yeo et al., 2011), the connectivity profile of a cortical region (vertex) was defined to be its functional coupling to 1175 regions of interest (ROIs). The 1175 ROIs consisted of single vertices uniformly distributed across the fsaverage5 surface meshes. For each rs-fMRI run of each subject, the Pearson’s correlation between the fMRI time series at each spatial location (18715 vertices) and the 1175 ROIs were computed. The 18715 x 1175 correlation matrix were then binarized by keeping the top 10% of the correlations to obtain the final functional connectivity profiles. Outlier volumes (flagged during preprocessing) were ignored when computing the correlations.

In the case of the HCP dataset, the preprocessed fMRI data have been projected onto the fs_LR surface space. The fs_LR32K surface meshes consisted of 59412 cortical vertices. We defined the connectivity profile of a cortical region (vertex) to be its functional coupling to 1483 ROIs. The 1483 ROIs consisted of single vertices uniformly distributed across the fs_LR32K surface meshes. For each rs-fMRI run of each subject, the Pearson’s correlation between the fMRI time series at each spatial location (59412 vertices) and the 1483 ROIs were computed. The 59412 × 1483 correlation matrix were then binarized by keeping the top 10% of the correlations to obtain the final functional connectivity profile. Outlier volumes were again ignored when computing the correlations.

To obtain a population-level parcellation from a group of subjects, each vertex’s connectivity profiles were averaged across all BOLD runs of all subjects. The averaged connectivity profiles were then clustered using a mixture of von Mises–Fisher distributions (Lashkari et al., 2010; Yeo et al., 2011).

### Multi-session hierarchical Bayesian model (MS-HBM)

The previous section described an approach to estimate a population-level parcellation from a group of subjects. Figure 1 illustrates the MS-HBM model for estimating individual-specific cerebral cortex parcellations using multi-session fMRI data. Let 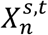 denote the (binarized) functional connectivity profile of cortical vertex *n* from session *t* of subject *s*. For example, Figure 1 (fourth row) illustrates the binarized functional connectivity profile for a posterior cingulate cortex vertex 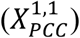 and a precuneus vertex 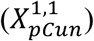 from the 1st session of the 1st subject. The shaded circle indicates that 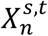 is the only observation in the entire model. Based on the observed connectivity profiles of *all* vertices from *all* sessions of a single subject, the goal is to assign a network label 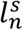 for each vertex *n* of subject *s*. Even though a vertex’s connectivity profile is unlikely to be the same across different fMRI sessions, the vertex’s network label was assumed to be the same across sessions. For example, Figure 1 (last row) illustrates the individual-specific parcellation of the 1st subject using data from *all* sessions.

**Figure 1.**
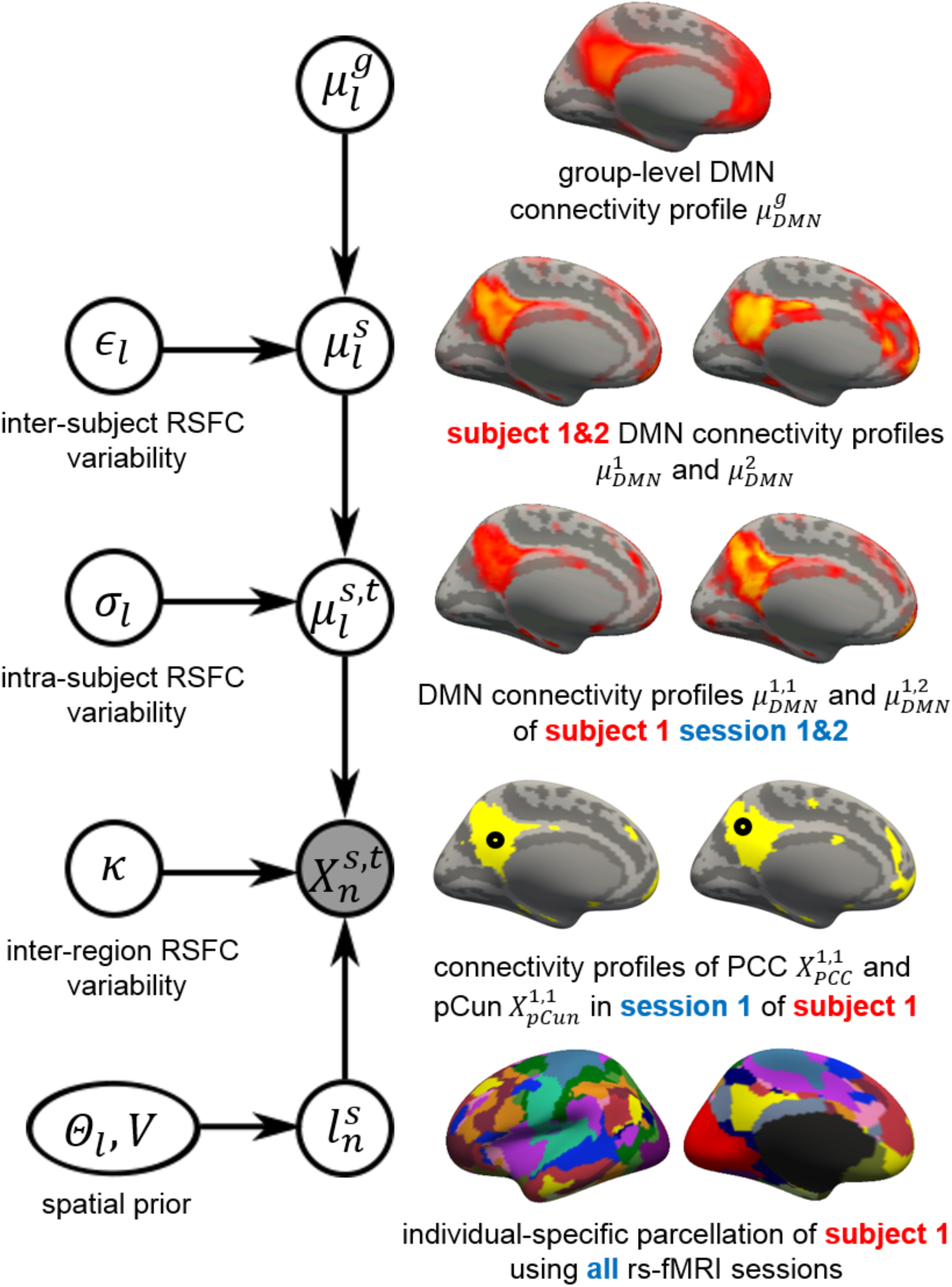
Multi-session hierarchical Bayesian model (MS-HBM) of individual-specific cortical parcellation. 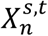 denote the RSFC profile at brain location *n* of subject *s* during rs-fMRI session *t*. The shaded circle indicates that 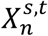 are the only observed variables in the entire model. The goal is to estimate the network label 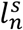 for subject *s* at each cortical location *n* given RSFC profiles from all sessions. 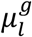 is the group-level RSFC profile of network *l*. 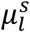 is the subject-specific RSFC profile of network *l*. A large *∈*_*l*_ indicates small inter-subject RSFC variability, i.e., the group-level and subject-specific RSFC profiles are very similar. 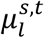 is the subject-specific RSFC profile of network *l* during session t. A large *σ*_*l*_ indicates small intra-subject RSFC variability, i.e., the subject-level and session-level RSFC profiles are very similar. κ captures inter-region RSFC variability. A large κ indicates small inter-region variability, i.e., two regions from the same network exhibit very similar RSFC profiles. Finally, *Θ*_*l*_ captures inter-subject variability in the spatial distribution of networks (e.g., high probability of the DMN being located at the PCC), while smoothness prior *V* encourages network labels to be spatially smooth. See text for details.

Some of the model parameters (e.g., inter-subject variability) must be estimated from a training set of subjects. A new subject (possibly from another dataset) could then be parcellated without access to the original training data. Even though the model was defined on multi-session fMRI data, an effective workaround (details below) was provided for single-session fMRI data. The exact mathematical model is found in Supplemental Methods S3. Here we provide the intuition behind this model.

To obtain the subject-specific parcellation, the MS-HBM assumes that each cortical network exhibits a distinctive RSFC profile. Let 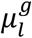 denote the group-level functional connectivity profile of network *l*. We can think of 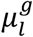 as the average connectivity profile of all vertices of all sessions of all subjects belonging to network *l*. For example, Figure 1 (top row) illustrates the group-level default mode network (DMN) connectivity profile 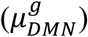.

To model inter-subject RSFC variability, let 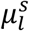 denote the functional connectivity profile of network *l* and subject *s*. We can think of 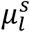 as the average connectivity profile of all vertices of all sessions of subject *s* belonging to network *l*. For example, Figure 1 (second row) illustrates the DMN connectivity profiles of two subjects (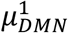 and 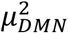. The subject-specific connectivity profile 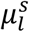 was assumed to follow a von Mises-Fisher distribution with mean direction 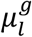 (group-level RSFC profile of network *l*) and concentration parameter *∈*_*l*_. A large *∈*_*l*_ indicates low inter-subject functional connectivity variability, i.e., 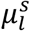 and 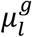 are very similar. The subscript *l* indicates that *∈*_*l*_ is different for each network.

To model intra-subject RSFC variability, let 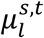 denote the functional connectivity profile of network *l* and subject *s* during session *t*. We can think of 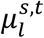 as the average connectivity profile of all vertices from session *t* of subject *s* belong to network *l*. For example, Figure 1 (third row) illustrates the DMN connectivity profiles of subject 1 during sessions 1 and 2 (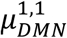 and 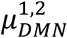). The session-specific connectivity profiles 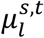 was assumed to follow a von Mises-Fisher distribution with mean direction 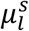 (subject-specific RSFC profile) and concentration parameter *σ*_*l*_. A large *σ*_*l*_ indicates low intra-subject functional connectivity variability, i.e., 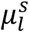 and 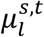 profiles are very similar. The subscript *l* indicates that *σ*_*l*_ is different for each network.

The observed connectivity profiles of two regions belonging to the same network are unlikely to be identical. For example, the connectivity profiles of PCC and precuneus are similar, but not identical (Figure 1 fourth row) even though they might both belong to the DMN. To account for this intra-network (inter-region) variability, the observed connectivity profile 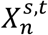 of vertex *n* (which has been assigned to network *l*) was assumed to follow a von Mises-Fisher distribution with mean direction 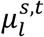 (session-specific and subject-specific connectivity profile of network *l*) and concentration parameter κ. A large κ indicates low inter-region functional connectivity variability. Here, κ was set to be the same across networks (see Supplementary Methods S3 for justification).

Given the previous modeling assumptions, if the observed connectivity profile 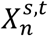 of vertex *n* were most similar to the session-specific and subject-specific connectivity profile 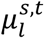 of the DMN (where similarity is measured via the likelihood of the *von Mises-Fisher* distribution), then vertex *n* would be assigned to the DMN. Therefore, at this point, the model is somewhat similar to the population-level parcellation approach (Yeo2011), except that the population-level approach does not account for intra-subject and inter-subject RSFC variability. Furthermore, unlike group averaged connectivity profiles, the observed functional connectivity profiles of individual subjects are generally very noisy. If the observed profiles of PCC and pCun were too noisy, the model might not assign both of them to the DMN. Therefore, additional priors were imposed on the parcellation. First, the spatial smoothness prior *V* encourages neighboring vertices (e.g., PCC and pCun) to be assigned to the same network. Second, the spatial prior *Θ*_*l,n*_ denotes the probability of network *l* occurring at a particular spatial location *n*. For example, PCC might have high prior probability of being assigned to the DMN.

Given a dataset of subjects with multi-session rs-fMRI data, the group-level network connectivity profiles 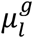, the inter-subject functional connectivity variability *∈*_*l*_, the intra-subject functional connectivity variability *σ*_*l*_, the spatial smoothness prior *V* and the inter-subject spatial variability prior *Θ*_*l*_ could be estimated. The estimated group-level priors 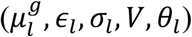 could then be used to parcellate a new subject. Here we utilized a variational Bayes Expectation-Maximization (VBEM) algorithm to learn the group-level priors from the training data and to estimate individual-specific parcellations. Details of the VBEM algorithm can be found in Supplementary Methods S4.

Although the MS-HBM was formulated for multi-session fMRI data, most studies only collect a single run of fMRI data. We considered the ad-hoc approach of splitting the single fMRI run into two and treating the resulting runs as two separate sessions. Our evaluations (see Results) suggest that this workaround worked surprisingly well.

### Characterizing inter-subject and intra-subject network variability

We first evaluate whether the MS-HBM can yield robust estimates of inter-subject and intra-subject variability across datasets. For the purpose of subsequent experiments, the GSP dataset was divided into training (N = 37) and validation (N = 32) sets. The CoRR-HNU dataset (N = 30) was kept unchanged. The HCP dataset was divided into training (N = 40), validation (N = 40) and test (N = 596) sets. Furthermore, different fMRI runs within the same session were treated as data from different sessions. For example, each HCP subject underwent two fMRI sessions on two consecutive days. Within each session, there were two rs-fMRI runs. For the purpose of our analyses, we treated each HCP subject as having four sessions of data. Future work might differentiate between intra-session and inter-session variability.

The group-level parcellation algorithm was applied to the GSP training set. The resulting group-level parcellation was then used to initialize the estimation of the group-level network connectivity profiles 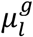, the inter-subject functional connectivity variability *∈*_*l*_, the intra-subject functional connectivity variability *σ*_*l*_, and the inter-subject spatial variability prior *Θ*_*l*_. For the purpose of computational efficiency, the spatial smoothness prior *V* was ignored in this analysis (see Supplementary Methods S4.1 for justification). The procedure was repeated for the CoRR-HNU dataset and HCP training set.

### Comparison with alternative approaches

Having confirmed previous literature (Mueller et al., 2013; Laumann et al., 2015) that inter-subject and intra-subject functional connectivity variability were indeed different across cortical networks, we compared MS-HBM with four alternative approaches. The first approach was to apply the population-level parcellation (Yeo et al., 2011) to individual subjects, which we will refer to as “Yeo2011”. The second approach is “YeoBackProject”, which is analogous to the ICA back-projection algorithm (Calhoun et al., 2009; Beckmann et al., 2009; Filippini et al., 2009; Zuo et al., 2010; Calhoun and Adali 2012). The third approach is the influential individual-specific parcellation algorithm of Gordon and colleagues (Gordon et al., 2017a; Gordon et al., 2017b), which we will refer to as “Gordon2017”. The fourth approach is the prominent individual-specific parcellation algorithm of Wang and colleagues (Wang et al., 2015), which we will refer to as “Wang2015”. See Supplementary Methods S5 for more details.

All algorithms were applied to the CoRR-HNU dataset and the HCP test set. In the case of the CoRR-HNU dataset, the model parameters of all algorithms were estimated from the GSP dataset and then utilized to infer the parcellations of CoRR-HNU subjects. This is important because inter-subject and intra-subject variability might differ across datasets, so it was important to evaluate whether MS-HBM model parameters estimated from one dataset could be generalized to another dataset. More specifically, the training procedure for the MS-HBM was the same as the previous section, except that the GSP validation set was also used to tune the spatial smoothness prior *V*. Similarly, “free” parameters in Wang2015 and Gordon2017 were tuned using the GSP validation set.

In the case of the HCP dataset, recall that the HCP data were in a different surface space from the GSP data, so the GSP model parameters could not be applied to the HCP subjects. Instead, the model parameters of all algorithms were estimated from the HCP training and validation sets, and then utilized to infer the parcellation of each subject in the HCP test set.

### Quantitative evaluation measures

Evaluating the quality of individual-specific resting-state parcellations is difficult because of a lack of ground truth. Here, we considered two common evaluation metrics utilized in previous studies (Gordon et al., 2016; Chong et al., 2017; Gordon et al., 2017c; Schaefer et al., in press): resting-state connectional homogeneity and task functional inhomogeneity measures. These metrics encode the principle that if an individual-specific parcellation captured the system-level organization of the individual’s cerebral cortex, then each network should have homogeneous connectivity and function:

1. Resting-state connectional homogeneity. Resting-state connectional homogeneity was computed by averaging the Pearson’s correlations between the rs-fMRI time courses of all pairs of vertices within each network (Schaefer et al., in press). The average correlations were then averaged across all networks while accounting for network size:

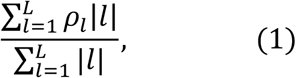

where *ρ*_*l*_ is the resting-state homogeneity of network *l* and |*l*| is the number of vertices within network *l* (Schaefer et al., in press). For each subject from CoRR-HNU (N = 30) and HCP test set (N = 596), we used one session to infer the individual-specific parcellation and computed the resting-state homogeneity of the individual-specific parcellation with the remaining sessions. Because the HNU dataset has the most amount of data (100 min), we also parcellated each CoRR-HNU subject using one or more rs-fMRI sessions and evaluated the resting-state homogeneity with the remaining sessions. This allowed us to estimate how much the various algorithms would improve with more data. When comparing between parcellations, the effect size (Cohen’s d) of differences was computed. It is worth emphasizing that the evaluation utilized new rs-fMRI data not used for estimating the individual-specific parcellations.
2. Task functional inhomogeneity. The HCP task-fMRI data consisted of seven functional domains: social cognition, motor, gambling, working memory, language processing, emotional processing and relational processing, each with multiple task contrasts (Barch et al., 2013). For a given task contrast, task inhomogeneity was defined as the standard deviation of (activation) z-values within each network (Gordon et al., 2017c; Schaefer et al., in press). A lower standard deviation indicated higher functional homogeneity within the network. The standard deviations were averaged across all networks while accounting for network size:

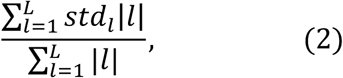

where *std*_*l*_ is the standard deviation of task activation z-values for network *l* and |*l*| is the number of vertices in parcel *l* (Gordon et al., 2017c; Schaefer et al., in press). For each subject in the HCP test set (N = 596), the first rs-fMRI run from the first session was used to infer the individual-specific parcellation. The individual-specific parcellation was then utilized to evaluate task inhomogeneity for each task contrast (Eq. (2)) and then averaged across all contrasts within a functional domain, resulting in a single functional inhomogeneity measure per functional domain. The number of task contrasts per functional domain ranged from three for the emotion domain to eight for the working memory domain. When comparing between parcellations, the inhomogeneity metric (Eq. (2)) was averaged across all contrasts within a functional domain before the effect size (Cohen’s d) of differences was computed for each functional domain.

We note that a cortical parcellation with more networks would on average perform better on the proposed evaluation metrics. The reason is that a cortical parcellation with more networks will have smaller networks (on average), resulting in higher connectional homogeneity and lower functional inhomogeneity. For example, if a network consisted of only two vertices, then it would be highly homogeneous. However, this was not an issue in our experiments because all approaches were constrained to estimate the same number of networks (i.e., 17 networks). Furthermore, the evaluation metrics (Equations (1) and (2)) accounted for network size, so a network with only two vertices would only contribute minimally to the final homogeneity metric.

### Intra-subject reproducibility and inter-subject similarity of MS-HBM network topography

Having established that the MS-HBM was better than other approaches in generating individual-specific parcellations, the reproducibility of individual-specific MS-HBM networks was further characterized using the CoRR-HNU data and HCP test set. Given that intra-subject and inter-subject network variability were different across networks, we were interested in evaluating whether intra-subject network reproducibility and inter-subject network similarity were also different across networks.

Individual-specific MS-HBM parcellations were independently inferred using the first two runs and the last two runs of the HCP test set. Therefore, there were two individual-specific parcellations for each subject based on data from two independent sets of rs-fMRI data. MS-HBM parcellations were also independently inferred using sessions 1-5 and sessions 6-10 of the CoRR-HNU dataset. Therefore, there were two individual-specific parcellations for each subject based on data from two independent sets of five sessions.

To evaluate the reproducibility of individual-specific parcellations, the Dice coefficient was computed for each network from the two parcellations of each subject. The Dice coefficients were then averaged across all networks and all subjects to provide an overall measure of intra-subject parcellation reproducibility. To evaluate inter-subject parcellation similarity, for each pair of subjects, the Dice coefficient was computed for each network. Since there were two parcellations for each subject, there were a total of four Dice coefficients for each network, which were then averaged. The Dice coefficients were then averaged across all networks and all pairs of subjects to provide an overall measure of inter-subject parcellation similarity.

### HCP behavioral data

Given that individual-specific functional networks exhibited unique topographical features not observed in group-level networks, we further investigated whether the spatial configuration of individual-specific cortical parcellations was behaviorally meaningful. Since the HCP dataset has a rich repertoire of behavioral data, we selected 58 behavioral phenotypes measuring cognition, personality and emotion (Table S1). 18 subjects were excluded from further analyses because they did not have all behavioral phenotypes, resulting in a final set of 577 subjects. Individual-specific MS-HBM parcellations were estimated for each HCP test subject (N = 577) using all four rs-fMRI runs, where each run was treated as coming from an independent session. We note that very similar parcellations were obtained if we averaged the connectivity profiles across the two fMRI runs within each day, treating each day as an independent session.

Because the 58 behavioral measures were correlated, we also considered a subset of five minimally correlated behavioral measures. The five behavioral measures were selected as follows. We randomly picked a pair of behavioral measures with an absolute correlation of less than 0.1. Three more behavioral measures were added one at a time, while ensuring that each newly added behavioral measure was minimally correlated (absolute r < 0.1) with the current set of behavioral measures. This procedure was repeated 100 times, resulting in 100 sets of five behavioral measures. The behavioral set with the smallest maximum absolute correlation was selected. The final set of five behavioral measures corresponded to reading (pronunciation), positive affect, grip strength, social cognition (random) and contrast sensitivity (Table S1). The maximum absolute correlation was r = 0.068 (p = 0.104, which is not significant even if we ignore multiple comparisons).

### Can individual-specific network spatial topography be used to predict behavior?

Kernel regression (Murphy et al., 2012) was utilized to predict each behavioral phenotype in individual subjects. Suppose *y* is the behavioral measure (e.g., fluid intelligence) and *l* is the individual-specific parcellation of a test subject. In addition, suppose *y*_*i*_ is the behavioral measure (e.g., fluid intelligence) and *l*_*i*_ is the individual-specific parcellation of the *i*-th training subject. Then kernel regression would predict the behavior of the test subject as the weighted average of the behaviors of the training subjects: *y* ≈ ∑_*i*∈training set_ Similarity(*l*_*i*_, *l*)*y*_*i*_, where Similarity(*l*_*i*_, *l*) was the Dice overlap coefficient between corresponding networks of the test subject and *i*-th training subject, averaged across 17 networks. Therefore, successful prediction would indicate that subjects with more spatially overlapping networks (i.e., network topography) have similar behavioral measures.

In practice, we included an *l*_2_-regularization term (i.e., kernel ridge regression) to reduce overfitting (Supplementary Methods S6; Murphy et al., 2012). The *l*_2_-regularization parameter was determined via an inner-loop cross-validation procedure. More specifically, we performed 20-fold cross-validation for each behavioral phenotype. Care was taken so that family members were not split between folds. For each test fold, 20-fold cross-validation was repeatedly applied to the remaining 19 folds with different regularization parameters (i.e., inner-loop cross-validation). The optimal regularization parameter from the inner-loop cross-validation was then used to predict the behavioral phenotype in the test fold. Accuracy was measured by correlating the predicted and actual behavioral measure across all subjects within the test fold (Finn et al., 2015). By repeating the procedure for each test fold, each behavior yielded 20 correlation accuracies, which were then averaged across the 20 folds. Because a single 20-fold cross-validation might be sensitive to the particular split of the data into folds (Varoquaux et al., 2017), the above 20-fold cross-validation was repeated 100 times. The mean accuracy and standard deviation across the 100 cross-validations will be reported.

Finally, certain behavioral measures are known to correlate with motion (Siegel et al., 2016). Therefore, age, sex, framewise displacement (FD), DVARS, body mass index and total brain volume were regressed from the behavioral data before kernel ridge regression. To prevent any information leak from the training data to test data, for each test fold, the nuisance regression was performed on the training folds and the regression coefficients were applied to the test fold.

### Comparison with alternative parcellation approaches

The above prediction procedure was repeated using parcellations estimated by YeoBackProject, Gordon2017 and Wang2015. The procedure could not be applied to the Yeo2011 approach because the group-level approach results in the same parcellations across subjects.

### Network size versus topography

In the previous sections, a test subject’s behavior was predicted based on the similarity between the individual-specific parcellations of the test subject and the training subjects, where similarity was measured based on how much corresponding networks spatially overlapped (i.e., Dice). Here, we investigated whether individual differences in network size could also predict behavior. This was achieved by defining the similarity between two parcellations to be the correlation between network sizes. More specifically, let *s* and *s*_*i*_ be 17 × 1 vectors with the *j*-th entries corresponding to the surface areas belonging to the *j*-th network of the test subject and *i*-th training subject respectively. Here, surface areas were measured in the subjects’ native space. The similarity between the individual-specific parcellations of the test subject and *i*-th training subject was set to be the correlation between *s* and *s*_*j*_. Therefore, test subject and *i*-th training subject might be similar due to corresponding networks having similar sizes, even though the networks might not significantly overlap. If the prediction accuracies were significantly higher for network overlap (compared with network size), then this would suggest that network topography (location), and not network size, was driving the behavioral prediction.

### Topography of task-relevant networks

We also investigated whether the topography of task-relevant networks might contribute to the prediction of various behavioral measures. For example, the frontoparietal control network is typically activated during working memory tasks. Would the topography of the individual-specific frontoparietal control network be more predictive of working memory performance than the topography of all networks?

To explore this question in a systematic fashion, we considered 13 cognitive measures highlighted in the HCP data dictionary (first 13 items in Table S1). For each cognitive measure, we searched for the “forward inference” map of the most relevant term in the NeuroSynth database (Yarkoni et al., 2011). The forward inference map quantified the likelihood that a particular brain voxel was activated in studies using that search term. The reverse inference maps were not considered because they turned out to be extremely sparse. Table S2 shows the search term utilized to obtain the forward inference map for each cognitive measure. There was no appropriate search term for two cognitive measures (processing speed and picture vocabulary), so they were excluded from further analyses. Each forward inference map was projected to fs_LR surface space (Buckner et al., 2011; Van Essen et al., 2012a) and compared with the group-level parcellation estimated from the HCP training set, in order to select the task-relevant networks (Table S2). When predicting a particular behavior, the similarity between the test subject and *i*-th training subject was set to be the dice coefficient averaged across task-relevant networks. For example, in the case of fluid intelligence, Control network A and Dorsal Attention network A overlapped the most with the forward inference map associated with “intelligence” (Table S2). When applying kernel regression to predict fluid intelligence, similarity between the test subject and *i*-th training subject was set to be the average dice coefficient for Control network A and Dorsal Attention network A.

### Code availability

Code for this work is freely available at the github repository maintained by the Computational Brain Imaging Group (https://github.com/ThomasYeoLab/CBIG). More specifically, the GSP and CoRR-HNU datasets were preprocessed using an in-house pipeline (https://github.com/ThomasYeoLab/CBIG/tree/master/stable_projects/preprocessing/CBIG_f MRI_Preproc2016). The group-level parcellation code (Yeo et al., 2011) are available here (https://github.com/ThomasYeoLab/CBIG/tree/master/stable_projects/brain_parcellation/Yeo 2011_fcMRI_clustering). Finally, the individual-specific parcellation code is also available (GITHUB_LINK_TO_BE_ADDED).

## Results

### Overview

The MS-HBM (Figure 1) was applied to three multi-session rs-fMRI datasets to ensure that the model can reliably estimate inter-subject and intra-subject variability despite significant acquisition differences across datasets. After confirming previous literature (Mueller et al., 2013; Laumann et al., 2015) that inter-subject and intra-subject RSFC variability were different across networks, we then established that the MS-HBM produced better parcellations than other approaches. Finally, we investigated whether the topography (location and spatial arrangement) and size of individual-specific cortical networks were behaviorally relevant.

### Sensory-motor networks exhibit lower inter-subject, but higher intra-subject, functional connectivity variability than association networks

Figure 2A shows the 17-network population-level parcellation estimated from the HCP training set. The 17 networks were divided into eight groups (Visual, Somatomotor, Auditory, Dorsal Attention, Salience/Ventral Attention, Control, Default and TempPar), which broadly corresponded to major networks discussed in the literature. The 17 networks were referred to as “Default A”, “Default B” and so on (Figure 2A).

**Figure 2.**
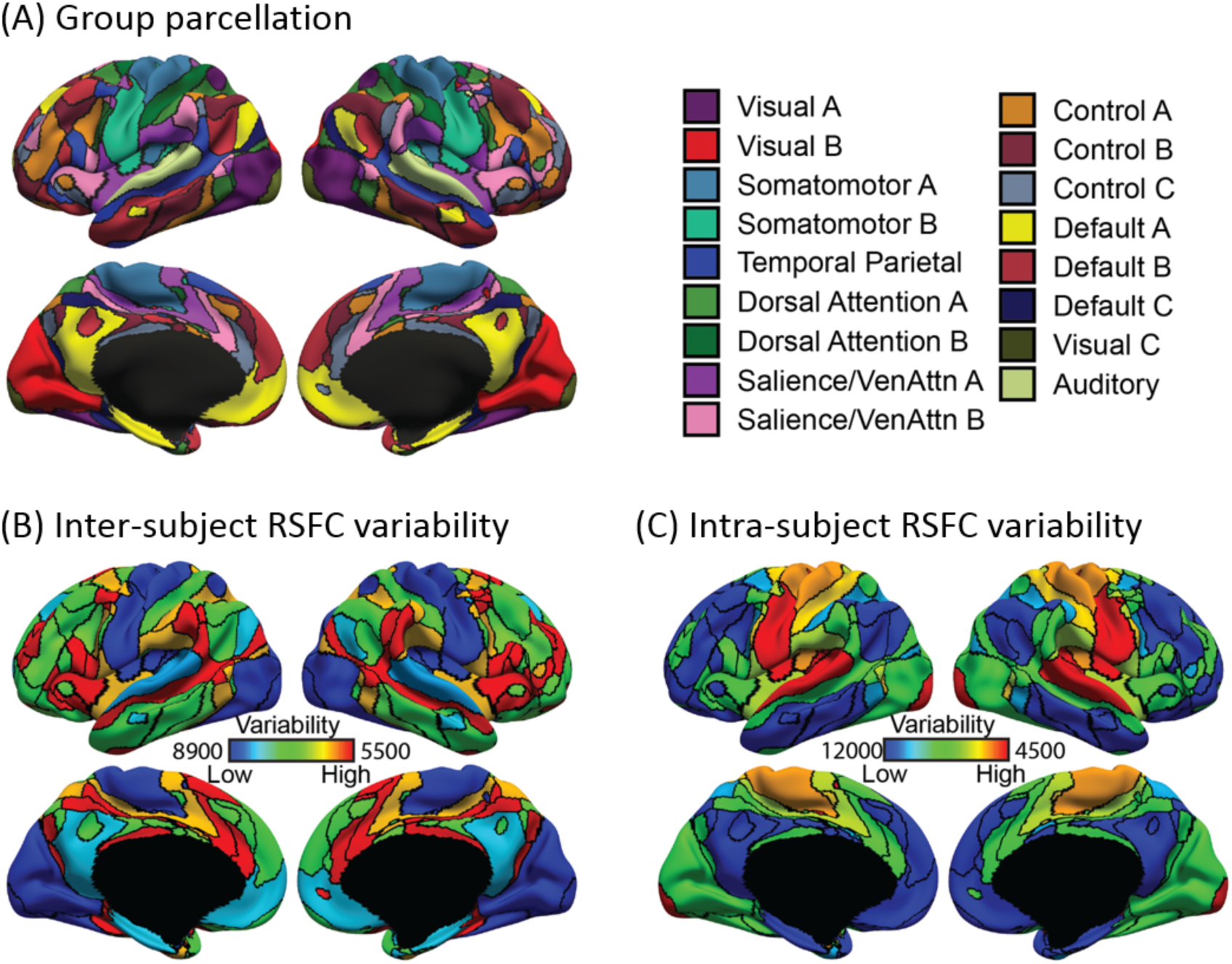
Sensory-motor networks exhibit lower inter-subject, but higher intra-subject, functional connectivity variability than association networks in the HCP training set. (A) 17-network group-level parcellation. (B) Inter-subject functional connectivity variability for different cortical networks. (C) Intra-subject functional connectivity variability for different cortical networks. Results were replicated in the GSP (Figure S1) and CoRR-HNU (Figure S2) datasets. Note that (B) and (C) correspond to the *∈*_*l*_ and *σ*_*l*_ parameters in Figure 1, where higher values indicate lower variability.

The HCP population-level parcellation was replicated in the GSP (Figure S1A) and CoRR-HNU (Figure S2A) datasets, although there were some interesting distinctions. For example, the Limbic (A and B) networks from the GSP population-level parcellation (Figure S1A) were absorbed into the Default (A and B) networks in the HCP population-level parcellation (Figure 2A). Instead, there were two additional networks in the HCP population-level parcellation: Visual C and Auditory networks. The Visual C network (Figure 2A) might correspond to the foveal representation within the primary visual cortex, while the Auditory network (Figure 2A) appeared to have split off from the Somatomotor B network in the GSP population-level parcellation (Figure S1A).

Increasing the number of subjects only resulted in minor changes in the group-level parcellations, so differences between population-level parcellations were probably due to acquisition differences, rather than sampling variability. For example, the higher resolution HCP data might allow the separation of the Auditory and Somatomotor B networks, which were in close spatial proximity.

Recall that the inter-subject functional connectivity variability *∈*_*l*_ was estimated for each network. Hence, *∈*_*l*_ could be visualized by coloring each corresponding population-level network from Figure 2A. Figure 2B shows *∈*_*l*_ estimated from the HCP training set. Consistent with previous literature (Laumann et al., 2015), sensory-motor networks exhibited lower inter-subject functional connectivity variability than association networks. More specifically, Somatomotor (A and B) and Visual (A and B) networks were the least variable, while Salience/Ventral Attention B network was the most variable. The results were largely consistent in the GSP (Figure S1B) and CoRR-HNU (Figure S2B) datasets, although there were some notable differences. For example, the Somatomotor B network exhibited low variability in both the GSP and HCP datasets, but intermediate variability in the CoRR-HNU dataset.

Similar to *∈*_*l*_, the intra-subject functional connectivity variability *σ*_*l*_ was estimated for each network. Hence, *σ*_*l*_ could be visualized by coloring each corresponding population-level network from Figure 2A. Figure 2C shows *σ*_*l*_ estimated from the HCP training set. Consistent with previous literature (Laumann et al., 2015), association networks exhibited lower intra-subject functional connectivity variability than sensory-motor networks. More specifically, Default (A and B) networks were the least variable, while Somatomotor (A and B), Auditory and Visual C networks were the most variable. The results were largely consistent in the GSP (Figure S1C) and CoRR-HNU (Figure S2C) datasets, although there were some interesting differences. Of particular note is that Visual B Network exhibited high intra-subject functional connectivity variability in the GSP dataset, but low or intermediate functional connectivity variability in the CoRR-HNU and HCP datasets. This difference might be due to subjects instructed to fixate on a cross in the CoRR-HNU and HCP datasets, while subjects were told to keep their eyes open (with no fixation cross) in the GSP dataset.

It is worth noting that in the MS-HBM (Figure 1), higher values of *∈*_*l*_ and *σ*_*l*_ indicate lower variability. The values in Figure 2C are much larger than Figure 2B, suggesting that intra-subject functional connectivity variability is much lower than inter-subject functional connectivity variability. These results were replicated in the GSP (Figure S1) and CoRR-HNU (Figure S2) datasets.

### Sensory-motor networks are less spatially variable than association networks across subjects

The MS-HBM model differentiated between inter-subject RSFC and network spatial variability. Like inter-subject functional connectivity variability, the sensory-motor networks were found to be less spatially variable than association networks across subjects. For example, Figure S3 shows the inter-subject spatial variability maps of four representative networks from the HCP training set. Yellow color at a spatial location indicates that across subjects, there is a high probability of the network appearing at that spatial location, suggesting low inter-subject spatial variability. The Somatomotor A network and Visual B network showed higher probabilities (more yellow color) than the Dorsal Attention networks, suggesting that Somatomotor A network and Visual B network exhibited lower inter-subject spatial variability than Dorsal Attention networks. These results were consistent in the GSP (Figure S4) and CoRR-HNU (Figure S5) datasets.

### Individual-specific networks generated by MS-HBM exhibit higher resting-state homogeneity than other approaches

Individual-specific parcellations were estimated using one rs-fMRI session from the CoRR-HNU dataset and HCP test set. The resting-state homogeneity of the parcellations were evaluated in the leave-out sessions (Figure 3A). Across both CoRR-HNU and HCP datasets, the group-level parcellation (Yeo2011) achieved the worst resting-state homogeneity, while MS-HBM performed the best. In the CoRR-HNU dataset, compared with Yeo2011, YeoBackProject, Gordon2017 and Wang2015, the MS-HBM achieved an improvement of 16.6% (Cohen’s d = 4.6), 5.3% (Cohen’s d = 3.5), 6.9% (Cohen’s d = 3.4) and 4.2% (Cohen’s d = 3.4) respectively. In the HCP dataset, compared with Yeo2011, YeoBackProject, Gordon2017 and Wang2015, the MS-HBM achieved an improvement of 9.8% (Cohen’s d = 3.2), 9.5% (Cohen’s d = 3.0), 5.7% (Cohen’s d = 2.1) and 4.4% (Cohen’s d = 3.1) respectively.

**Figure 3.**
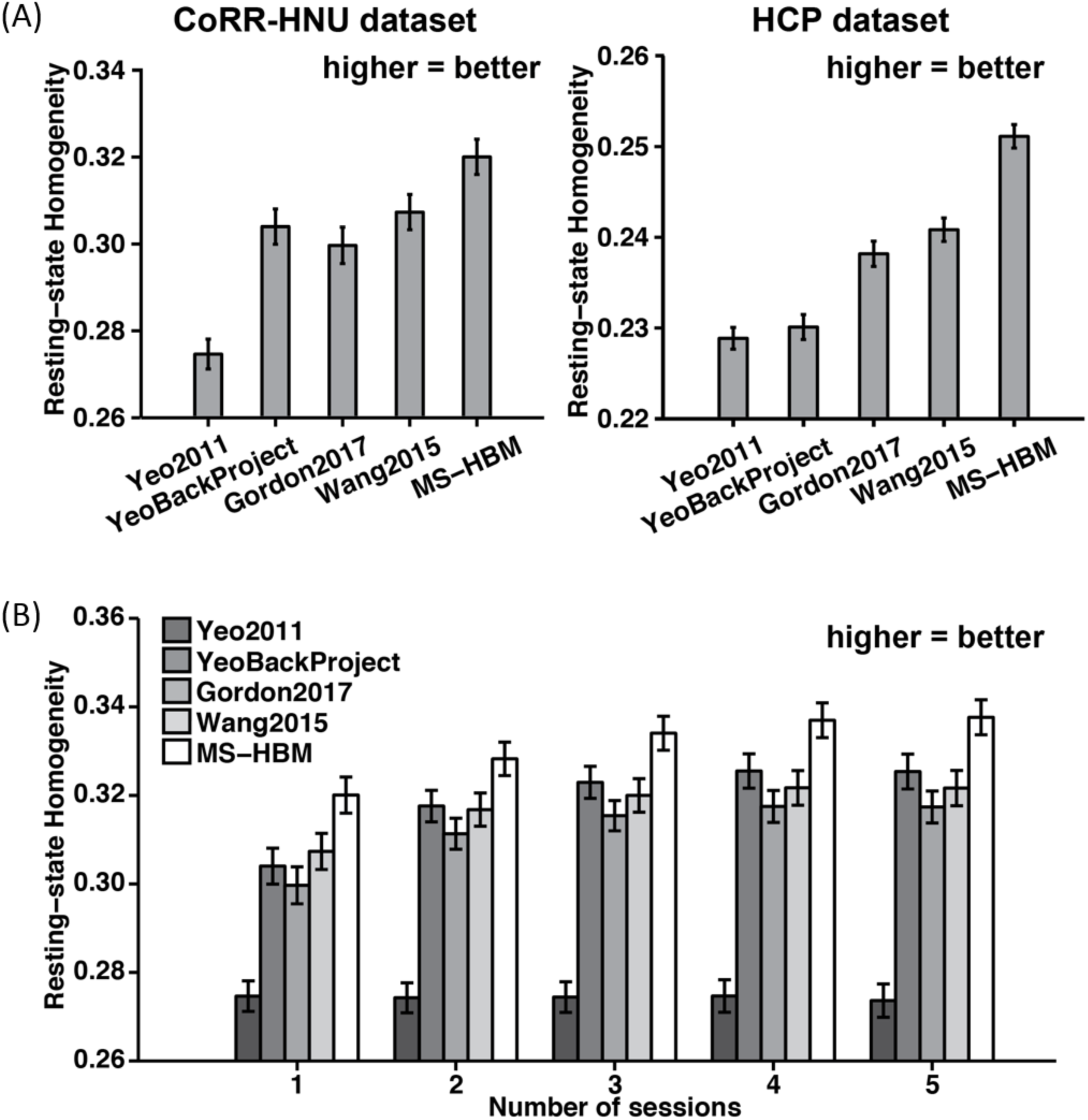
Resting-state homogeneity in the CoRR-HNU and GSP dataset. (A) 17-network individual-specific parcellations were estimated using one rs-fMRI session and resting-state homogeneity were computed on the remaining sessions for each subject from the CoRR-HNU and HCP dataset. (B) 17-network individual-specific parcellations were estimated using different number of rs-fMRI sessions and resting-state homogeneity were computed on the remaining sessions for each subject from the CoRR-HNU dataset. Error bars correspond to standard errors. Using just one single fMRI sessions (10 min), the MS-HBM algorithm was able to match the homogeneity achieved with Wang2015 and Gordon2017 using five fMRI sessions (50 min).

Individual-specific parcellations were estimated with increasing number of rs-fMRI sessions using the CoRR-HNU dataset. The resting-state homogeneity of the parcellations were evaluated in the leave-out sessions (Figure 3B). Not surprisingly, performance of the Yeo2011 group-level parcellation remained constant regardless of the amount of data. The remaining approaches (YeoBackProject, Gordon2017, Wang2015 and MS-HBM) exhibited higher homogeneity with increased number of sessions. Critically, the improvement of our model over the other approaches grew with the inclusion of additional fMRI sessions. For example, as the number of sessions was increased from two to three to four to five, the MS-HBM achieved improvement of 5.4%, 5.9%, 6.1% and 6.4% respectively over Gordon2017. Interestingly, the improvement of our approach over Gordon2017 was largest when only one rs-fMRI session was utilized (6.9%). On the other hand, the MS-HBM achieved improvement of 3.6%, 4.4%, 4.8% and 5.0% respectively over Wang2015. Furthermore, using just one fMRI sessions (10 min), the MS-HBM was able to match the homogeneity achieved with the Wang2015 and Gordon2017 approaches using five fMRI sessions (50 min).

### Individual-specific networks generated by the MS-HBM exhibit lower task functional inhomogeneity than other approaches

Individual-specific parcellations were estimated using one rs-fMRI run (15 min) from the HCP test set. Figure S6 shows the task inhomogeneity of the different approaches. Compared with Yeo2011, the MS-HBM achieved a small improvement of 0.63% (Cohen’s d = 0.12, 0.09, 0.66, 1.0, 0.9, 1.1, 0.46 for social, motor, gambling, relational, language, working memory and emotion respectively). Compared with YeoBackProject, Gordon2017 and Wang2015, MS-HBM achieved improvements of 2.0% (Cohen’s d > 1.3 for all domains), 1.04% (Cohen’s d > 0.99 for all domains) and 0.7% (Cohen’s d > 0.79 for all domains) respectively. Interestingly, the Yeo2011 group-level parcellation performed as well as (or even better than) YeoBackProject and Gordon2017.

### Individual-specific MS-HBM parcellations exhibit high intra-subject reproducibility and low inter-subject similarity

To assess intra-subject reproducibility and inter-subject similarity, our model (Figure 1) was tuned on the HCP training and validation sets, and then applied to the HCP test set. Individual-specific parcellations were generated by using the first two runs and last two runs separately for each subject. Figures 4 and S7 show the parcellations of four representative subjects. The 17 networks were present in all individual-specific parcellations. However, network shapes, sizes and topologies were varied across subjects, consistent with previous studies of individual-specific brain networks (Harrison et al., 2015; Laumann et al., 2015; Wang et al., 2015; Gordon et al., 2017c; Braga and Buckner, 2017).

**Figure 4.**
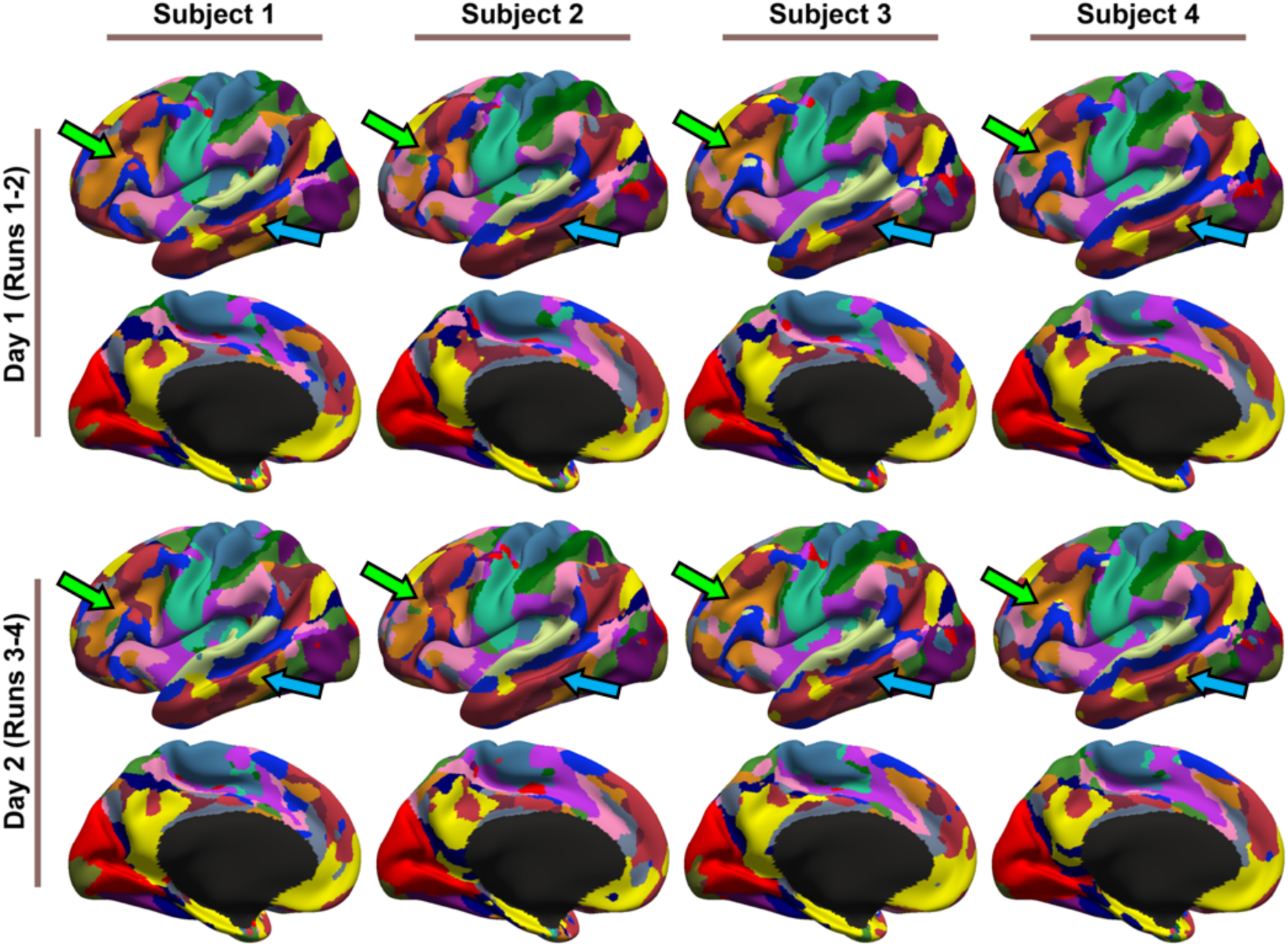
17-network parcellations were estimated using runs 1-2 and runs 3-4 separately for each subject from the HCP test set. Parcellations of four representative subjects are shown here. Blue and green arrows indicate individual-specific parcellation features. Right hemisphere parcellations are shown in Figure S7.

For example, the Default A (yellow) network exhibited a lateral temporal component in certain subjects (blue arrows in Figure 4), but was missing in other subjects. As another example, the two lateral prefrontal components of the Control A (orange) network (Figure 2A) were fused into a single component in certain subjects (green arrows in Figure 4). These features were mostly replicated across sessions. Examples from the CoRR-HNU dataset are shown in Figures S8 and S9.

Figure S10A shows the across-subject spatial similarity (Dice coefficient) of individual-specific parcellations. A higher value (hot color) indicates greater inter-subject agreement. Figure S10B shows the within-subject reproducibility (Dice coefficient) of individual-specific parcellations. A higher value (hot color) indicates greater inter-session agreement within subjects. Further quantification is shown in Figure S10C, where the Dice coefficients were averaged across sub-networks.

Across all networks, intra-subject reproducibility was greater than inter-subject similarity. Compared with association networks, the Somatomotor (A and B) networks and Visual (A and B) networks were more spatially similar across subjects, but also exhibited greater within subject inter-session reproducibility. Overall, the MS-HBM parcellation model achieved 77.9% intra-subject reproducibility and 65.4% inter-subject similarity.

The results were similar in the CoRR-HNU dataset (Figure S11), although intra-subject reproducibility was higher (81.6%) and inter-subject similarity was lower (59.4%). The improvement might be the result of longer scan duration in the CoRR-HNU dataset (50 min versus 30 min).

### Individual differences in network topography can predict cognition, personality and emotion

While it is well-known that individual-specific networks exhibit unique topographic features that are replicable across sessions (Laumann et al., 2015; Gordon et al., 2017c; Braga and Buckner, 2017), their behavioral relevance remains unclear. Here, we found that individual-specific network topography was able to predict the 58 behavioral measures with varying degree of accuracies.

Figure 5 shows the prediction accuracy for 13 cognitive measures highlighted in the HCP data dictionary. Average prediction accuracy was r = 0.1321 ± 0.0053. Reading (pronunciation) and delay discounting could be predicted relatively well with accuracies of r = 0.2918 ± 0.0141 (mean ± std) and r = 0.2398 ± 0.0166. The prediction accuracies for the remaining cognitive, emotion and personality measures are found in Figures S12 and S13. In the case of the NEO-5 personality scores (Figure S12), average predication accuracy was r = 0.0955 ± 0.0085. Interestingly, the prediction of emotional recognition (Figure S13) was poor with an average prediction accuracy of r = −0.0445 ± 0.0101. The remaining emotional measures (all items in Figure S13 except for emotional recognition) could be predicted with an average accuracy of r = 0.1038 ± 0.0070.

**Figure 5.**
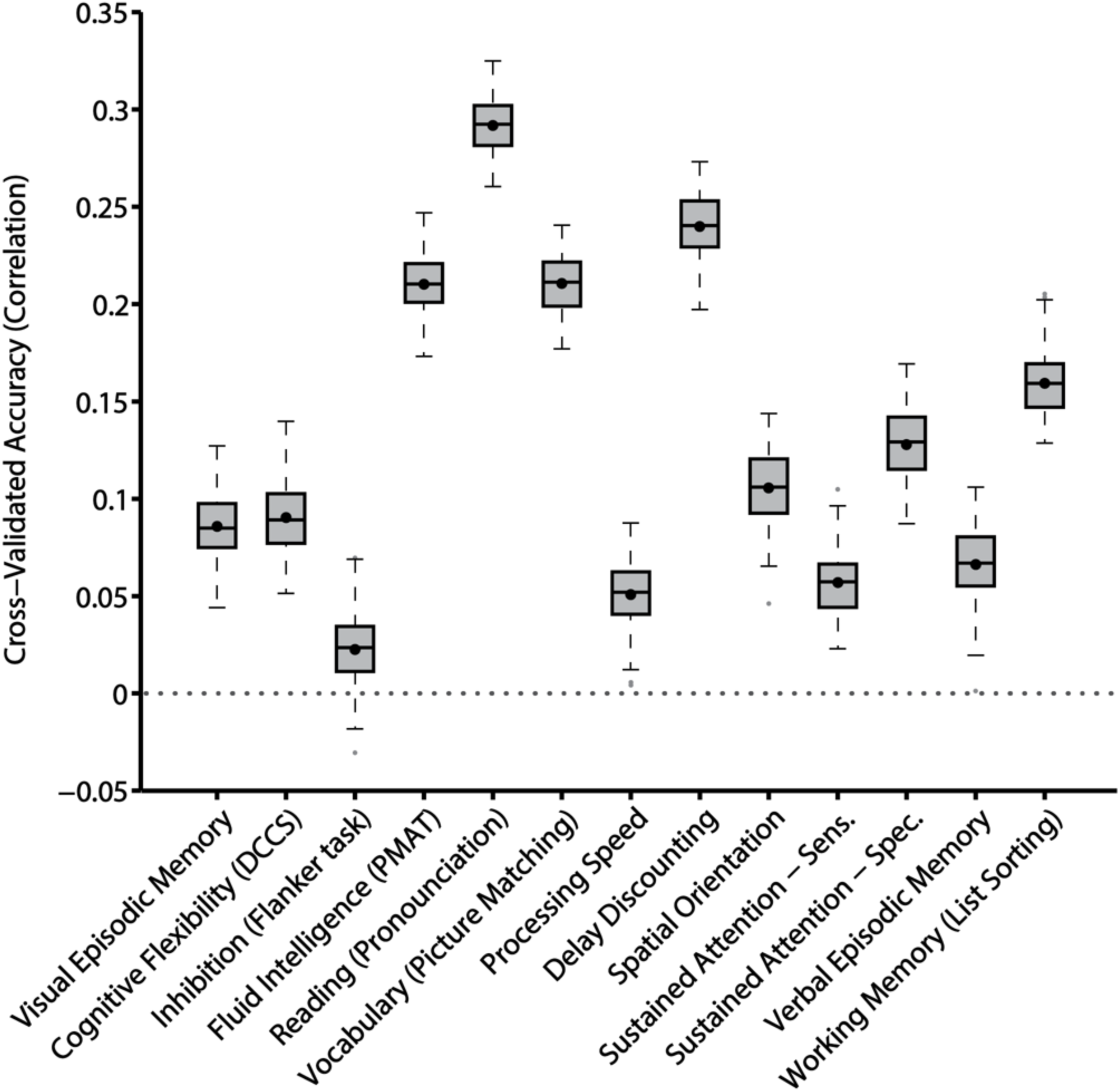
Prediction accuracy of 13 cognitive measures based on inter-subject differences in the spatial arrangement of cortical networks. Boxplots utilized default Matlab parameters, i.e., box shows median and inter-quartile range (IQR). Whiskers indicate 1.5 IQR. Dot indicates mean. Average prediction accuracy was r = 0.1321 ± 0.0053 (mean ± std) for the 13 measures. Other behavioral measures are found in Figures S12 and S13.

In the case of the minimally correlated set of 5 behavioral measures, the average prediction accuracy was r = 0.1327 ± 0.0065. Across all 58 behavioral measures, an average prediction accuracy of r = 0.0803 ± 0.0032 (mean ± std) was obtained. While the accuracy might seem modest, they were comparable to (if not better than) other studies using functional connectivity *strength* for behavioral prediction (HCP MegaTrawl; https://db.humanconnectome.org/megatrawl/; Noble et al., 2017; Dubois et al., biorxiv). For example, of the 58 behavioral measures, 49 of them were also utilized in the HCP MegaTrawl. For the 300-dimensional group-ICA results, HCP MegaTrawl achieved an average accuracy of r = 0.0592 (original data space), while kernel regression yielded an average accuracy of r = 0.0874 ± 0.0036.

### MS-HBM network topography is more predictive of behavioral measures compared with other parcellation approaches

Table S3 summarizes the average prediction accuracies for different sets of behavioral measures (58 behavioral measures, 13 cognitive measures, NEO-5 personality measures, emotion recognition measures, emotional measures and minimally correlated set of 5 behaviors). Overall, MS-HBM network topography achieved better prediction accuracies compared with other approaches.

Figure 6 shows the average prediction accuracies of the minimally correlated behavioral set across different parcellation approaches. Compared with YeoBackProject, Gordon2017 and Wang2015, MS-HBM achieved percentage improvements of 29%, 61% and 28% respectively. Furthermore, MS-HBM achieved the best prediction accuracy for the minimally correlated behavioral set for each of the 100 20-fold cross-validations (Table S3).

**Figure 6.**
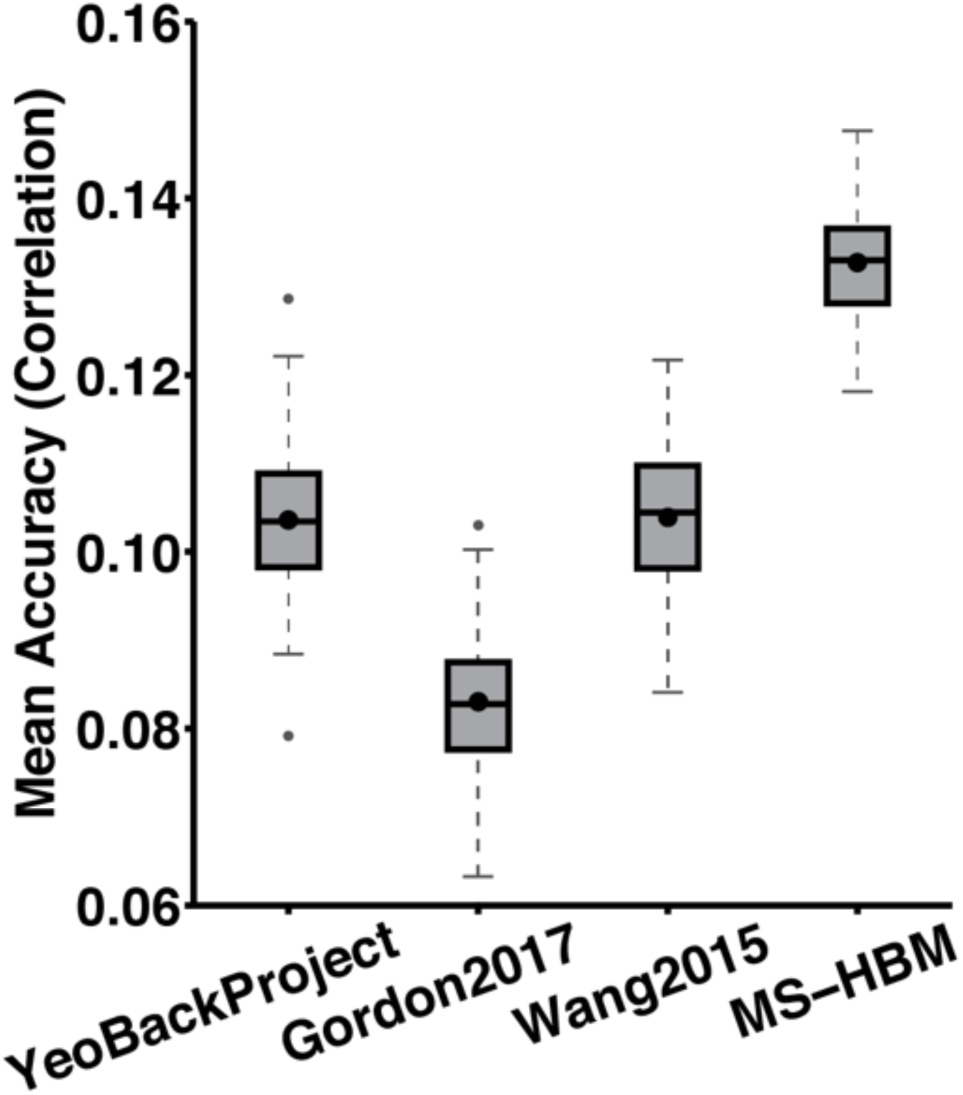
Average prediction accuracies of the minimally correlated set of 5 behavioral measures across different parcellation approaches. Prediction was based on individual-specific network topography. Boxplots utilized default Matlab parameters, i.e., box shows median and inter-quartile range (IQR). Whiskers indicate 1.5 IQR. Dot indicates mean. MS-HBM, YeoBackProject, Gordon2017, Wang2015 achieved average prediction accuracies of r = 0.1327 ± 0.0065 (mean ± std), r = 0.1036 ± 0.0080, r = 0.0830 ± 0.0080 and r = 0.1039 ± 0.0080.

There were two exceptions. First, MS-HBM performed the worst in the emotional recognition measures. Second, network topography estimated with YeoBackProject was better than MS-HBM in predicting NEO-5 personality measures.

### Controlling for motion-related imaging artifacts

Given that certain behavioral measures are known to correlate with motion (Siegel et al., 2016), we tested if network topography could predict FD and DVARS (without regressing any nuisance covariates). Network topography estimated by all four parcellation approaches (MS-HBM, YeoBackProject, Gordon2017, Wang2015) could predict FD and DVARS quite well (Table S4), suggesting that individual-specific network topography might encode information about motion-related imaging artifacts or motion-related traits or both (Zeng et al., 2014; Siegel et al., 2017).

If individual-specific MS-HBM parcellations were corrupted by motion-related imaging artifacts, then state-related FD and DVARS would have a significant effect on intra-subject parcellation reproducibility. However, further analyses involving participants with very different FD and DVARS across the two scan days indicated that state-related FD and DVARS had little effect on intra-subject reproducibility (Figures S14 and S15). Together with the successful prediction of FD and DVARS (Table S4), this suggests that individual-specific network topography (estimated by MS-HBM) likely encoded a significant amount of information about motion-related traits. Nevertheless, we cannot completely rule out the possibility that aspects of network topography might be corrupted by motion-related imaging artifacts.

To address the concern that the behavioral prediction results in the previous sections might simply be due to the regression algorithm encoding motion-related imaging artifacts that were correlated with the behavioral measure of interest, we considered the five behavioral measures most correlated with FD: endurance, cognitive flexibility (DCCS), vocabulary (picture matching) and reading (pronunciation). Figure S16A shows that the average correlation between the prediction of the five behavioral measures and FD was close to zero, and was significantly lower than the average prediction accuracy of the five behavioral measures. In fact, across the 58 behavioral measures, higher correlation with FD was associated with worse behavioral prediction accuracy (r = −0.22). Similar results were obtained for DVARS (Figure S16B). Together, this suggests that the behavioral prediction results in the previous section could not be simply explained by motion-related imaging artifacts.

### Network size versus network topography

When utilizing individual differences in network size to predict behavior, the average accuracy of the minimally correlated set of 5 behavioral measures was r = 0.0865 ± 0.0105 (mean ± std; Figure 7). This was worse than using network topography (Figure 7). Average prediction accuracy for the 13 cognitive measures was r = 0.0483 ± 0.0073. In the case of the NEO-5 personality scores, average prediction accuracy was r = 0.0751 ± 0.0100, while the average prediction accuracy of the emotional measures was r = 0.0782 ± 0.0086. The average prediction accuracy of 58 behavioral measures was r = 0.0412 ± 0.0047. Thus, individual differences in network size (at least at the resolution of large-scale networks) could not account for the ability of network topography to predict behavior.

**Figure 7.**
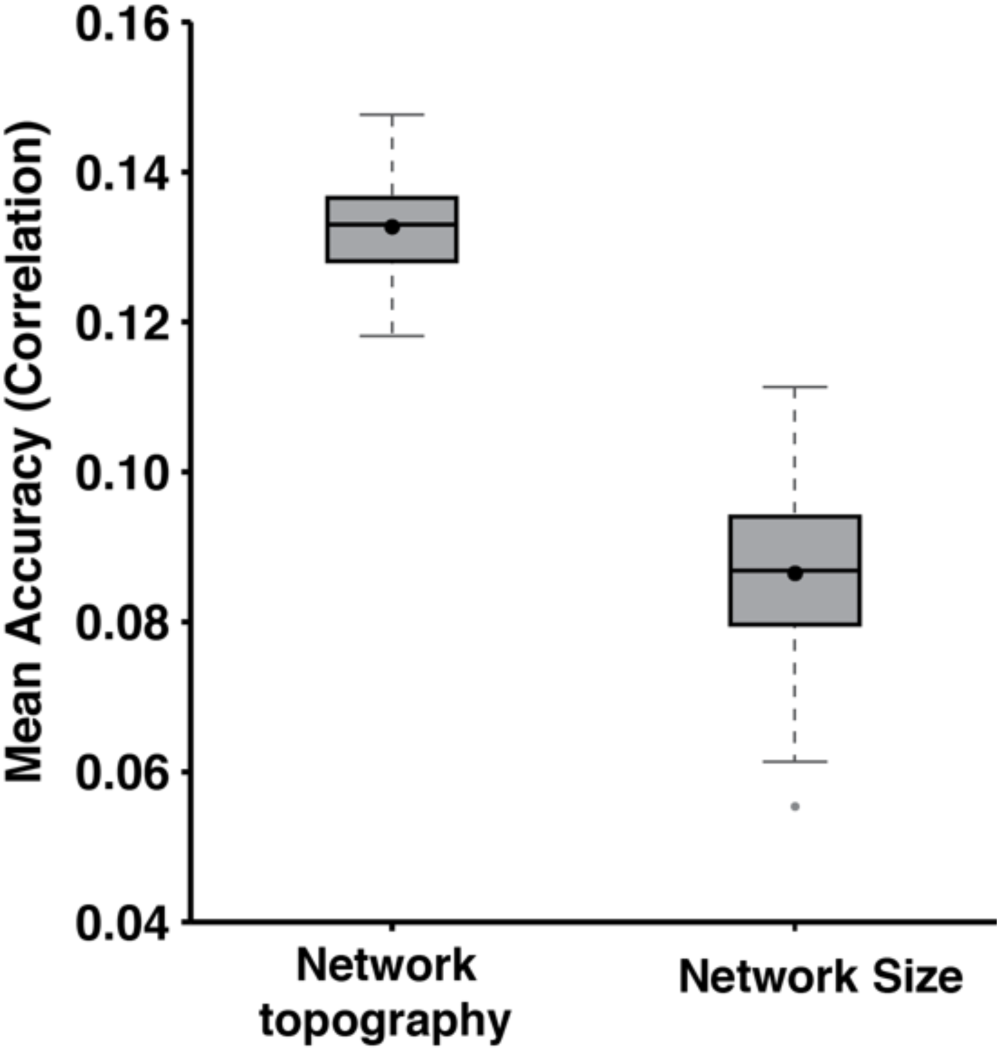
Average prediction accuracies of the minimally correlated set of 5 behavioral measures based on inter-subject differences in network topography or network size. Boxplots utilized default Matlab parameters, i.e., box shows median and inter-quartile range (IQR). Whiskers indicate 1.5 IQR. Dot indicates mean. Average prediction accuracy based on network topography was r = 0.1327 ± 0.0065 (mean ± std). Average prediction accuracy based on network size was r = 0.0865 ± 0.0105.

### Topography of task-relevant networks

When utilizing the topography of only task-relevant networks (rather than all networks) to predict behavior, the average accuracy across 11 cognitive measures (see Methods) was r = 0.1129 ± 0.0062 (mean ± std; Figure 8). This was worse than utilizing the topography of all networks r = 0.1324 ± 0.0056.

**Figure 8.**
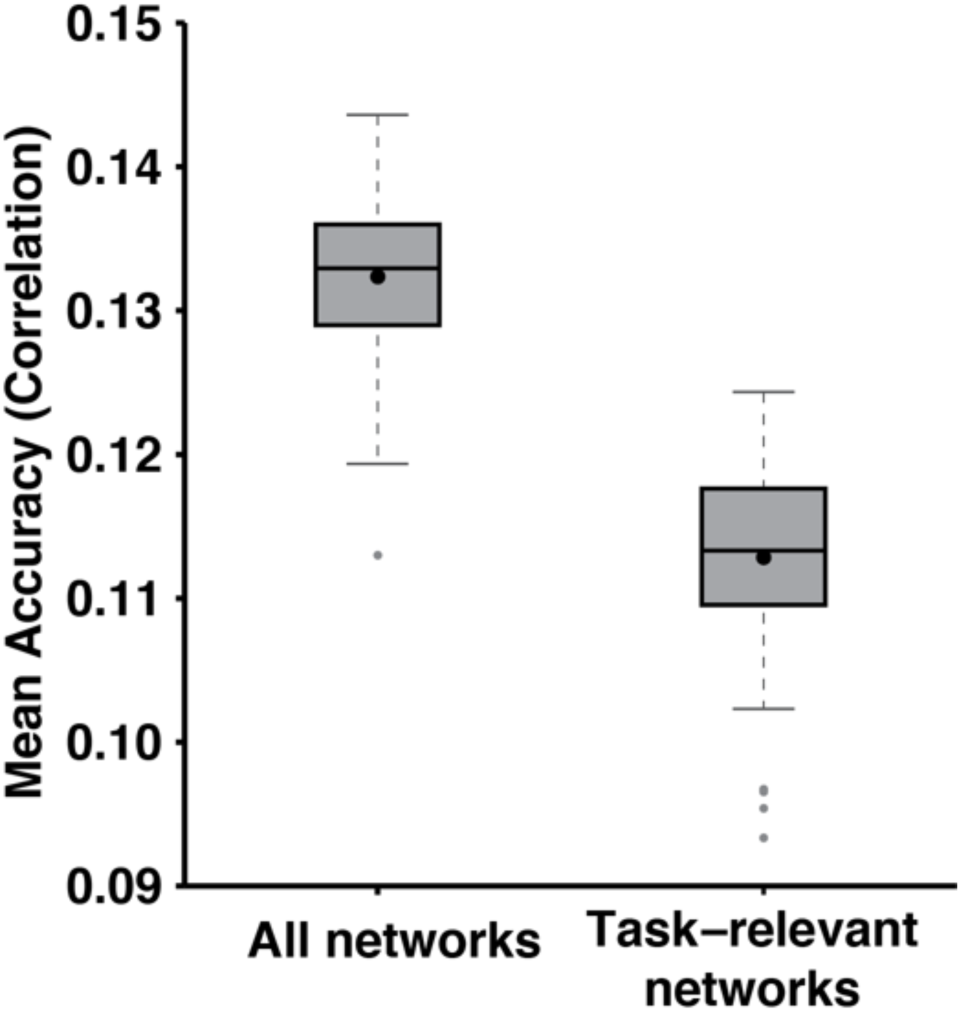
Average prediction accuracy across 11 cognitive measures based on topography of all networks or only task-relevant networks. Boxplots utilized default Matlab parameters, i.e., box shows median and inter-quartile range (IQR). Whiskers indicate 1.5 IQR. Dot indicates mean. Average prediction accuracy based on all networks was r = 0.1324 ± 0.0056. Average prediction accuracy based only on task-relevant networks was r = 0.1129 ± 0.0062. Prediction accuracy of each cognitive measure is found in Figure S17.

## Discussion

Using a novel multi-session hierarchical Bayesian model (MS-HBM), we demonstrate that individually-specific cortical network topography could predict multiple behavioral phenotypes across cognition, personality and emotion. The prediction accuracies could not be accounted for by individual differences in network size. The MS-HBM allowed the joint estimation of inter-subject variability, intra-subject variability and individual-specific cortical networks within the same statistical framework. The resulting MS-HBM individual-specific parcellations were more homogeneous than parcellations derived with four alternative approaches during both resting and task states. These analyses suggest that the *spatial topography* of individuals-specific brain networks might serve as a generalizable fingerprint of human behavior, similar to the preponderance of studies utilizing functional connectivity *strength*.

### Neurobiological interpretation of MS-HBM networks

By assuming individual-specific parcellations to be the same across sessions, the MS-HBM essentially treats inter-session differences as noise. The implication is that individual-specific MS-HBM parcellations seek to capture stable, trait-like network organization in individuals. However, it is well-known that certain factors (e.g., caffeine intake, sleepiness, attention) result in different brain states and thus functional network organization (Tagliazucchi and Laufs, 2014; Laumann et al., 2015; Poldrack et al., 2015; Yeo et al., 2015b; Wang et al., 2016; Shine et al., 2016). Moreover, in longitudinal studies of certain populations, e.g., Alzheimer’s Disease dementia, the goal is to detect longitudinal changes across consecutive sessions (Misra et al., 2009; Raj et al., 2015; Risacher et al., 2010; Zhang et al., 2016; Lindemer et al., 2017). To capture transient session-specific or longitudinal changes in brain network organization, the model could be modified to allow for spatial differences in individual-specific parcellations across sessions.

The human cerebral cortex is hierarchically organized (Churchland and Sejnowski 1988) from molecules (1Å) to synapses (1µm) to neurons (100µm) to areas (1 cm) and systems (10cm). Even at the relatively coarse spatial resolution of MRI, hierarchical organization can be observed. Here, we focused on parcellating the cerebral cortex into less than twenty spatially distributed networks. Each spatial (e.g., parietal) component of a network likely spans multiple cytoarchitectonically, functionally and connectionally distinct cortical areas (Kaas 1987; Felleman and Van Essen 1991; Amunts and Zilles 2015; Eickhoff et al., 2018). We are working on extending the MS-HBM to estimate a finer division of the cerebral cortex that approximate these finer regions, i.e., individual-specific areal-level parcellation (Laumann et al., 2015; Glasser et al., 2016; Gordon2017c).

A biologically plausible individual-specific parcellation should simultaneously capture (genuinely) shared features across individuals, while preserving individual-specific trait-level variation. All individual-specific parcellation approaches ultimately make explicit or implicit assumptions about what features are shared or varied across individuals. For example, in the MS-HBM, an important parameter is the number of networks. While there is no penalty for a participant to exhibit a smaller number of networks, in practice, we do find all networks in all subjects. One could potentially estimate different number of networks in each participant by parcellating each participant independently (Laumann et al., 2015; Gordon et al, 2017c). However, given the significantly less data available in each subject (compared with Gordon et al., 2017c), any network differences between subjects (e.g., less or more networks) could simply be due to convergence to different local optima as a result of noise.

It is also worth noting that although Gordon and colleagues did not explicitly constrain the number of networks to be the same in each participant (Laumann et al., 2015; Gordon et al., 2017c), certain parameters that could dramatically affect the estimated networks (e.g., range and sampling intervals of edge density thresholds) were assumed to be the same across subjects. This is a key challenge for all individual-specific parcellation approaches: it is not possible to set completely different parameters for every single individual, since it is unclear how those might be set, and we would also like the parcellations to be comparable across individuals. Yet, setting the same parameters for every individual might not be biologically plausible.

### Spatial configuration of individual-specific cortical networks is behaviorally meaningful

Recent work has suggested that individual-specific functional networks exhibit unique topological features not observed in group-level networks (Harrison et al., 2015; Laumann et al., 2015; Wang et al., 2015; Glasser et al., 2016; Langs et al., 2016; Braga & Buckner, 2017; Gordon et al., 2017a; 2017b; 2017c). This was also clearly the case with individual-specific MS-HBM parcellations (Figures 4, S8). While we have pointed out two examples (Default A and Control A networks), it was also obvious that many other individual-specific parcellation features were replicable across sessions.

A major unanswered question in the literature is whether individual differences in cortical parcellations are behaviorally meaningful. A recent paper (Salehi et al., 2018) found that individual-specific brain parcellations could be used to predict individuals’ sex, while another paper (Bijsterbosch et al., 2018) has associated individual-specific network topography with a single positive-negative axis of behavior. In contrast, we utilized kernel regression to investigate whether individual-specific network topography and size could be used to predict 58 behavioral measures. The kernel regression framework utilized in this work tested the possibility that subjects with more similar parcellations exhibited similar behavior. Successful prediction (Figures 5, S12, S13) suggests that inter-subject variation in the spatial configuration of cortical networks is strongly related to inter-subject variation in behavior.

The topography of task-relevant networks was also effective for predicting cognitive performance, although interestingly, the prediction accuracies were no better than when using the topography of all networks. It might seem somewhat counterintuitive that the inclusion of task-irrelevant networks (as measured by meta-analysis of task activation) did not dilute prediction accuracy. However, previous studies have suggested that regions not activated by a task might nevertheless exhibit connectivity changes during task performance (Cole et al., 2014; Krienen et al., 2014). Therefore, some of the task-irrelevant networks might potentially be involved in executing a task, despite not being activated through traditional task contrast or subtraction analyses. Alternatively, non-task regions might provide predictive information about traits associated with task performance without direct involvement in the task mechanisms.

Finally, a vast body of literature has utilized inter-region functional connectivity *strength* to predict behavior (Hampson et al., 2006; Finn et al., 2015; Rosenberg et al., 2016; Smith et al., 2015; Yeo et al., 2015b), in some cases implicitly, e.g., by binarizing a functional connectivity matrix and then utilizing the resulting graph metrics for behavioral association (van den Heuvel et al., 2009). It is worth noting that while spatial topography and functional connectivity strength might seem conceptually orthogonal (Bijsterbosch et al., 2018), they are not trivial to separate in practice. After all, most (if not all) individual-specific parcellation (or network estimation) approaches rely to some extent on functional connectivity strength. Therefore, although it seems plausible that network topography estimated by MS-HBM contains neural information complementary to that of functional connectivity strength, some aspects of network topography might still reflect functional connectivity strength.

Consequently, we believe that it will be a worthwhile endeavor to investigate whether individual differences in network topography and individual differences in functional connectivity strength could be combined to further improve behavioral prediction. We note that because only 17 networks were estimated in this work, we could only obtain 17 × 17 connectivity matrices, which were not optimal for behavioral prediction (e.g., Finn et al., 2015). We are currently developing an individual-specific areal-level parcellation approach, which would allow us to more effectively explore the possibility of combining network topography and connectivity strength.

### Network size versus network topography

It is well-known that the amount of brain “real estate” devoted to a cognitive function often predicts functional importance or capability. For example, larger hippocampal size has also been associated with better memory (Erickson et al., 2011). As another example, the acquisition of reading skills coincides with a change in the sizes of functionally defined visual areas (Dehaene et al., 2010).

Consistent with the vast literature, our results suggest that individual differences in network size can predict human behavior (Figure 7). However, the prediction accuracy is significantly weaker than network topography. This is surprising given that there is little evidence in the literature that the *location* of functional areas or networks might be functionally important.

One important caveat is that most previous studies associating anatomical size with functional importance are focused on smaller brain structures (Erickson et al., 2011; Holmes et al., 2016; Sabuncu et al., 2016), rather than large-scale networks. Therefore, it is possible that prediction accuracies could significantly improve with individual-specific areal-level parcellations.

### Individual-specific MS-HBM parcellations are more homogeneous than other approaches during resting and task states

If an individual-specific parcellation is capturing the organization of a subject’s cerebral cortex, then to the first order approximation, one might expect regions within the same network to have similar resting-state time series, as well as similar activation amplitude for any given task contrast (Gordon et al., 2017c; Schaefer et al., in press). Across the CoRR-HNU and HCP datasets, individual-specific MS-HBM parcellations exhibited greater resting-state functional connectivity homogeneity than parcellations from four other approaches (Figure 3), suggesting that MS-HBM parcellations better capture the “intrinsic” organization of individuals’ cerebral cortex. Importantly, group-level priors (e.g., inter-subject and intra-subject variability) estimated from the GSP dataset could improve the estimation of individual-specific parcellations in the CoRR-HNU dataset (Figure 3A). This is important because estimates of inter-subject and intra-subject functional connectivity variability were similar, but not the same across datasets (Figures 2, S1, S2). Therefore, our results suggest that the MS-HBM approach can be used to parcellate individuals from new datasets (using the same preprocessing pipeline), without having to re-estimate the group-level priors.

In the HCP dataset, individual-specific MS-HBM parcellations also exhibited greater task functional homogeneity than parcellations from four other approaches (Figure S6), suggesting that MS-HBM parcellations better capture the “extrinsic” organization of individuals’ cerebral cortex. Given the strong link between task fMRI and rs-fMRI (Smith et al., 2009; Mennes et al., 2010; Cole et al., 2014; Krienen et al., 2014; Bertolero et al., 2015; Yeo et al., 2015a; Tavor et al., 2016; Gordon et al., 2017c), this might not seem surprising. However, it is worth pointing out that the group-level parcellation performed as well as, if not better than two of the individual-specific parcellation approaches (Figure S6). Furthermore, the MS-HBM only demonstrated small improvements over the group-level parcellation in five of seven functional domains, while there was no statistical difference in the two remaining functional domains. One possibility is that the large-scale networks were too coarse to capture the finer details of task activation, consistent with our previous observation that the NeuroSynth forward inference maps do not cleanly match subtle features of the group-level large-scale networks. For example, the right-hand motor task preferentially activates the hand region of the left somatomotor cortex. However, Somatomotor A network covers the hand, foot and body regions of bilateral somatomotor cortex. As such, even if individual-specific Somatomotor A network were highly accurate, the resulting task inhomogeneity would still be relatively high.

### MS-HBM approach works well with single-session rs-fMRI data

Increasing the scan duration of rs-fMRI can improve the reliability of functional connectivity measures (Van Dijk et al., 2010; Xu et al., 2016). While earlier studies have suggested that 5 to 12 minutes of resting-state scan might be sufficient to provide reliable measurements (Van Dijk et al., 2010; Birn et al., 2013), more recent studies have suggested the need for 25 to 30 minutes of data (Anderson et al., 2011; Laumann et al., 2015; Gordon et al., 2017c). However, it is important to note that the amount of data necessary for reliable measurements depends on the functional connectivity measures being computed (Gordon et al., 2017c), as well as the methods employed.

Consistent with previous studies, our experiments showed that the quality of individual-specific parcellations improved with more rs-fMRI data, although the improvements plateaued after around 30 to 40 minutes of data (Figure 3B). Importantly, even though the MS-HBM was developed for multi-session rs-fMRI, the algorithm performed well even with single-session data. For example, the individual-specific MS-HBM parcellations estimated with one rs-fMRI session (10 minutes) exhibited comparable resting-state connectional homogeneity with parcellations estimated by two prominent approaches using five times the amount of data (Wang et al., 2015; Gordon et al., 2017a, 2017b). This improvement was made possible by exploiting prior information (e.g., group-level connectivity profiles, inter-subject RSFC variability and intra-subject RSFC variability) learned from multi-session rs-fMRI data. We expect high quality individual-specific parcellations to require significantly longer scan time if no such prior information was used.

### Methodological considerations

Consistent with recent studies (Mueller et al., 2013; Chen et al., 2015; Laumann et al., 2015), we found that association networks exhibited higher inter-subject RSFC variability than sensory-motor networks (Figures 2, S1, S2). One important methodological consideration is that previous studies assumed functional correspondence across subjects after macro-anatomical alignment (Mueller et al., 2013; Chen et al., 2015; Laumann et al., 2015). However, it is well-known that macro-anatomical alignment (or even functional alignment) is not sufficient to achieve perfect functional correspondence across subjects (Fischl et al 1999b; Yeo et al., 2010; 2010b; Harrison et al., 2015; Langs et al., 2016; Glasser et al., 2016). Indeed, a portion of the inter-subject functional connectivity variability observed in previous studies might be the result of residual functional misalignment across subjects (Harrison et al., 2015; Bijsterbosch et al., 2018; Salehi et al., 2018).

Consequently, despite the use of functionally aligned HCP data, we avoided assuming vertex-level functional correspondence provided by the MSMAll functional alignment (Robinson et al., 2014). Instead, the MS-HBM estimated inter-subject and intra-subject variability at the network level, and also explicitly differentiated between inter-subject network spatial variability and inter-subject RSFC variability. This allowed the possibility that for certain networks, inter-subject variability might be attributed to spatial variability, rather than RSFC variability. Nevertheless, we found that networks with higher inter-subject functional connectivity variability (Figures 2, S1, S2) also exhibited greater inter-subject spatial variability (Figures S3 to S5).

Although the MS-HBM approach did not account for inter-site variability, we demonstrated that model parameters estimated from one site can generalize to another site with a different acquisition protocol and scanner (Figures 3, S8, S9, S11). Given the increasing availability of multi-session rs-fMRI from many different research groups (Zuo et al., 2014; Holmes et al., 2015; Poldrack et al., 2015; Filevich et al., 2017; Gordon et al., 2017c), it might be possible to add another layer to the hierarchical model to account for inter-site variability, in addition to intra-subject and inter-subject variability. Furthermore, our experiments did not differentiate between rs-fMRI runs collected within the same session versus rs-fMRI runs collected from different sessions. Another layer could again be inserted into the model to differentiate between within-subject intra-session and within-subject inter-session variability. However, we suspect diminishing returns.

## Conclusions

We developed a multi-session hierarchical Bayesian model (MS-HBM) that differentiated between inter-subject and intra-subject variability when estimating individual-specific cortical network parcellations. Using a single rs-fMRI session (10 min), our approach yielded parcellations comparable to those estimated by two state-of-the-art algorithms using five rs-fMRI sessions (50 min), as evaluated by generalizability to new rs-fMRI data from the same subjects. Furthermore, inter-subject variation in the spatial topography (location and arrangement) of cortical networks could be used to predict inter-subject variation in behavior, suggesting their potential utility as fingerprints of human behavior. Finally, spatial topography estimated by MS-HBM was more effective for behavioral prediction than spatial topography estimated by alternative parcellation approaches.

## Acknowledgement

We like to thank Julien Dubois for providing us with the corrected NEO5 personality scores for the HCP dataset. This work was supported by Singapore MOE Tier 2 (MOE2014-T2-2-016), NUS Strategic Research (DPRT/944/09/14), NUS SOM Aspiration Fund (R185000271720), Singapore NMRC (CBRG/0088/2015), NUS YIA and the Singapore National Research Foundation (NRF) Fellowship (Class of 2017). Computational work for this article was partially performed on resources of the National Supercomputing Centre, Singapore (https://www.nscc.sg). HL was supported by the National Institute of Mental Health (1R01NS091604, P50MH106435) and the Beijing Municipal Science & Technology Commission (Z161100002616009). AJH was supported by the National Institute of Mental Health (573 K01MH099232). XNZ was supported by the National Basic Research (973) Program (2015CB351702), the Natural Science Foundation of China (81471740, 81220108014), the National R&D Infrastructure and Facility Development Program of China (DKA2017-12-02-21) and the Beijing Municipal Science and Tech Commission (Z161100002616023, Z171100000117012). SBE was supported by the Deutsche Forschungsgemeinschaft (DFG, EI 816/11-1), the National Institute of Mental Health (R01-MH074457), the Helmholtz Portfolio Theme “Supercomputing and Modeling for the Human Brain” and the European Union’s Horizon 2020 Research and Innovation Programme under Grant Agreement No. 7202070 (HBP SGA1). Our research also utilized resources provided by the Center for Functional Neuroimaging Technologies, P41EB015896 and instruments supported by 1S10RR023401, 1S10RR019307, and 1S10RR023043 from the Athinoula A. Martinos Center for Biomedical Imaging at the Massachusetts General Hospital. Data were provided by the Brain Genomics Superstruct Project of Harvard University and the Massachusetts General Hospital (Principal Investigators: Randy Buckner, Joshua Roffman, and Jordan Smoller), with support from the Center for Brain Science Neuroinformatics Research Group, the Athinoula A. Martinos Center for Biomedical Imaging, and the Center for Human Genetic Research. Twenty individual investigators at Harvard and MGH generously contributed data to the overall project. Data were also provided by the Human Connectome Project, WU-Minn Consortium (Principal Investigators: David Van Essen and Kamil Ugurbil; 1U54MH091657) funded by the 16 NIH Institutes and Centers that support the NIH Blueprint for Neuroscience Research; and by the McDonnell Center for Systems Neuroscience at Washington University.

## Supplemental Material

This supplemental material is divided into *Supplemental Methods* and *Supplemental Results* to complement the Methods and Results sections in the main text, respectively.

### Supplementary Methods

This section provides additional information about preprocessing, multi-session hierarchical Bayesian model (MS-HBM) and alternative parcellation algorithms that we compared with. Section S1 provides more details about the preprocessing of the GSP and CoRR-HNU data. Section S2 provides more details about the preprocessing of the HCP data. Section S3 provides mathematical details about the MS-HBM. Section S4 describes the algorithms for estimating group-level priors and deriving the individual-specific parcellations and how “free” parameters of the model are set. Section S5 provides more details about the alternative parcellation algorithms that were compared against MS-HBM. Section S6 describes the kernel ridge regression model for behavioral prediction.

### S1. Processing of GSP and CoRR-HNU data

Structural data were processed using FreeSurfer. FreeSurfer constitutes a suite of automated algorithms for reconstructing accurate surface mesh representations of the cortex from individual subjects’ T1 images (Dale et al., 1999; Fischl et al., 2001; Ségonne et al., 2007). The cortical surface meshes were then registered to a common spherical coordinate system (1999b). The GSP subjects were processed using FreeSurfer 4.5.0 (Holmes et al., 2015), while the CoRR-HNU subjects were processed using FreeSurfer 5.3.0.

Resting-state fMRI data of GSP and CoRR-HNU were initially pre-processed with the following steps: (i) removal of first 4 frames, (ii) slice time correction with the FSL package (Jenkinson et al., 2002; Smith et al., 2004), (iii) motion correction using rigid body translation and rotation with the FSL package. The structural and functional images were aligned using boundary-based registration (Greve and Fischl 2009) using the FsFast software package (http://surfer.nmr.mgh.harvard.edu/fswiki/FsFast).

Framewise displacement (FD) and voxel-wise differentiated signal variance (DVARS) were computed using fsl_motion_outliers (Smith et al., 2004). Volumes with FD > 0.2mm or DVARS > 50 were marked as outliers. Uncensored segments of data lasting fewer than 5 contiguous volumes were also flagged as outliers (Gordon et al., 2016). BOLD runs with more than half of the volumes flagged as outliers were removed completely. For the CoRR-HNU dataset, no session (and therefore no subject) was removed. For the GSP subjects, only one run was removed (out of a total of 222 runs). No individuals in the GSP dataset lost an entire session, and therefore, all subjects were retained.

Linear regression using multiple nuisance regressors was applied. Nuisance regressors consisted of global signal, six motion correction parameters, averaged ventricular signal, averaged white matter signal, as well as their temporal derivatives (18 regressors in total). The flagged outlier volumes were ignored during the regression procedure. The data were interpolated across censored frames using least squares spectral estimation of the values at censored frames (Power et al., 2014). Finally, a band-pass filter (0.009 Hz ≤ f ≤ 0.08 Hz) was applied.

The preprocessed fMRI data was projected onto the FreeSurfer fsaverage6 surface space (2mm vertex spacing). The projected fMRI data was smoothed using a 6mm full-width half-maximum (fwhm) kernel and then downsampled onto fsaverage5 surface space (4mm vertex spacing). Smoothing on the fsaverage6 surface, rather than in the volume minimized the blurring of fMRI signal across sulci.

### S2. Processing of HCP data

Details of the HCP preprocessing can be found elsewhere (HCP S900 manual; Van Essen et al. 2012b; Glasser et al. 2013; Smith et al. 2013). Of particular importance is that the rs-fMRI data has been projected to the fs_LR surface space (Van Essen et al. 2012a), smoothed by 2mm fwhm and denoised with ICA-FIX (Salimi-Khorshidi et al. 2014; Griffanti et al., 2014) and aligned with MSMAll (Robinson et al., 2014).

However, recent studies have shown that ICA-FIX does not fully eliminate global and head-motion related artifacts (Burgess et al., 2016; Siegel et al., 2016). Therefore, further processing steps were performed on the rs-fMRI data in fs_LR surface after ICA-FIX denoising, which included nuisance regression, motion censoring and interpolation, and band-pass filtering. Volumes with FD > 0.2mm or DVARS > 75, as well as uncensored segments of data lasting fewer than 5 contiguous volumes were flagged as outliers. BOLD runs with more than half the volumes flagged as outliers were completely removed. Consequently, 56 subjects were removed. Furthermore, for this work, only subjects with all four runs remaining (N = 676) were considered.

Nuisance regression utilized regressors consisting of global signal, six motion parameters, averaged ventricular signal, averaged white matter signal, and their temporal derivatives (18 regressors in total). The outlier volumes were ignored during the regression procedure. The data were interpolated across censored frames using least squares spectral estimation (Power et al., 2014). A band-pass filter (0.009 Hz ≼ f ≼ 0.08 Hz) was then applied to the data. Finally, the data was smoothed by 4mm fwhm. Given that the HCP team has already smoothed the data by 2mm, this results in an effective smoothing of 6mm fwhm.

### S3. Mathematical model (MS-HBM)

In this section, we describe our model for individual-level parcellation of the cerebral cortex. We assume a common surface coordinate system, where the cerebral cortex is represented by left and right hemisphere spherical meshes such as FreeSurfer fsaverage surface meshes. Each mesh consists of a collection of vertices and edges connecting neighboring vertices into triangles (https://en.wikipedia.org/wiki/Triangle_mesh).

Let *N* denote the total number of vertices, *T* denote the number of resting-state fMRI (rs-fMRI) sessions, *S* denote the number of subjects, *L* denote the number of networks, and *N_n_* denote the neighboring vertices of vertex *n* (as defined by the cortical mesh). For each subject *s* and session *t*, there is a preprocessed rs-fMRI time course associated with each vertex *n*. For each subject *s*, there is an unknown parcellation label 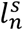 at vertex *n*. Note that the parcellation label is assumed to be the same across sessions (hence there is no index on the session). In this work, we use 1: *S* to denote a set of subjects {1, 2, …, *S*}, 1: *T* to denote a set of sessions {1, 2, …, *T*}, 1: *N* to denote a set of vertices {1, 2, …, *N*}, 1: *L* to denote a set of parcellation labels {1, 2, …, *L*}.

For each subject *s* at a particular session *t*, we computed the functional connectivity profile of each vertex (of the cortical mesh) by correlating the vertex’s fMRI time course with the time courses of uniformly distributed cortical regions of interests^1^ (ROIs). For the GSP and HNU datasets, the preprocessed data were in fsaverage5 surface space. In this case, the ROIs consisted of 1175 vertices approximately uniformly distributed across the two hemispheres (Yeo et al., 2011). For the HCP dataset, the preprocessed data is in fs_LR32k surface space. In this case, the ROIs consisted of 1483 vertices spaced approximately uniformly distributed across the two hemispheres. Each vertex’s connectivity profile was binarized (see Methods in main manuscript) and normalized to unit length. Let 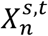 denote the binarized, normalized functional connectivity profile of subject *s* at vertex *n* during session *t*. Let *D* denote the total number of ROIs and hence the length of 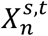. We denote the connectivity profiles from all sessions of all subjects at all cortical vertices as 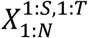.

Figure 1 (main text) illustrates the schematic of the multi-session hierarchical Bayesian model (MS-HBM). Following previous work (Yeo et al., 2011), the functional connectivity profile 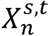 of subject *s* from a session *t* at vertex n is assumed to be generated from a von Mises-Fisher distribution,

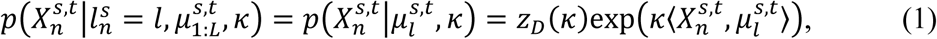

where 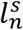 is the parcellation label at vertex *n* of subject *s*, and 〈,〉 denote inner product. 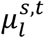 and κ are the mean direction and concentration parameter of the von Mises-Fisher distribution for network label *l* of subject *s* during session *t*. 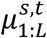 are the mean directions for networks 1 to *L*. We can think of 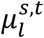 as the mean connectivity profile of network label *l* normalized to unit length. If functional connectivity profile 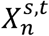 is similar to mean connectivity profile 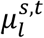 (i.e., 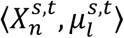 is big), then vertex *n* is more likely to be assigned to network *l*.

The concentration parameter κ controls the variability of the functional connectivity profiles within each network. A higher κ results in a lower dispersion (i.e., lower variance), which means that vertices belonging to the same network are more likely to possess functional connectivity profiles that are close to the mean connectivity profile of the network. In theory, it might make sense for κ to be different across networks because certain networks might exhibit more inter-region functional connectivity variability (Gordon et al., 2017a). However, in practice, certain networks (e.g., limbic networks) with low signal-to-noise ratio (SNR) would end up with very low κ, resulting in their encroachment into the peripheries of other networks. Therefore, κ was set to be the same across all networks, subjects and sessions. Finally, *z*_*D*_(κ) is a normalization constant to ensure a valid probability distribution (Banerjee et al., 2005):

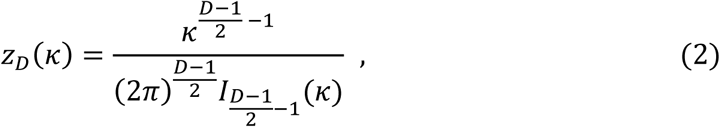

where 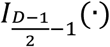 is the modified Bessel function of the first kind with order 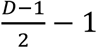

To model intra-subject functional connectivity variability, we assume a conjugate prior on the subject-specific and session-specific mean connectivity profiles 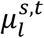, which turns out to also be a von Mises-Fisher distribution:

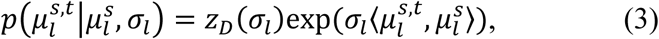

where 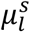 and σ_*l*_ are the mean direction and concentration parameter of the von Mises-Fisher distribution for network label *l* of subject *s*. We can think of 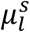 as the individual-specific functional connectivity profile of network *l* of subject *s*. The concentration parameter σ_*l*_ controls how much the session-specific mean direction 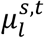 of subject *s* during session *t* can deviate from the subject-specific mean direction 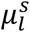. A higher σ_*l*_ would imply lower intra-subject functional connectivity variability across sessions. σ_*l*_ is network-specific but is assumed to be the same for all subjects.

To model inter-subject functional connectivity variability, we assume a conjugate prior on the subject-specific mean connectivity profiles 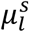, which is again a von Mises-Fisher distribution whose mean direction corresponded to the group-level mean direction *μ* ^*g*^:

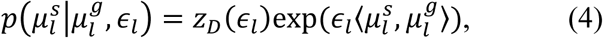

where *μ* ^*g*^ and ∈_*l*_ are the mean direction and concentration parameter of the von Mises-Fisher distribution for network label *l*. We can think of 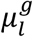 as the group-level functional connectivity profile of network *l*. The concentration parameter ∈_*l*_ controls how much the individual-specific connectivity profile 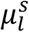 can deviate from the group-level connectivity profile 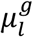. A higher ∈_*l*_ would imply lower inter-subject functional connectivity variability across subjects.

Because the functional connectivity profiles of individual subjects are generally very noisy, we impose a MRF prior on the hidden parcellation labels 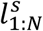

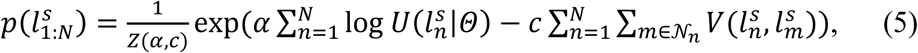

where *Z*(*α, c*) is a normalization term (partition function) to ensure 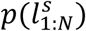 is a valid probability distribution. 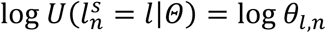 is a singleton potential encouraging certain vertices to be associated with certain labels. 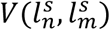 is a pairwise potential (Potts model) encouraging neighboring vertices to have the same parcellation labels:

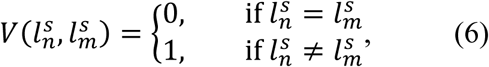

The parameters *α* and *c* are tunable parameters greater than zero, and control the tradeoffs between the various terms in the generative model. Assuming that *α* = 1 and *c* = 0, then *θ*_*l,n*_ can be interpreted as the probability of label *l* occurring at vertex *n* of subject *s*.

### S4. Model estimation for MS-HBM

In this section, we describe how model parameters are estimated from a training set and a validation set (Section S2.1), and how the parameters can be used to parcellate a new subject (Section S2.2). Throughout the entire section, we assume that the number of networks *L* = 17 without loss of generality.

#### S4.1 Learning model parameters

Our goal is to estimate the model parameters 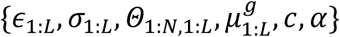 from a training set and a validation set of binarized and normalized functional connectivity profiles, which can then be utilized for estimating individual-specific parcellations in unseen data of new subjects (Section S2.2). As a reminder, ∈_1:*L*_ is a group prior representing inter-subject functional connectivity variability, σ_1:*L*_ is a group prior corresponding to intra-subject functional connectivity variability, *Θ*_1:*N*,1:*L*_ is a group prior representing inter-subject spatial variability and reflects the probability of a network occurring at particular spatial location, and 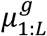 is the group-level connectivity profile for each network. The parameters *α* and *c* tradeoff between various terms in the generative model. Because the partition function *Z*(*α, c*) (Eq. (5)) is NP-hard to compute, for computational efficiency, we first assume *α* = 1, *c* = 0 in order to estimate 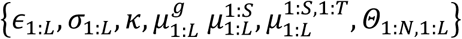 from the training dataset^2^. Under this scenario, *Z*(*α, c*) = 1, and *θ*_*l,n*_ can be interpreted as the probability of label *l* occurring at vertex *n* of subject *s*. The tunable parameters *α* and *c* are then estimated in the validation set using a grid search.

##### S4.1.1 Estimating 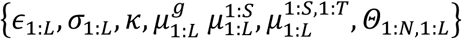 from training set

Given observed binarized, normalized functional connectivity profiles 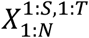 from the training set, we seek to estimate 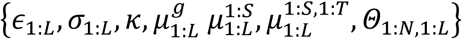 using Expectation-Maximization (EM). As previously explained, we assume *α* = 1, *c* = 0.

Let 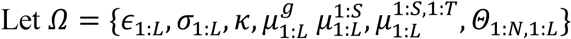. We consider the following maximum-a-posterior (MAP) estimation problem:

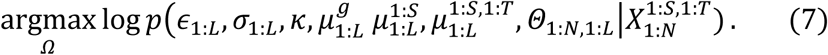

Assuming a uniform (improper) prior on {*Θ*_1:*N*,1:*L*_, κ, σ_1:*L*_, ∈_1:*L*_}, the MAP problem can be written as

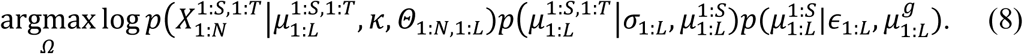

We then introduce the parcellation labels 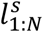 for each subject *s* as latent variables, and use Jensen’s inequality to define a lower bound ℒ(*λ, Ω*), where 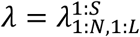 are the parameters of the *q* functions 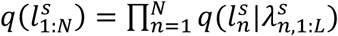:

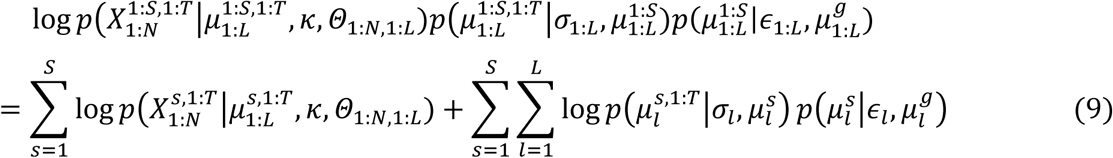

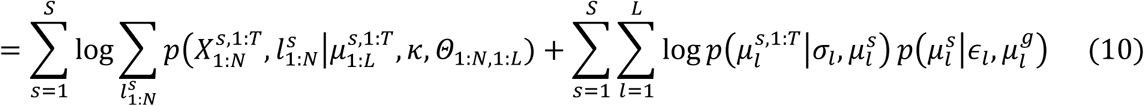

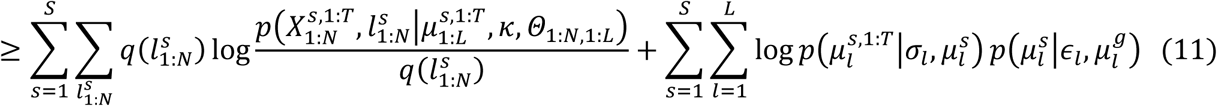

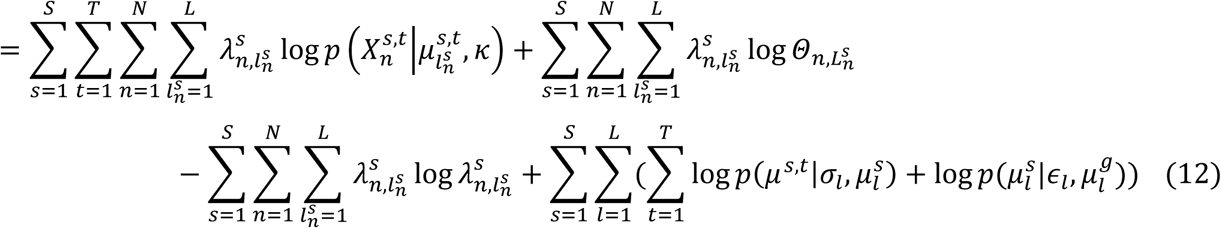

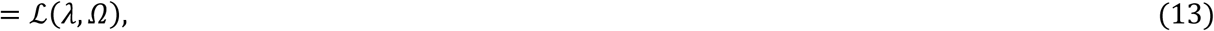

where equality is achieved when 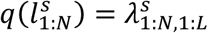 are the posterior probability of the individual-specific parcellation of subject *s* given the parameters *Ω*. Therefore, instead of maximizing the original MAP problem (Eq. (7)), we instead maximize the lower bound:

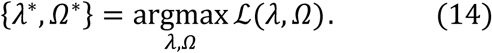

In the E-step, we fix 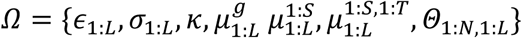, and estimate *λ*

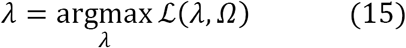

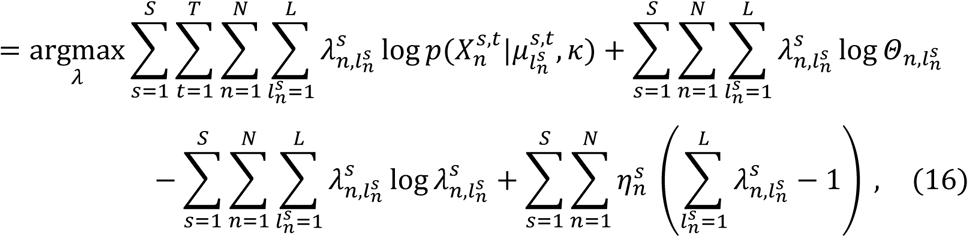

where 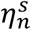 are the Lagrange multipliers enforcing the constraint 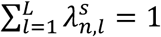. Optimizing Eq. (16), we

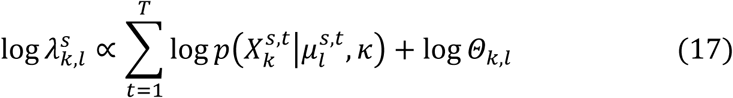

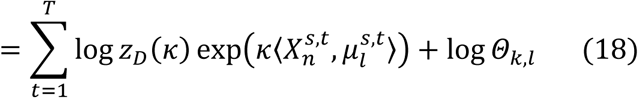

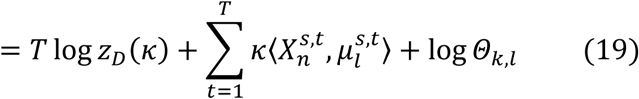

In the M-step, we fix *λ* and estimate *Ω*:

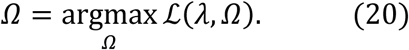

By using the constraints that 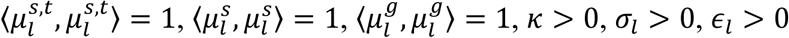, κ > 0, σ_*l*_ > 0, ∈_*l*_ > 0, and differentiating ℒ(*λ, Ω*) with respect to 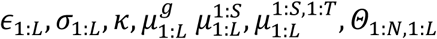, *Θ*_1:*N*,1:*L*_, and setting the derivatives to zero, we get the following update equations:

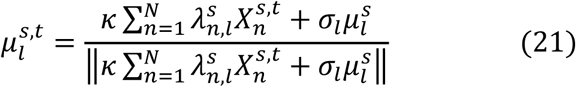

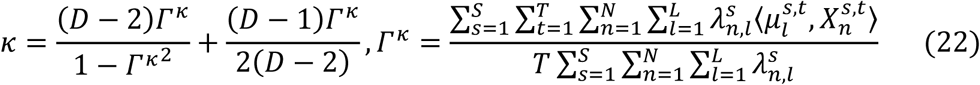

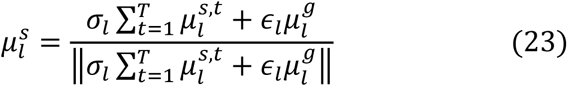

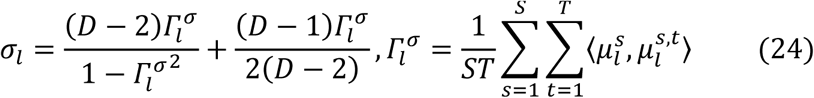

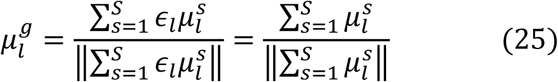

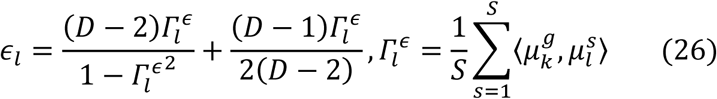

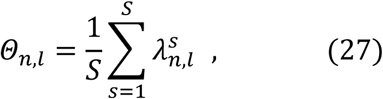

where *D* is the length of 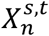 (i.e., number of ROIs in each functional connectivity profile), *S* is the number of subjects, *T* is the number of sessions, and ?·? corresponds to the *l*_2_-norm. Therefore, the estimate of the functional connectivity profile 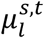 (Eq. (21)) of network *l* of subject *s* during session *t* is the weighted sum of the average time course of vertices constituting network *l* of subject *s* during session 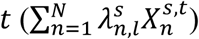 and the subject-specific mean direction 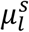, with weights κ and σ_*l*_ for each term, normalized to be unit norm. If σ_*l*_ is much greater than κ, then 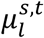 is more likely to be dominated by subject-specific mean direction 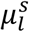, which means that the functional connectivity profile of network *l* is highly stable across sessions. Similarly, the estimate of the functional connectivity profile 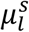 (Eq. (23)) of network *l* of subject *s* is the weighted sum of the average session-specific mean directions across all sessions for network *l* of subject 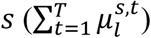 and the group-level mean direction 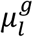, with weights σ_*l*_ and ∈_*l*_ for each term, normalized to be unit norm. If ∈_*l*_ is much greater than σ_*l*_, then 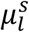 is more likely to be dominated by group-level mean direction 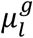, which means that the functional connectivity profile of network *l* is highly stable between subjects. Finally, the estimate of the group-level functional connectivity profile 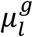 (Eq. (25)) of network *l* is the sum of the subject-specific mean directions across all subjects for network 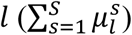, normalized to be unit norm. The estimate of *Θ*_*n,l*_ (Eq. (27)) is the posterior probability of network *l* being assigned to vertex *n*, averaged across all the subjects.

Given the training set, the algorithm first estimates a group-level parcellation (Yeo et al., 2011), which is then used to initialize the EM algorithm. The EM algorithm iterates E-step (Eq. (19)) and M-step (Eqs. (21-27)) till convergence. We note that the update equations (Eqs. (21-27)) in the M-step are dependent on each other. Therefore, within the M-step, the update equations (Eqs. (21-27) are iterated till convergence.

##### S4.1.2 Estimating tunable parameters c and α

In the previous subsection (Section S2.1.1), the training set was used to estimate 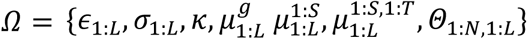, assuming *α* = 1, *c* = 0. To tune the parameters *c* and *α,* we assume access to a validation set.

Recall that each subject in the validation set has multiple rs-fMRI sessions. We consider *c* ∈ {10,20,30,40, 50,60} and *α* ∈ {100,150,200,250}. For a given pair of (*c, α*), and given 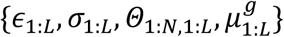 estimated from the training set, we estimate for each subject in the validation set, the individual-specific parcellation based on a subset of rs-fMRI sessions (see Section S2.2 for algorithm). Resting-state homogeneity (Eq. (1) in main text) is then computed in the remaining rs-fMRI sessions of the validation subjects. The pair of (*c, α*) with the highest homogeneity in the unseen rs-fMRI sessions of the validation subjects is then utilized for parcellating new subjects.

In the case of the GSP data, the optimal pair of parameters is *c* = 30 and *α* = 200. In the case of the HCP data, the optimal pair of parameters is *c* = 40 and *α* = 200. Note that we do expect the parameters to be different between the GSP and HCP datasets because of resolution differences between the fsaverage5 and fs_LR32k surface meshes.

Throughout the paper (main text), the reported quality (Figures 5, 6 and 7) of the individual-specific parcellations was evaluated using subjects not used to tune the parameters. For example, in the case of the CoRR-HNU subjects (Figures 5 and 6), the model parameters were estimated from the GSP training and validation sets. In the case of the HCP data (Figures 6 and 7), model parameters were estimated the HCP training and validation sets, while the reported quality of the individual-specific parcellations was evaluated using the HCP test set.

#### S4.2 Individual-level parcellation estimation

Using parameters 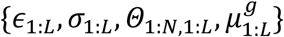 estimated from the training set (Section S2.1.1), and for a particular pair of (*c, α*), we can estimate the individual-specific parcellation 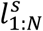 of a new subject *s* with *T* sessions by employing the variational Bayes expectation maximization (VBEM) algorithm.

Let 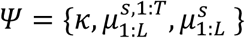. We consider the following maximum-a-posterior (MAP) estimation problem:

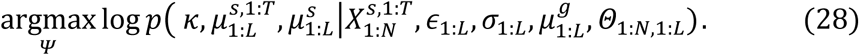

Assuming a uniform (improper) prior on κ, and by introducing the parcellation labels 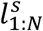 of the new subject *s* as latent variables, the lower bound ℒ(*λ, ψ*) of the MAP problem (Eq. (28)) can be written as:

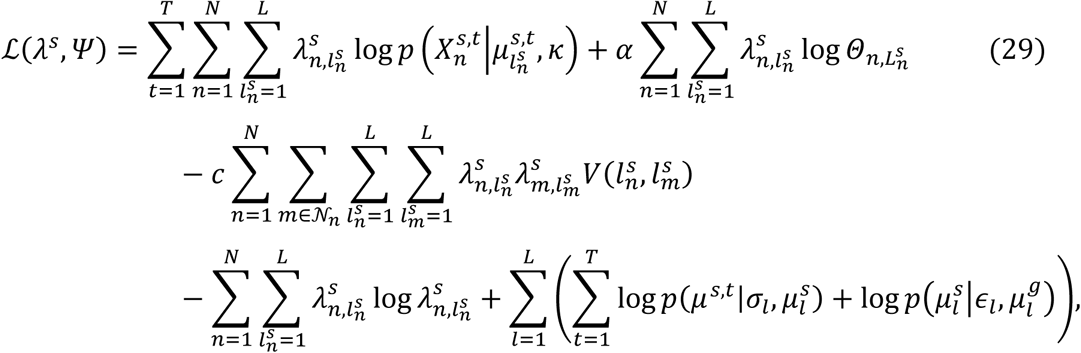

where equality is achieved when *λ*^*s*^ is the posterior probability of the individual-specific parcellation of subject *s* given the parameters *ψ*. Similar to Section S2.1.1, we can maximize the lower bound (Eq. (29)) by iteratively updating *λ*^*s*^ and *ψ*. Unlike Section S2.1.1, we cannot compute the exact posterior probability *λ*^*s*^ because of the pairwise potentials in the Markov random field (Wainwright and Jordan, 2008). Using the mean-field approximation (Wainwright and Jordan, 2008), an approximate posterior probability *λ*^*s*^ is estimated in the variational E-step, while *ψ* is updated in the variational M-step.

More specifically, in the variational E-step, *ψ* is fixed and *λ*^*s*^ is estimated as follows:

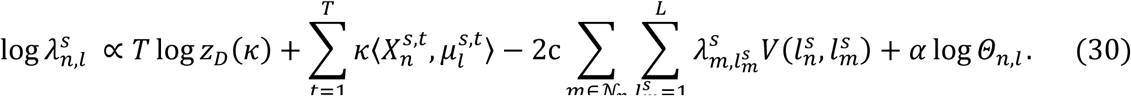

In the variational M-step, *λ*^*s*^ is fixed and 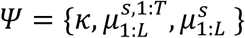 is estimated as follows:

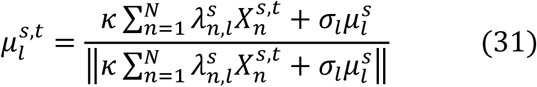

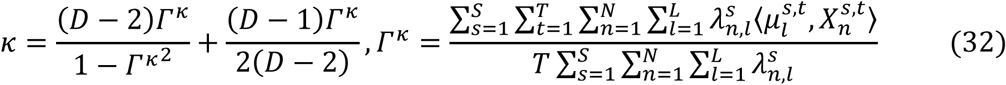

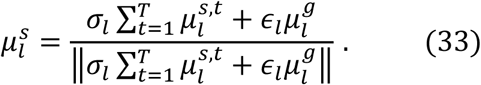

Once the VBEM algorithm converges, vertex *n* of subject *s* will be assigned to label *l* with the highest (approximate) posterior probability.

### S5. Alternative parcellation approaches

We compared the MS-HBM with four alternative parcellation approaches. The first approach was to apply the population-level parcellation (Yeo et al., 2011) to individual subjects. We will refer to this approach as “Yeo2011”. For the second approach, recall that the population-level parcellation algorithm was an expectation-maximization (EM) algorithm, which iteratively computed a network connectivity profile based on vertices assigned to the same network (M-step) and then re-assigned the network membership of vertices based on the similarity between each vertex’s connectivity profile and the network connectivity profile (E-step). Using the network connectivity profiles from the Yeo2011 population-level parcellation, we could estimate networks in an individual subject by assigning a network label to each vertex based on the similarity between the vertex’s connectivity profile (for that subject) and the population-level network connectivity profile (i.e., E-step). Since this approach is analogous to the ICA back-projection algorithm (Calhoun et al., 2009; Beckmann et al., 2009; Filippini et al., 2009; Zuo et al., 2010; Calhoun and Adali 2012), we will refer to this second alternative approach as “YeoBackProject”.

We also implemented the influential individual-specific parcellation algorithm of Gordon and colleagues (Gordon et al., 2017a; Gordon et al., 2017b), where the binarized functional connectivity map of each cortical vertex was matched to binarized network templates derived from the Yeo2011 population-level parcellation. Care was taken to verify that our implementation was consistent with Gordon’s algorithm (whose code is publicly available). We refer to this third approach as “Gordon2017”.

Finally, we considered the prominent individual-specific parcellation algorithm of Wang and colleagues (Wang et al., 2015), which adapted the Yeo2011 population-level parcellation to an individual subject, while accounting for inter-subject RSFC variability and SNR characteristics of the subject’s rs-fMRI data. However, scanner noise is just one component contributing to intra-subject variability. As shown by others (Mueller et al., 2013; Laumann et al., 2015) and also in our results, brain networks with the highest intra-subject variability do not correspond to low SNR regions. We refer to this fourth approach as “Wang2015”.

### S6. Behavioral prediction model

In this section, we describe our model for behavioral prediction based on individual differences in the spatial arrangement of cortical networks. Kernel regression (Murphy et al., 2012) was utilized to predict each behavioral phenotype in individual subjects. Suppose we have *M* training subjects, *y*_*i*_ is the behavioral measure (e.g., fluid intelligence) and *l*_*i*_ is the individual-specific parcellation of the *i*-th training subject. Given {*y*_1_, *y*_2_, …, *y*_*M*_} and {*l*_1_, *l*_2_, …, *l*_*M*_}, the kernel regression model will be:

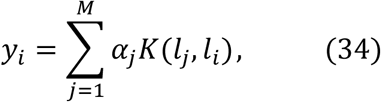

where *K*(*l*_*j*_, *l*_*i*_) is the Dice overlap coefficient between corresponding networks of the *i*-th and *j*-th training subjects, averaged across 17 networks. The classical way to estimate *α* in Eq. (34) is to minimize the quadratic cost:

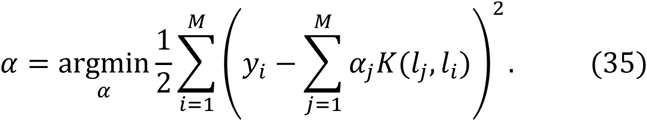

Defining ***y*** = [*y*_1_, *y*_2_, …, *y*_*M*_]^*T*^, ***α*** = [*α*_1_, *α*_2_, …, *α*_*M*_]^*T*^ and 𝕂 to be an *M* × *M* matrix, whose (*j, i*)-th element is *K*(*l*_*j*_, *l*_*i*_), Eq. (35) can be written as:

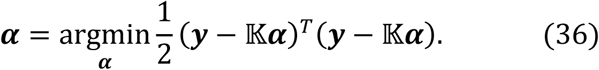

Differentiating Eq. (36) with respect to ***α***, we can get

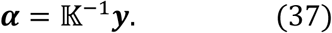

To predict the behavior measure *y*_*s*_ (e.g., fluid intelligence) of a test subject *s* with its individual-specific parcellation *l*_*s*_, we can compute *K*(*l*_*i*_, *l*_*s*_), which is the Dice overlap coefficient between corresponding networks of subject *s* and *i*-th training subject, averaged across 17 networks. The predicted behavior measure *y*_*s*_ can be calculated as

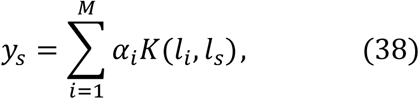

where *α*_*i*_ is estimated by Eq. (37). If we denote ***K***_*s*_ = [*K*(*l*_1_, *l*_*s*_), *K*(*l*_2_, *l*_*s*_), …, *K*(*l*_*M*_, *l*_*s*_)], then Eq. (38) can be written as:

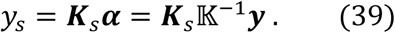

In practice, 𝕂^-1^ is a symmetric matrix whose diagonal elements are roughly the same and ∼100 times larger than the off-diagonal elements. Therefore, the predicted behavior measure *y*_*s*_ can be seen as the weighted average of the behaviors of the training subjects: 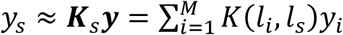. If the individual-specific parcellation *l*_*s*_ of test subject *s* is more similar to the parcellation of training subject *i* than training subject *j*, then weight *K*(*l*_*i*_, *l*_*s*_) will be larger than *K*(*l*_*j*_, *l*_*s*_), and so *y*_*s*_ will be more similar to *y*_*i*_ than *y*_*j*_.

To reduce overfitting, an *l*_2_-regularization term (i.e., kernel ridge regression) is typically added to cost function (Eq. (36)), resulting in a new regularized cost function:

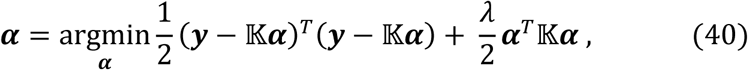

where *λ* is a tuning parameter, which controls the importance of the regularization term. Differentiating Eq. (40) with respect to ***α***, we get

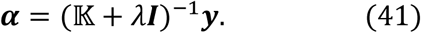

To predict the behavior measure *y*_*s*_ (e.g., fluid intelligence) of a test subject *s*, Eq. (41) is substituted into Eq. (38), resulting in

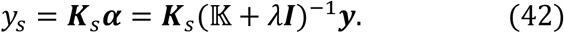

## Supplemental Results

**Table S1.** Lookup table showing the original HCP variable names with the corresponding descriptive labels used in the manuscript. More details of the behavioral measures can be found in the HCP data dictionary.

| **Description** | **HCP field** |
| --- | --- |
| Visual Episodic Memory | PicSeq_Unadj |
| Cognitive flexibility (DCCS) | CardSort_Unadj |
| Inhibition (Flanker task) | Flanker_Unadj |
| Fluid Intelligence (PMAT) | PMAT24_A_CR |
| Reading (pronunciation) | ReadEng_Unadj |
| Vocabulary (picture matching) | PicVocab_Unadj |
| Processing Speed | ProcSpeed_Unadj |
| Delay Discounting | DDisc_AUC_40K |
| Spatial orientation | VSLOT_TC |
| Sustained Attention - Sens. | SCPT_SEN |
| Sustained Attention - Spec. | SCPT_SPEC |
| Verbal Episodic Memory | IWRD_TOT |
| Working Memory (list sorting) | ListSort_Unadj |
| Cognitive status (MMSE) | MMSE_Score |
| Sleep quality (PSQI) | PSQI_Score |
| Walking endurance | Endurance_Unadj |
| Walking Speed | GaitSpeed_Comp |
| Manual dexterity | Dexterity_Unadj |
| Grip strength | Strength_Unadj |
| Odor identificaiton | Odor_Unadj |
| Pain Interference Survey | PainInterf_Tscore |
| Taste intensity | Taste_Unadj |
| Contrast Sensitivity | Mars_Final |
| Emotional Face Matching | Emotion_Task_Face_Acc |
| Arithmetic | Language_Task_Math_Avg_Difficulty_Level |
| Story comprehension | Language_Task_Story_Avg_Difficulty_Level |
| Relational processing | Relational_Task_Acc |
| Social Cognition - random | Social_Task_Perc_Random |
| Social Cognition - interaction | Social_Task_Perc_TOM |
| Working Memory (n-back) | WM_Task_Acc |
| Agreeableness (NEO) | NEOFAC_A |

Table S1 (cont.). Lookup table showing the original HCP variable names with the corresponding descriptive labels used in the manuscript. More details of the behavioral measures can be found in the HCP data dictionary.
| **Description** | **HCP field** |
| --- | --- |
| Openness (NEO) | NEOFAC_O |
| Conscientiousness (NEO) | NEOFAC_C |
| Neuroticism (NEO) | NEOFAC_N |
| Extraversion (NEO) | NEOFAC_E |
| Emot. Recog. - Total | ER40_CR |
| Emot. Recog. - Angry | ER40ANG |
| Emot. Recog. - Fear | ER40FEAR |
| Emot. Recog. - Happy | ER40HAP |
| Emot. Recog. - Neutral | ER40NOE |
| Emot. Recog. - Sad | ER40SAD |
| Anger - Affect | AngAffect_Unadj |
| Anger - Hostility | AngHostil_Unadj |
| Anger - Aggression | AngAggr_Unadj |
| Fear - Affect | FearAffect_Unadj |
| Fear - Somatic Arousal | FearSomat_Unadj |
| Sadness | Sadness_Unadj |
| Life Satisfaction | LifeSatisf_Unadj |
| Meaning & Purpose | MeanPurp_Unadj |
| Positive Affect | PosAffect_Unadj |
| Friendship | Friendship_Unadj |
| Loneliness | Loneliness_Unadj |
| Perceived Hostility | PercHostil_Unadj |
| Perceived Rejection | PercReject_Unadj |
| Emotional Support | EmotSupp_Unadj |
| Instrument Support | InstruSupp_Unadj |
| Perceived Stress | PercStress_Unadj |
| Self-Efficacy | SelfEff_Unadj |

**Table S2.** Task-relevant networks of 13 cognitive measures based on NeuroSynth database. The “initial search terms” were our initial queries in the NeuroSynth database for each cognitive measure. The “final search terms” were the terms that we finally utilized to retrieve the forward inference map. There was no appropriate search term for PicVocab_Unadj (picture vocabulary) and ProcSpeed_Unadj (processing speed). Each forward map was projected to fsLR surface space and compared with the group-level parcellation estimated from the HCP training set (Figure 2A) to select the task-relevant networks.

| HCP field | Initial search terms | Relevant (final) NeuroSynth term | Task-relevant networks |
| --- | --- | --- | --- |
| PicSeq_Unadj | Visual episodic memory | Episodic memory | DefaultA; ControlA |
| CardSort_Unadj | Cognitive flexibility, dimensional sort, Wisconsin card sorting, task switching | Cognitive control | DorsalAttentionA; VentralAttentionB; ControlA |
| Flanker_Unadj | Flanker task | Inhibitory control | ControlA; ControlC; VentralAttentionB |
| PMAT24_A_CR | Fluid intelligence | Intelligence | DorsalAttentionA; ControlA |
| ReadEng_Unadj | Reading decoding, pronunciation | Reading | DorsalAttentionA; ControlA; TemporalParietal |
| PicVocab_Unadj | Picture matching, vocabulary comprehension, vocabulary | NA | NA |
| ProcSpeed_Unadj | Processing speed | NA | NA |
| DDisc_AUC_40K | Delay discounting | Impulsivity | ControlA; ControlC; VentralAttentionB; |
| VSLOT_TC | Spatial orientation, orientation, visual spatial | Visual perception | VisualA |
| SCPT_SEN | Sustained attention | Attention | DorsalAttentionA; ControlA; VisualA |
| SCPT_SPEC | Sustained attention | Attention | DorsalAttentionA; ControlA; VisualA |
| IWRD_TOT | Verbal episodic memory, word memory | Episodic memory | DefaultA; ControlA |
| ListSort_Unadj | Working memory | Working memory | DorsalAttentionA; ControlA; |

**Table S3.** Average prediction accuracies for different sets of behavioral measures (minimally correlated set of 5 behaviors, 58 behavioral measures, 13 cognitive measures, NEO-5 personality measures, emotion recognition measures and emotional measures) across different parcellation approaches. Prediction was based on individual-specific network topography. The mean accuracy and standard deviation was calculated across 100 20-fold cross-validations. The percentage improvement and number of cross-validations that MS-HBM outperforms other approaches across 100 20-fold cross-validations was reported.

|  |  | Minimally correlated set of 5 behaviors | All 58 behaviors | 13 Cognitive measures |
| --- | --- | --- | --- | --- |
| Accuracy across 100 20-fold CVs (mean ± std) | MS-HBM | 0.1327±0.0065 | 0.0803±0.0032 | 0.1321±0.0053 |
| | YeoBackProject | 0.1036±0.0080 | 0.0728±0.0032 | 0.1057±0.0060 |
| | Gordon2017 | 0.0830±0.0080 | 0.0655±0.0036 | 0.0545±0.0062 |
| | Wang2015 | 0.1039±0.0080 | 0.0755±0.0033 | 0.1202±0.0054 |
| Percentage improvement (#CV splits MS-HBM outperforms other approaches) | MS-HBM vs YeoBackProject | 28.55% (100/100) | 10.35% (100/100) | 25.17% (100/100) |
|  | MS-HBM vs Gordon2017 | 61.07% (100/100) | 22.88% (100/100) | 145.12% (100/100) |
|  | MS-HBM vs Wang2015 | 28.20% (100/100) | 6.43% (95/100) | 10.05% (97/100) |

Table S3. Average prediction accuracies for different sets of behavioral measures (minimally correlated set of 5 behaviors, 58 behavioral measures, 13 cognitive measures, NEO-5 personality measures, emotion recognition measures and emotional measures) across different parcellation approaches.
|  |  | NEO-5 | Emotional recognition (ER) | Emotional measures excluding ER |
| --- | --- | --- | --- | --- |
| Accuracy across 100 20-fold CVs (mean ± std) | MS-HBM | 0.0955±0.0085 | -0.0445±0.0101 | **0.1038±0.0070** |
| | YeoBackProject | **0.1027±0.0084** | -0.0170±0.0092 | 0.0781±0.0070 |
| | Gordon2017 | 0.0820±0.0080 | **0.0297±0.0092** | 0.0794±0.0072 |
| | Wang2015 | 0.0772±0.0090 | 0.0068±0.0093 | 0.0802±0.0069 |
| Percentage improvement (#CV splits MS-HBM outperforms other approaches) | MS-HBM vs YeoBackProject | -6.82% (16/100) | -264.70% (0/100) | 33.46% (100/100) |
|  | MS-HBM vs Gordon2017 | 17.20% (94/100) | -269.91% (0/100) | 31.36% (100/100) |
|  | MS-HBM vs Wang2015 | 24.61% (99/100) | -913.27% (0/100) | 29.85% (100/100) |

**Table S4.** Average prediction accuracies of DVARS and FD across different parcellation approaches. Prediction was performed based on individual-specific network topography without regressing any nuisance covariates. The mean accuracy and standard deviation was calculated across 100 20-fold cross-validations.

| | Accuracy (mean ± std) across 100 20-fold cross-validations | | | |
| --- | --- | --- | --- | --- |
|  | MS-HBM | YeoBackProject | Gordon | Wang |
| DVARS | 0.3157±0.0182 | 0.4283±0.0169 | 0.2115±0.0180 | 0.3621±0.0175 |
| FD | 0.2120±0.0153 | 0.2733±0.0140 | 0.2565±0.0136 | 0.2066±0.0171 |

---

**Figure S1.**
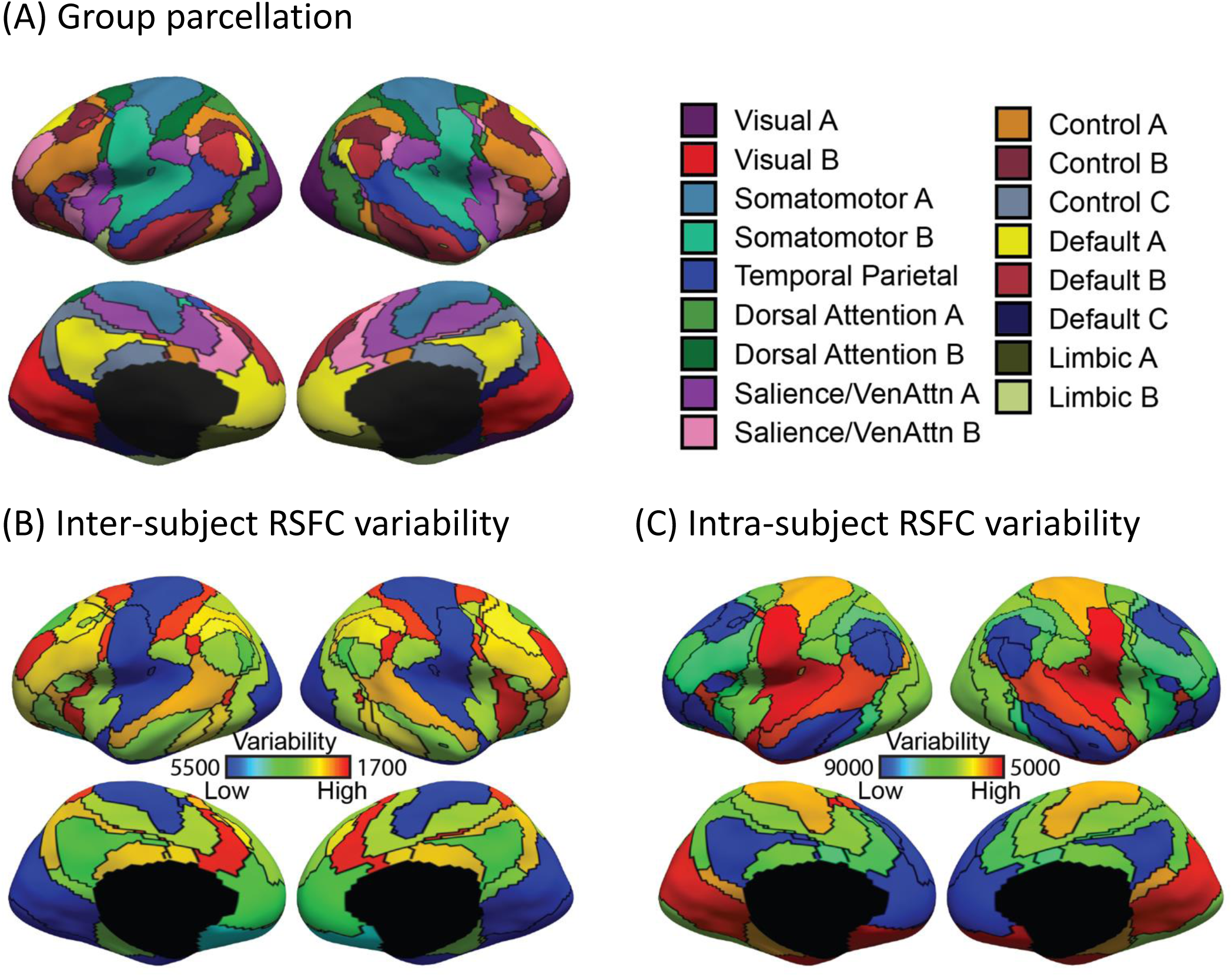
Sensory-motor networks exhibit lower inter-subject, but higher intra-subject, functional connectivity variability than association networks in the GSP training set. (A) 17-network group-level parcellation. (B) Inter-subject functional connectivity variability for different cortical networks. (C) Intra-subject functional connectivity variability for different cortical networks. Note that (B) and (C) correspond to the ϵ_*l*_ and σ_*l*_ parameters in Figure 1, where higher values indicate lower variability.

**Figure S2.**
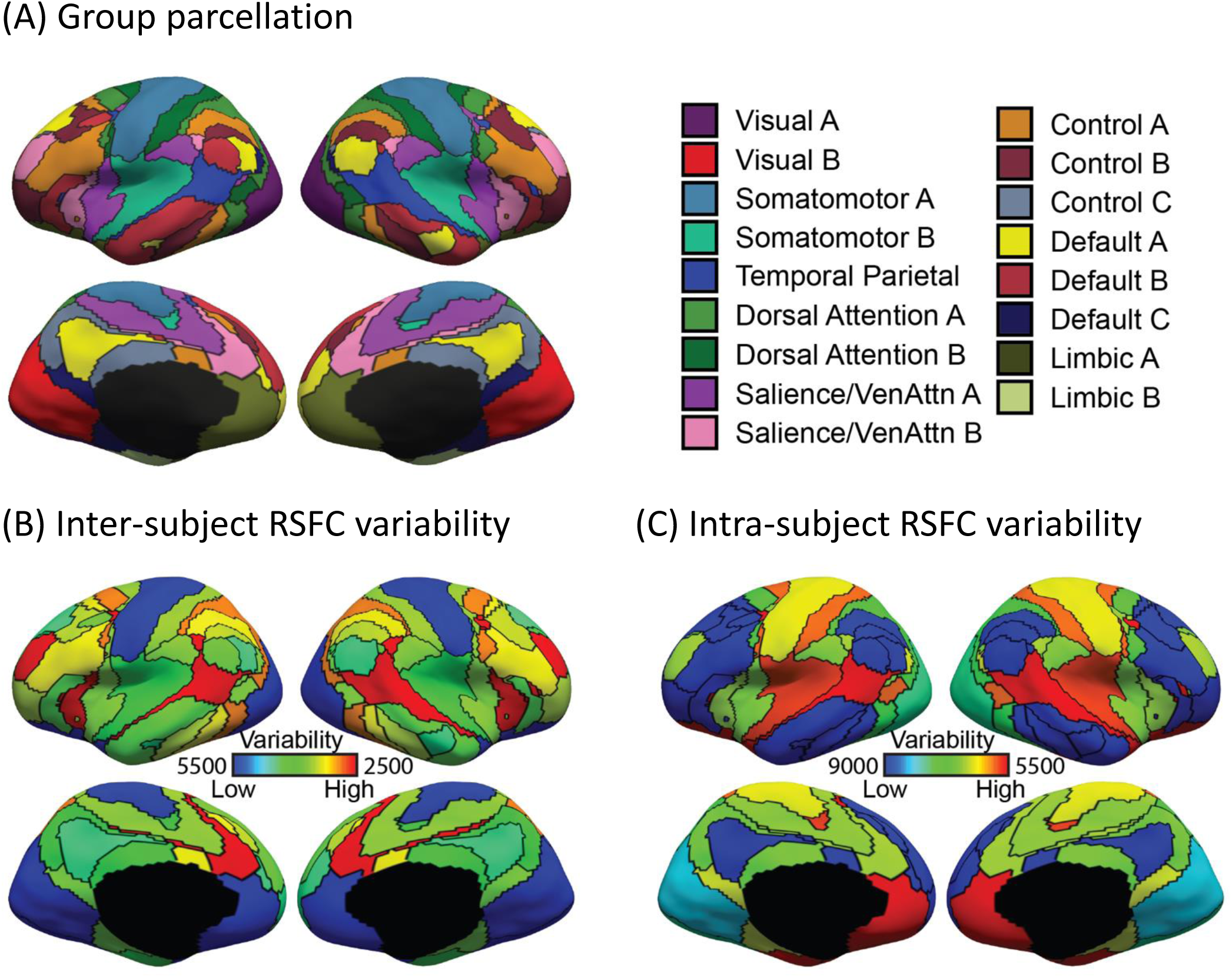
Sensory-motor networks exhibit lower inter-subject, but higher intra-subject, functional connectivity variability than association networks in the CoRR-HNU dataset. (A) 17-network group-level parcellation. (B) Inter-subject functional connectivity variability for different cortical networks. (C) Intra-subject functional connectivity variability for different cortical networks. Note that (B) and (C) correspond to the ϵ_*l*_ and σ_*l*_ parameters in Figure 1, where higher values indicate lower variability.

**Figure S3.**
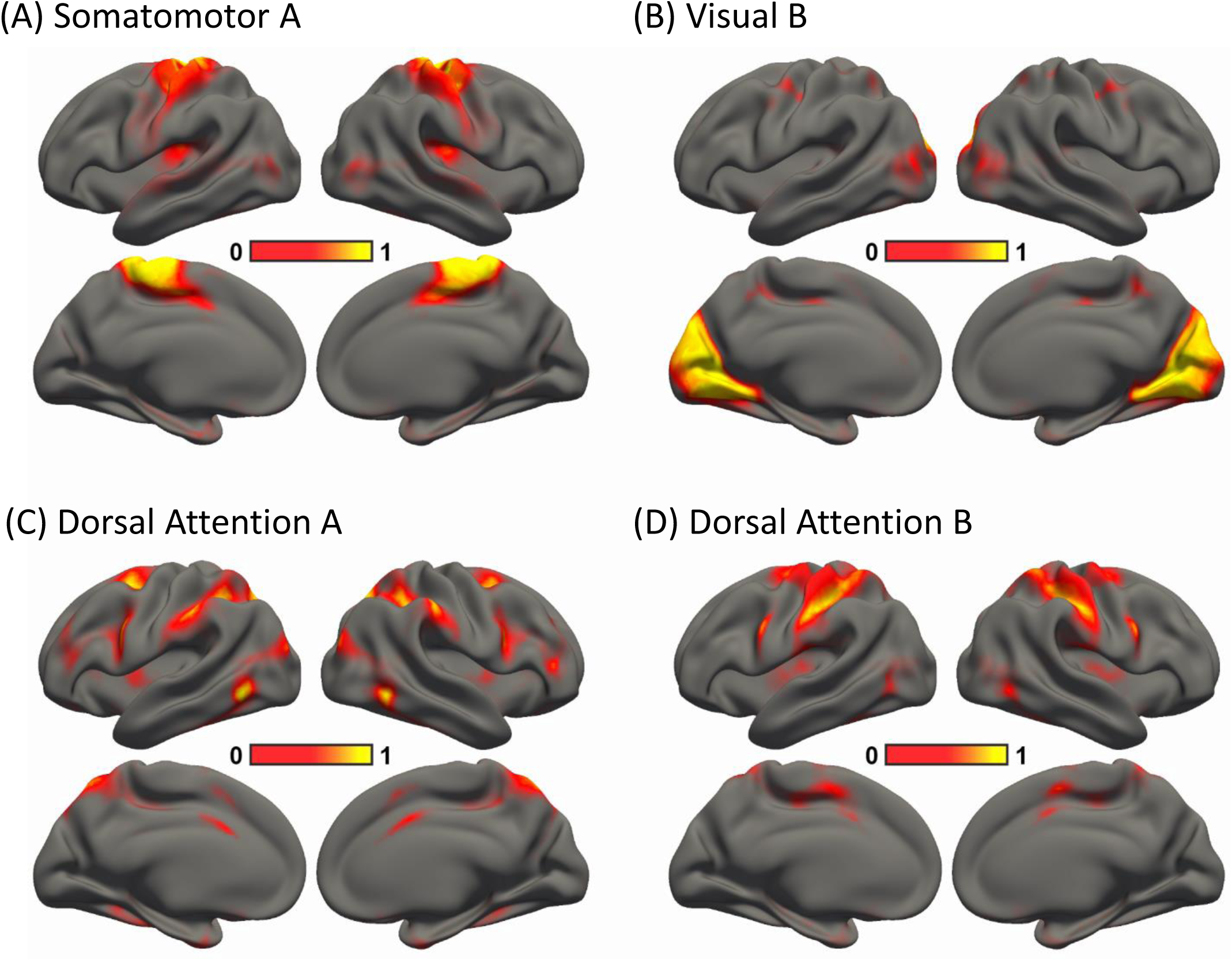
Sensory-motor networks are less spatially variable than association networks across subjects in the HCP training set. Spatial probability maps of (A) Somatomotor network A, (B) Visual network B, (C) Dorsal Attention network A, and (D) Dorsal Attention network B. A higher value (bright color) at a spatial location indicates high probability of a network appearing at that spatial location. Results were replicated in the GSP (Figure S4) and Corr-HNU (Figure S5) datasets. Note that this corresponds to the *Θ*_*l*_ parameter in Figure 1.

**Figure S4.**
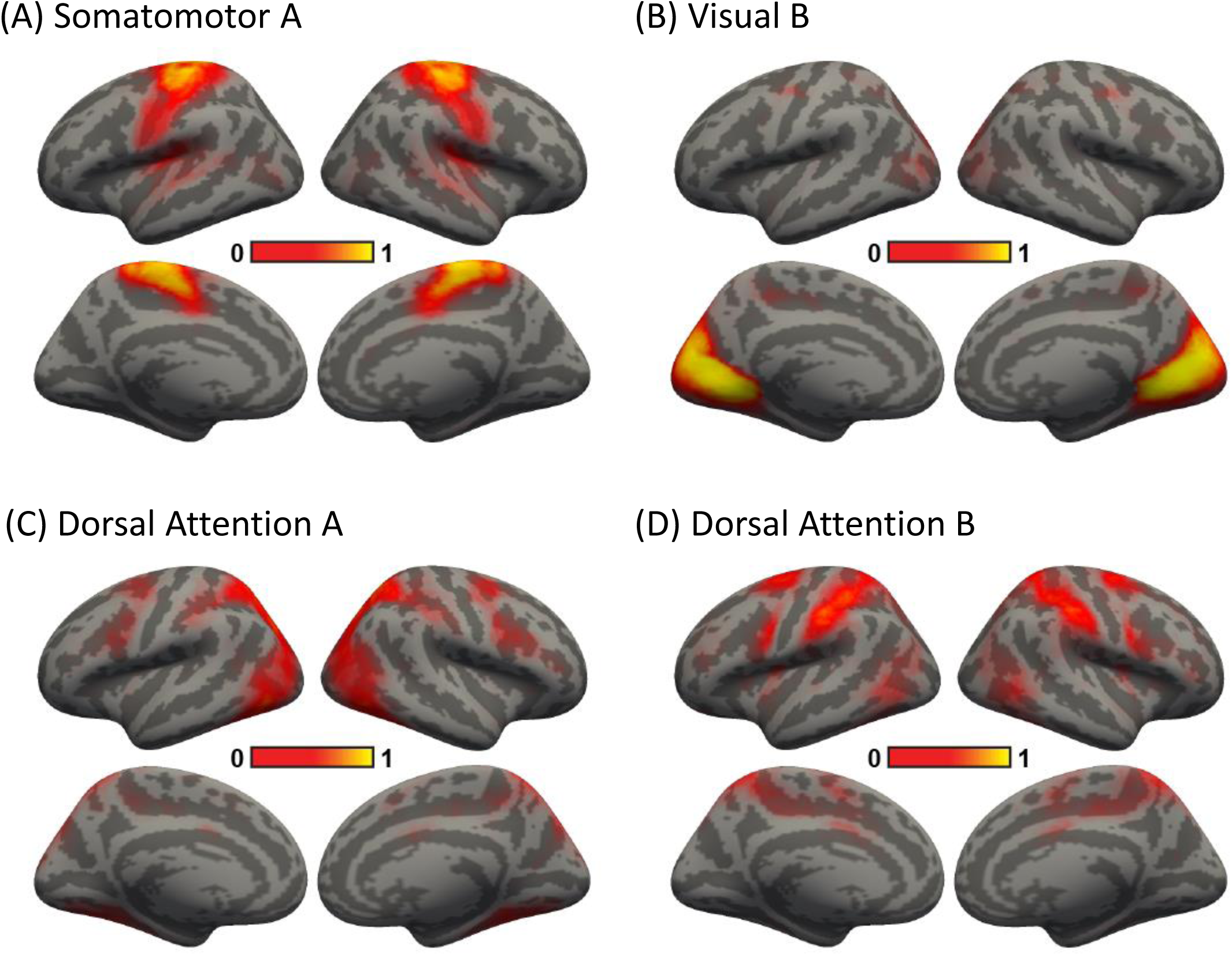
Sensory-motor networks are less spatially variable than association networks across subjects in the GSP dataset. Spatial probability maps of (A) Somatomotor network A, (B) Visual network B, (C) Dorsal Attention network A, and (D) Dorsal Attention network

B. A higher value (bright color) at a spatial location indicates high probability of a network appearing at that spatial location. Note that this corresponds to the *Θ*_*l*_ parameter in Figure 1.

**Figure S5.**
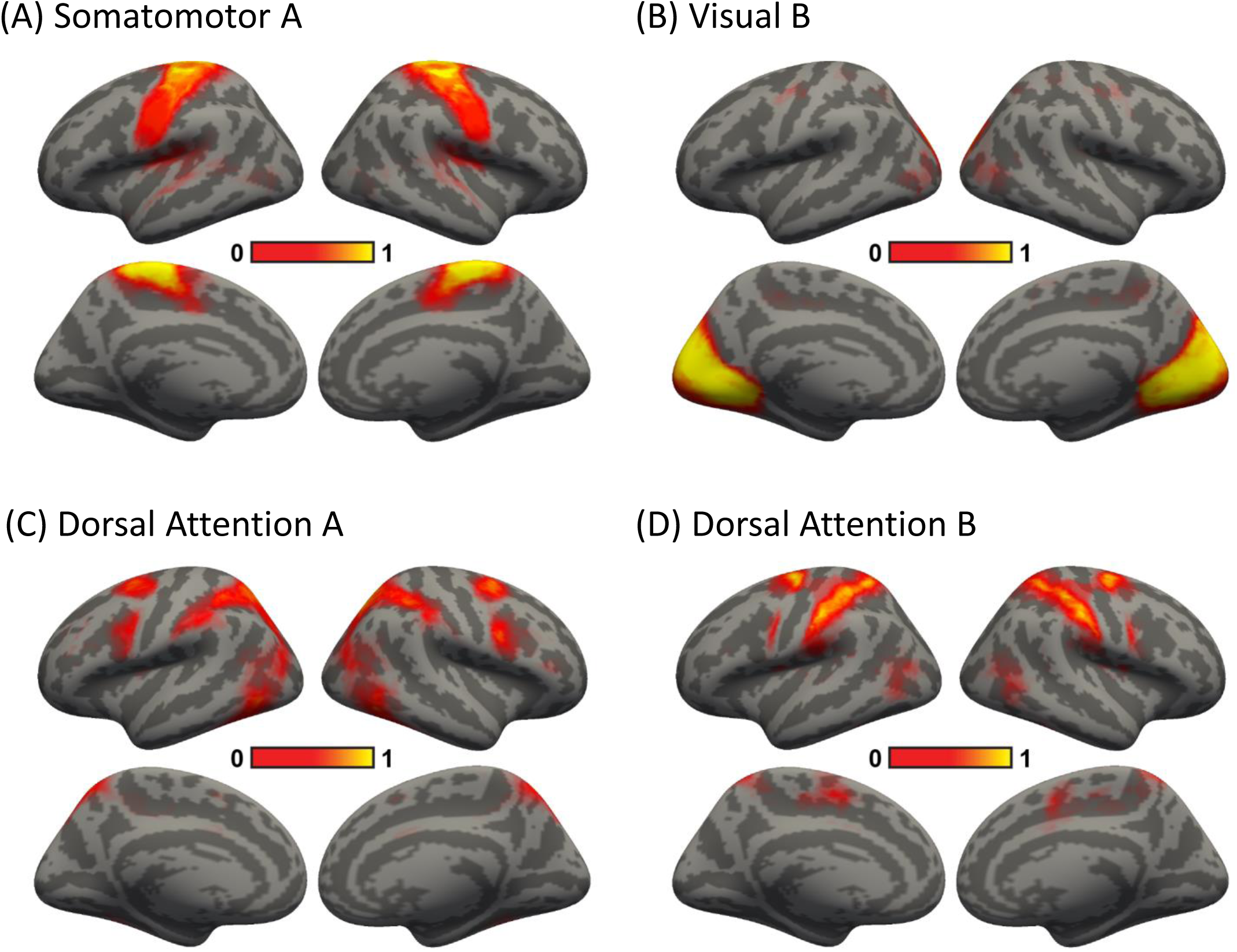
Sensory-motor networks are less spatially variable than association networks across subjects in the CoRR-HNU dataset. Spatial probability maps of (A) Somatomotor network A, (B) Visual network B, (C) Dorsal Attention network A, and (D) Dorsal Attention network B. A higher value (bright color) at a spatial location indicates high probability of a network appearing at that spatial location. Note that this corresponds to the *Θ*_*l*_ parameter in Figure 1.

**Figure S6.**
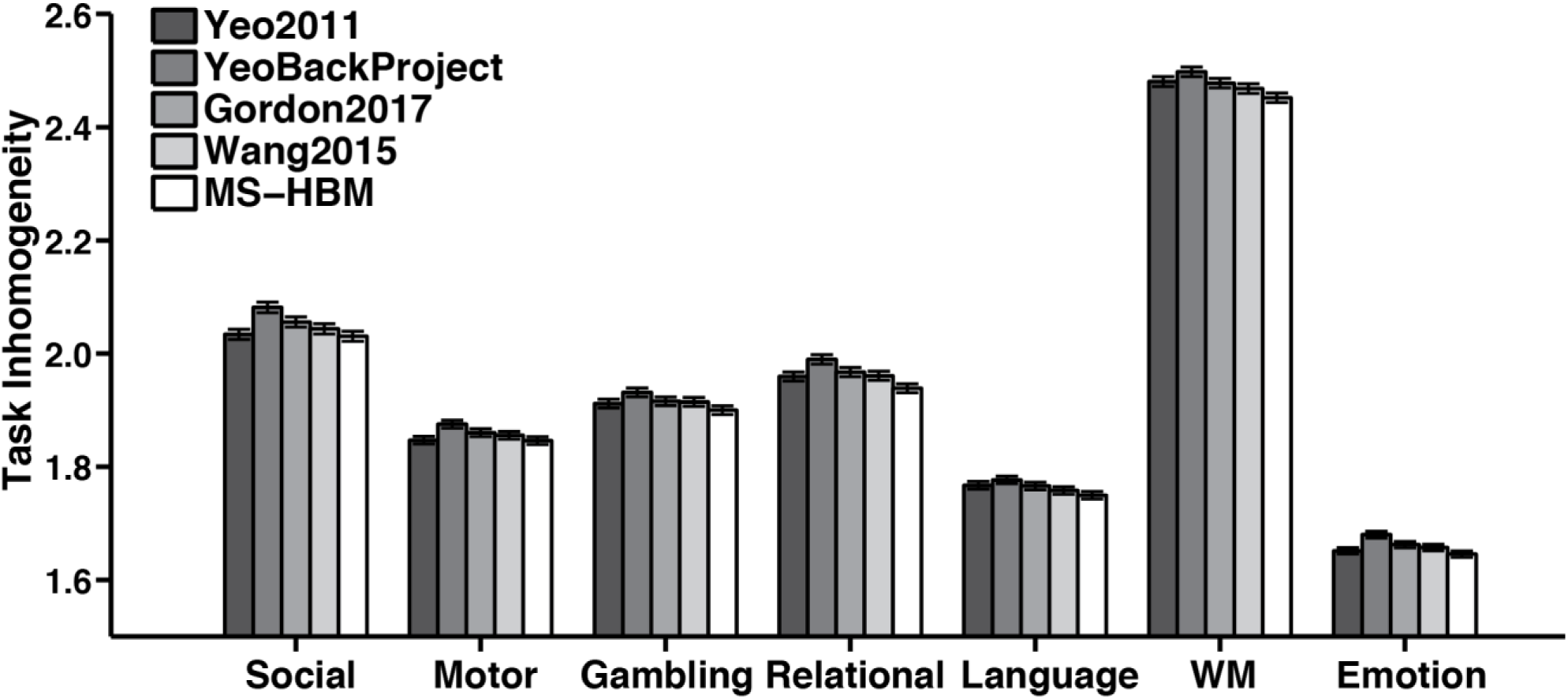
Task inhomogeneity of resting-state parcellations in the HCP dataset. 17-network individual-specific parcellations were estimated using one rs-fMRI session. Task inhomogeneity was then defined as the standard deviation of task activation within each network, and then averaged across all networks and contrasts within each behavioral domain. Lower value indicates better functional homogeneity. Error bars correspond to standard errors. Compared with Yeo2011, YeoBackProject, Gordon2017 and Wang2015, the MS-HBM individual-specific parcellations achieved a modest average improvement 0.63% (Cohen’s d = 0.12, 0.09, 0.66, 1.0, 0.9, 1.1, 0.46 for social, motor, gambling, relational, language, working memory and emotion respectively), 2.0% (Cohen’s d > 1.3 for all domains), 1.04% (Cohen’s d > 0.99 for all domains) and 0.7% (Cohen’s d > 0.79 for all domains) respectively.

**Figure S7.**
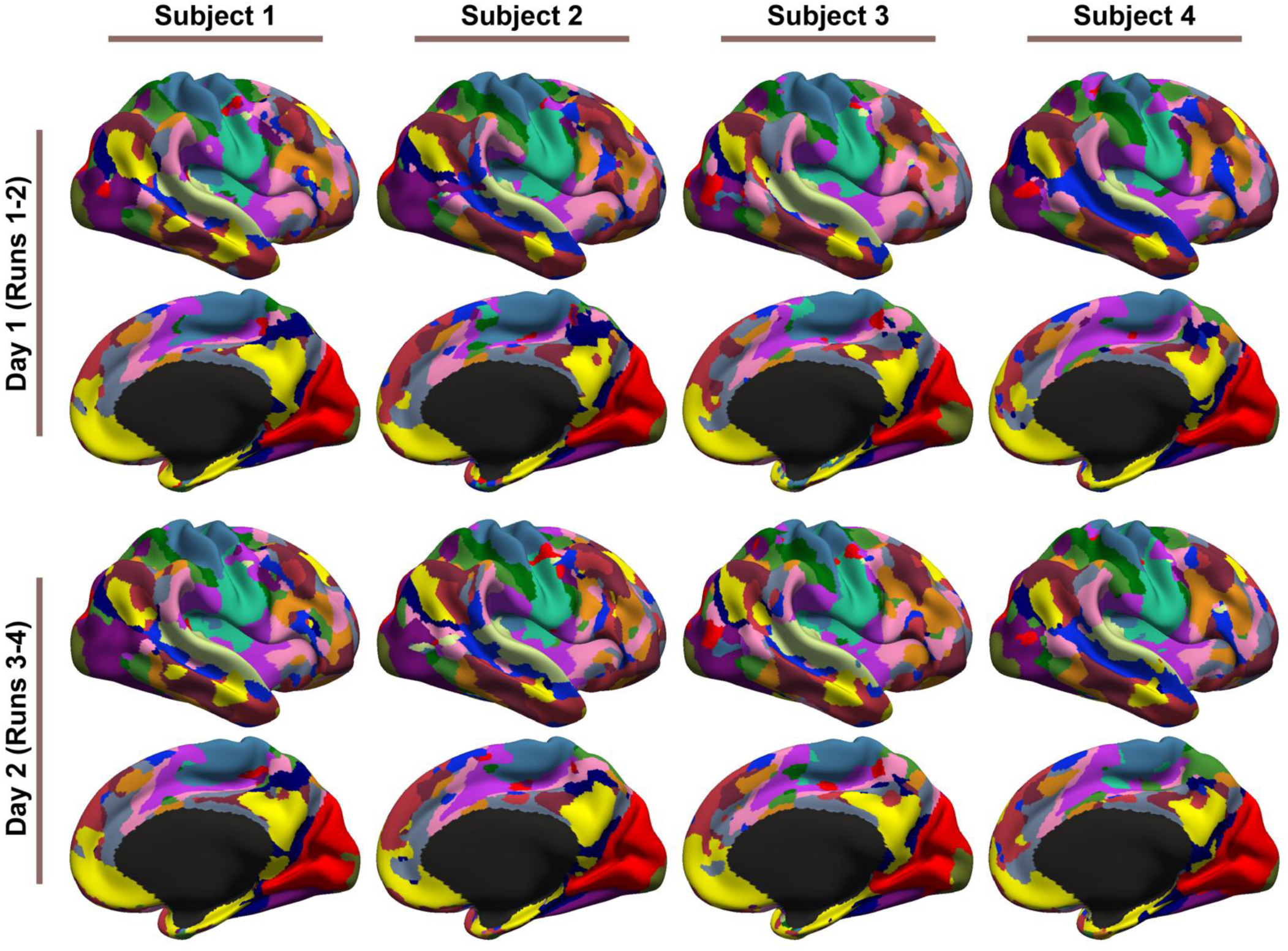
17-network parcellations were estimated using runs 1-2 and runs 3-4 separately for each subject from the HCP test set. Parcellations of four representative subjects are shown here. Left hemisphere parcellations are shown in Figure 4.

**Figure S8.**
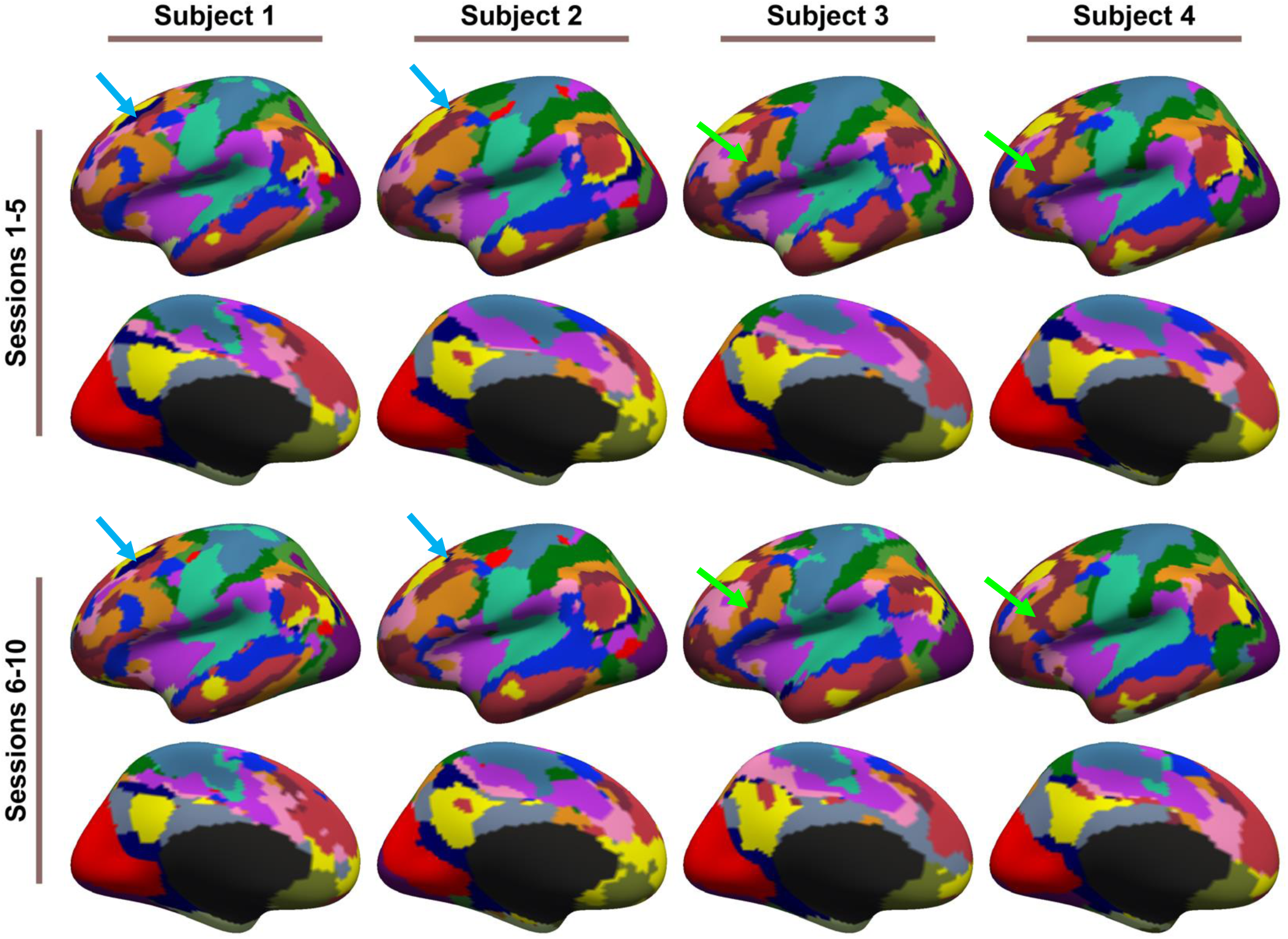
17-network parcellations were estimated using sessions 1-5 and sessions 6-10 separately for each subject from the CoRR-HNU dataset. Parcellations of four representative subjects are shown here. Black and green arrows indicate individual-specific parcellation features. The Default C (dark blue) network exhibited a dorsal prefrontal component for certain subjects (blue arrows), but was missing in other subjects. As another example, the lateral prefrontal component of the Control A (orange) network was separated into two separate components by the Control B (brown) network (green arrows). These features were mostly replicated across sessions. Right hemisphere parcellations are shown in Figure S9.

**Figure S9.**
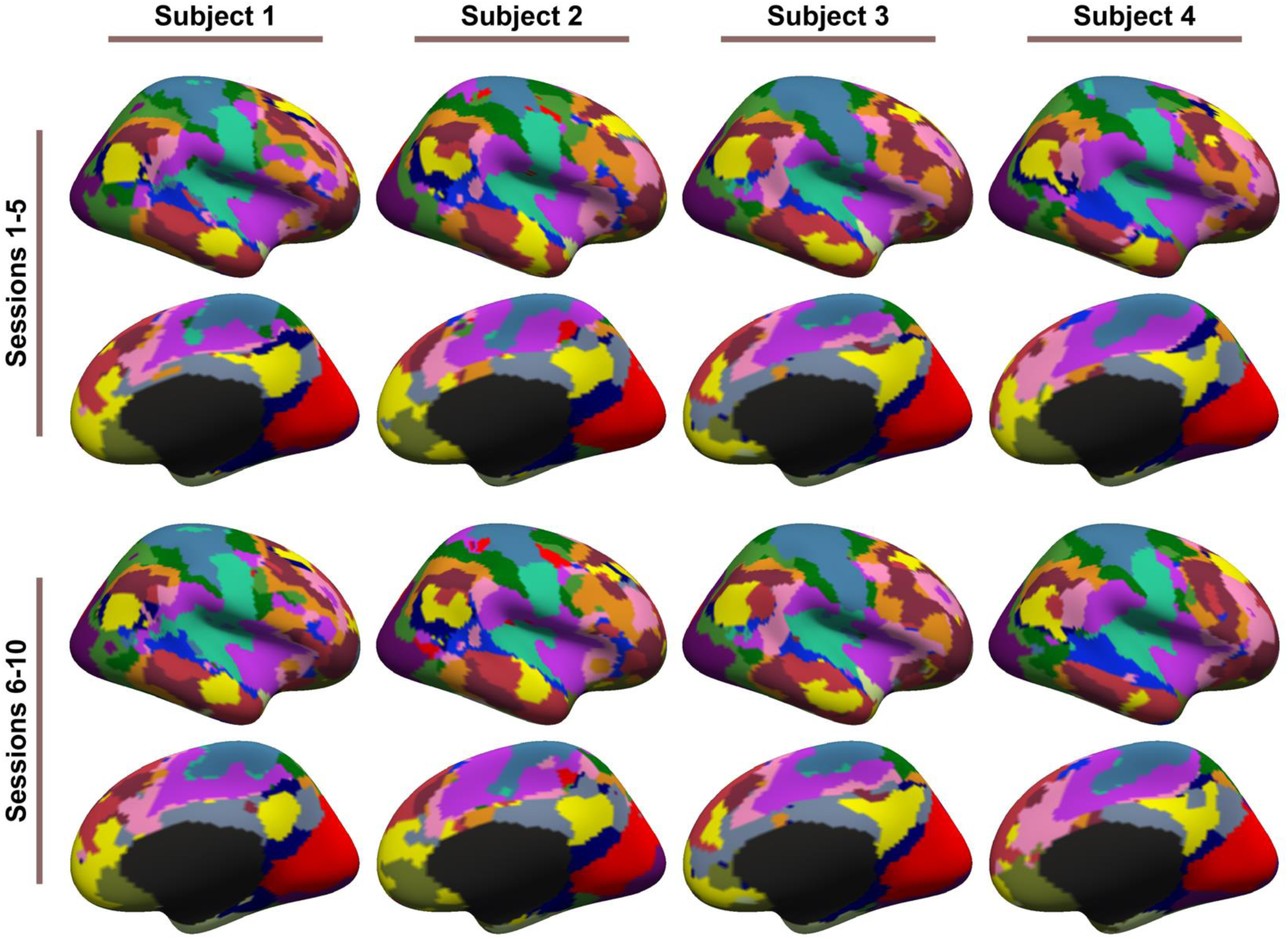
17-network parcellations were estimated using sessions 1-5 and sessions 6-10 separately for each subject from the CoRR-HNU dataset. Parcellations of four representative subjects are shown here. Left hemisphere parcellations are shown in Figure S8.

**Figure S10.**
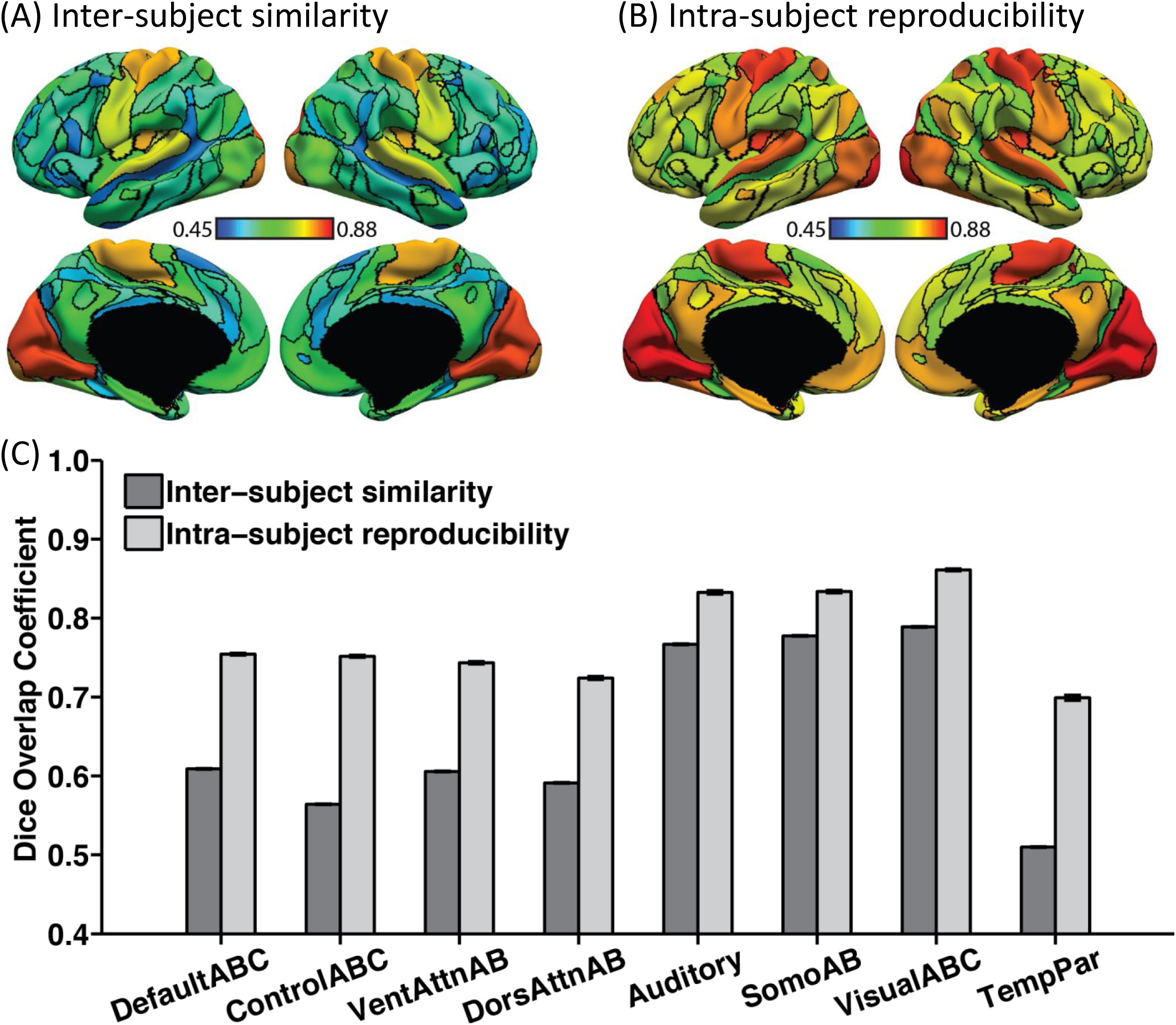
Individual-specific MS-HBM parcellations show high within-subject reproducibility and low across-subject similarity in the HCP test set. Individual-specific MS-HBM parcellations were generated by using the first two runs (day 1) and last two runs (day 2) separately for each subject. (A) Inter-subject spatial similarity for different networks. (B) Intra-subject reproducibility for different networks. Warm color indicates higher overlap. Cool color indicates lower overlap. (C) Quantification of inter-subject similarity and intra-subject reproducibility for different networks. “VentAttnAB” corresponds to Salience/Ventral Attention (A and B) networks. “SomoAB” corresponds to Somatomotor (A and B) networks. Error bars correspond to standard errors.

**Figure S11.**
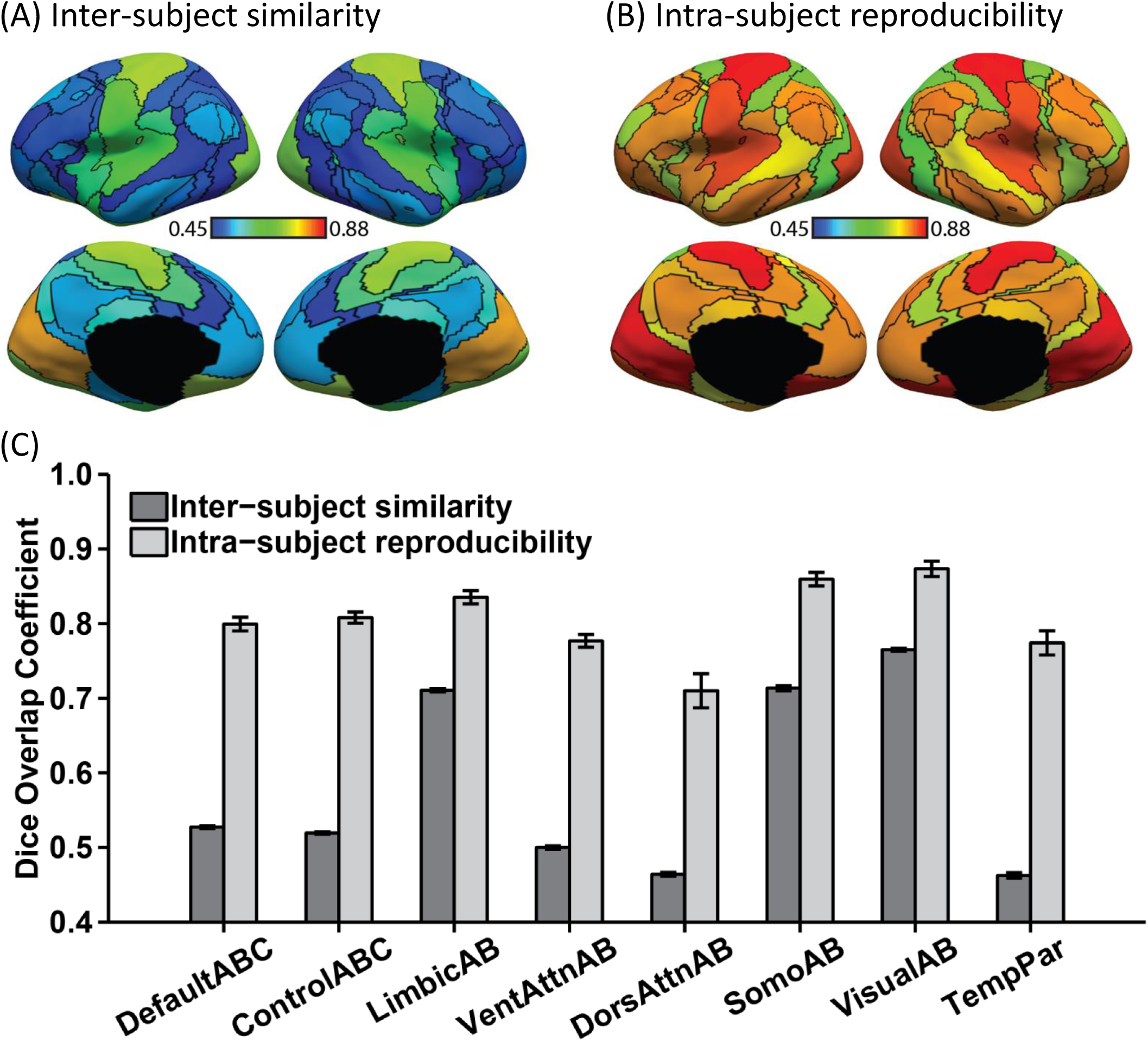
Individual-specific MS-HBM parcellations show high within-subject reproducibility (overlap = 81.6%) and low across-subject similarity (overlap = 59.4%) in the CoRR-HNU dataset. (A) Inter-subject spatial similarity for different networks. (B) Intra-subject reproducibility for different networks. Warm color indicates higher overlap. Cool color indicates lower overlap. (C) Quantification of inter-subject similarity and intra-subject reproducibility for different networks. “VentAttnAB” corresponds to Salience/Ventral Attention networks A and B. “SomoAB” corresponds to Somatomotor networks A and B. Error bars correspond to standard errors.

**Figure S12.**
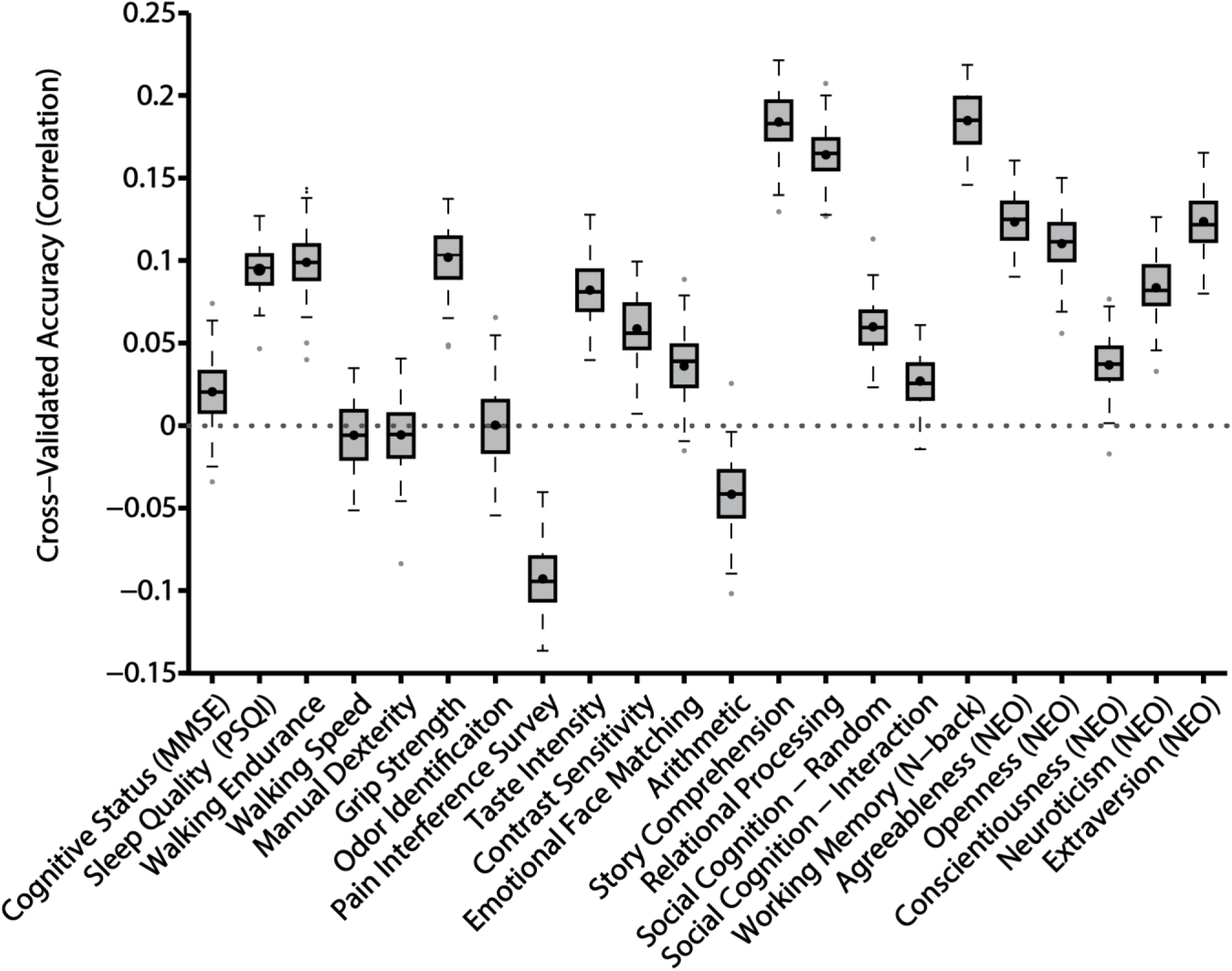
Prediction accuracy of 22 cognitive, emotion, personality and other non-imaging measures based on inter-subject differences in the spatial arrangement of cortical networks. Boxplots utilized default Matlab parameters, i.e., box shows median and inter-quartile range (IQR). Whiskers indicate 1.5 IQR. Dot indicates mean. In the case of the NEO-5 personality scores, average predication accuracy was r = 0.0955 ± 0.0085 (mean ± std). Other measures are found in Figures 5 and S13.

**Figure S13.**
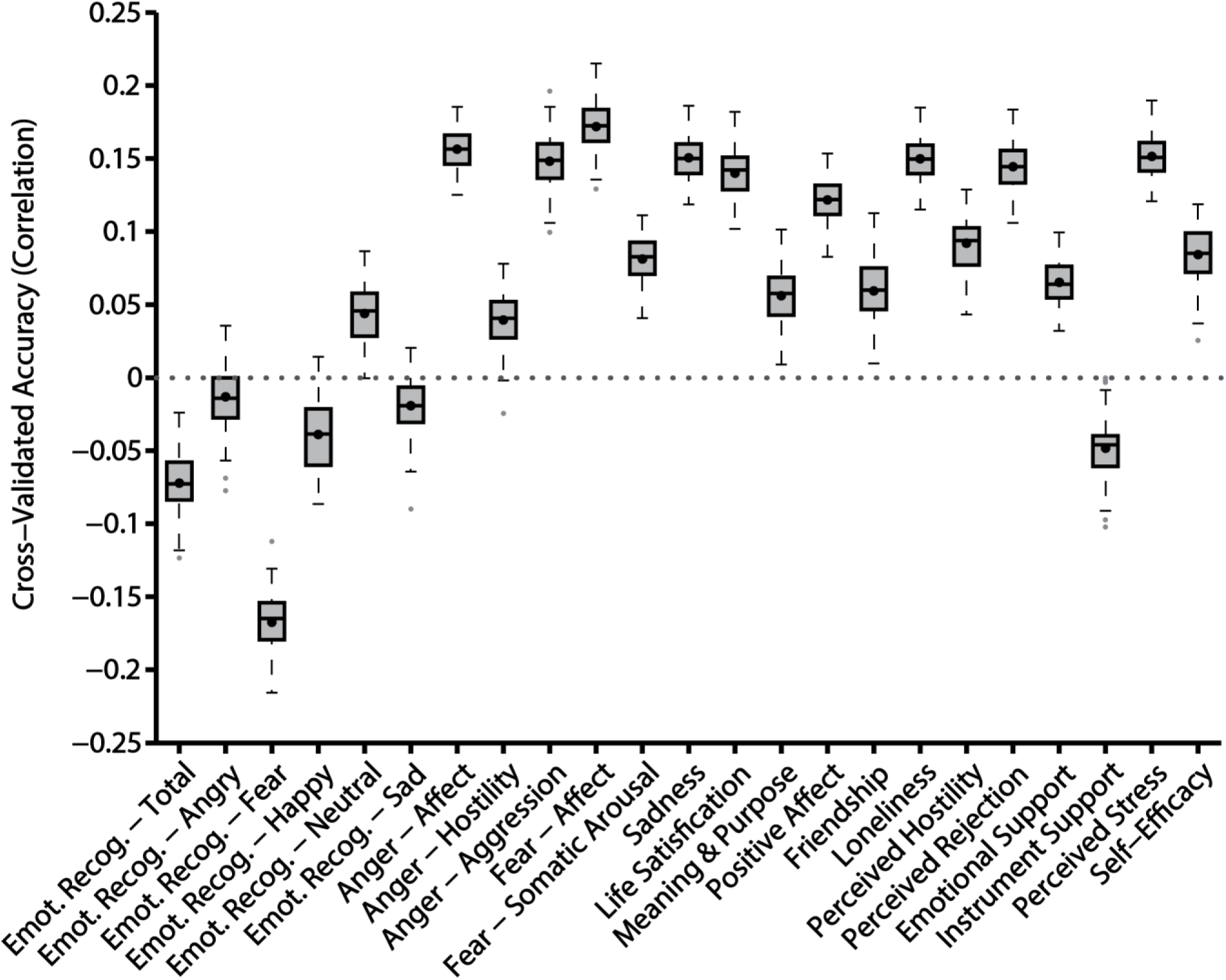
Prediction accuracy of 23 cognitive, emotion, personality and other non-imaging measures based on inter-subject differences in the spatial arrangement of cortical networks. Boxplots utilized default Matlab parameters, i.e., box shows median and inter-quartile range (IQR). Whiskers indicate 1.5 IQR. Dot indicates mean. In the case of the emotional measures (all items in Figure S15 except for emotional recognition), the average prediction accuracy was r = 0.1038 ± 0.0070 (mean ± std). Other measures are found in Figures 5 and S12. Interestingly, prediction accuracy for the emotion recognition task was poor.

**Figure S14.**
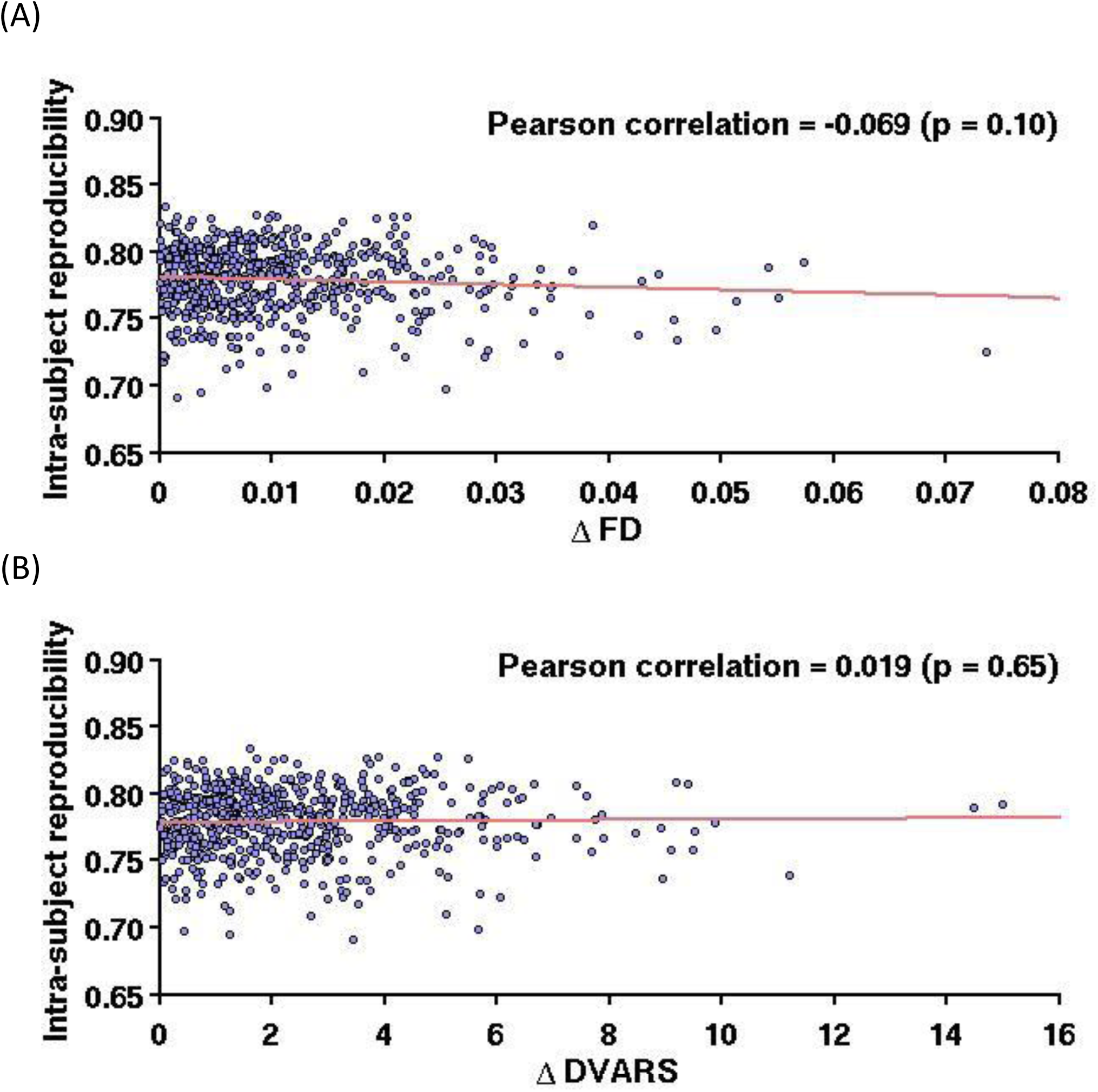
Intra-subject reproducibility is not significantly affected by FD or DVARS difference between the two scan days of the HCP subjects. (A) Scatterplot of intra-subject reproducibility versus absolute FD difference between the two scan days. (B) Scatterplot of intra-subject reproducibility versus absolute DVARS difference between the two scan days. Only HCP test set subjects with all 58 behavioral measures were considered. If network topography was corrupted by motion-related imaging artifacts, then one would expect subjects who moved by very different amounts on the two different scan days to have poorer intra-subject parcellation reproducibility than subjects who moved by similar amounts on both days. As shown above, this is not the case.

**Figure S15.**
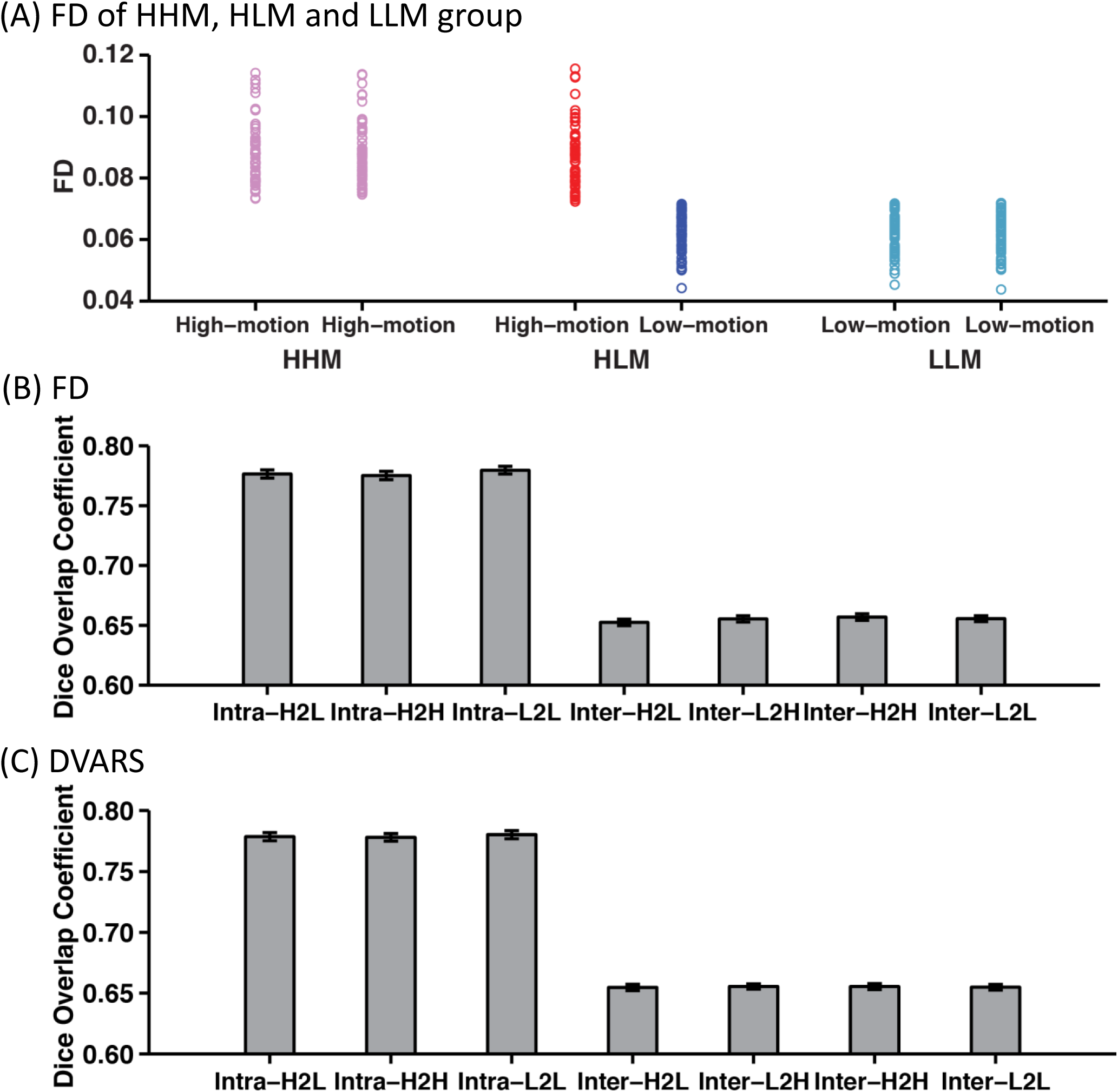
Motion-related imaging artifacts have little effect on intra-subject reproducibility and inter-subject similarity. (A) One issue with the previous analysis (Figure S14) is that a subject with high absolute FD difference between the two scan days might still exhibit low motion on both days (relative to other subjects). Therefore, we also considered three groups of 60 HCP subjects. The high-low-motion (HLM) subjects consisted of subjects, whose FD were above the median FD in one day, and below the median FD in another day. The high-high-motion (HHM) group consisted of subjects whose FD were high on both days and matched the FD of the HLM subjects on the high FD days. The low-low-motion (LLM) motion group consisted of subjects whose FD were low on both days and matched the FD of the HLM subjects on the low FD days. (B) Intra-subject parcellation reproducibility were basically identical among HLM subjects (Intra-H2L), HHM subjects (Intra-H2H) and LLM subjects (Intra-L2L). For comparisons, the inter-subject parcellation similarity between HHM subjects and HLM subjects during their high motion days (Inter-H2H), between LLM subjects and HLM subjects during their low motion days (Inter-L2L), between HHM subjects and HLM subjects during their low motion days (Inter-H2L) and between LLM subjects and HLM subjects during their high motion days (Inter-L2H) were basically identical and significantly lower than intra-subject parcellation reproducibility. (C) Similar results were obtained with DVARS.

**Figure S16.**
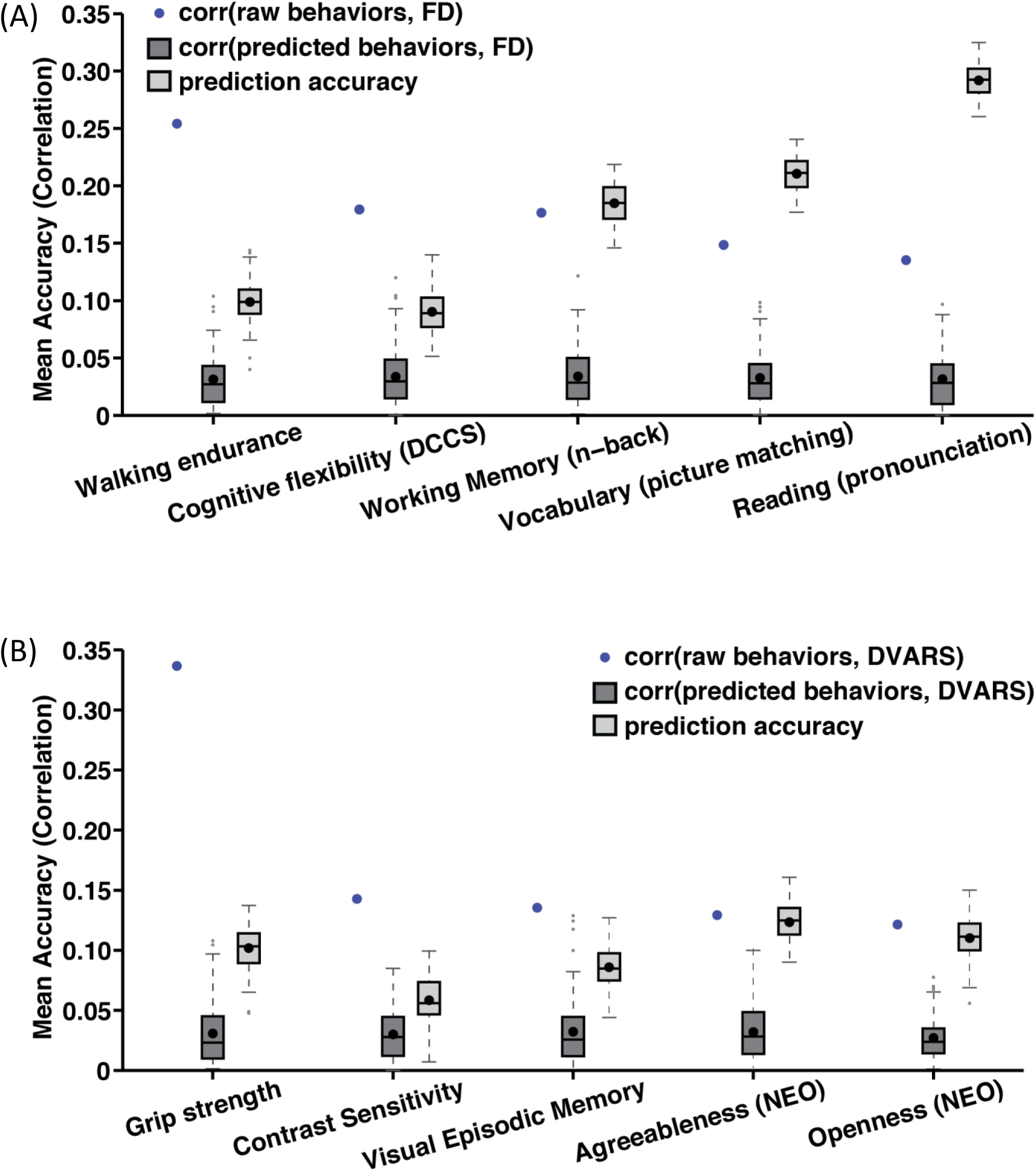
Prediction accuracies of 5 behavioral measures with the highest correlation with (A) FD and (B) DVARS. The correlation between the behavioral measures and FD/DVARS was depicted as the blue dot. Boxplots utilized default Matlab parameters, i.e., box shows median and inter-quartile range (IQR). Whiskers indicate 1.5 IQR. Dot indicates mean. The prediction accuracies of these 5 behaviors were higher than the correlation between the prediction of these 5 behavioral measures and FD/DVARS.

**Figure S17.**
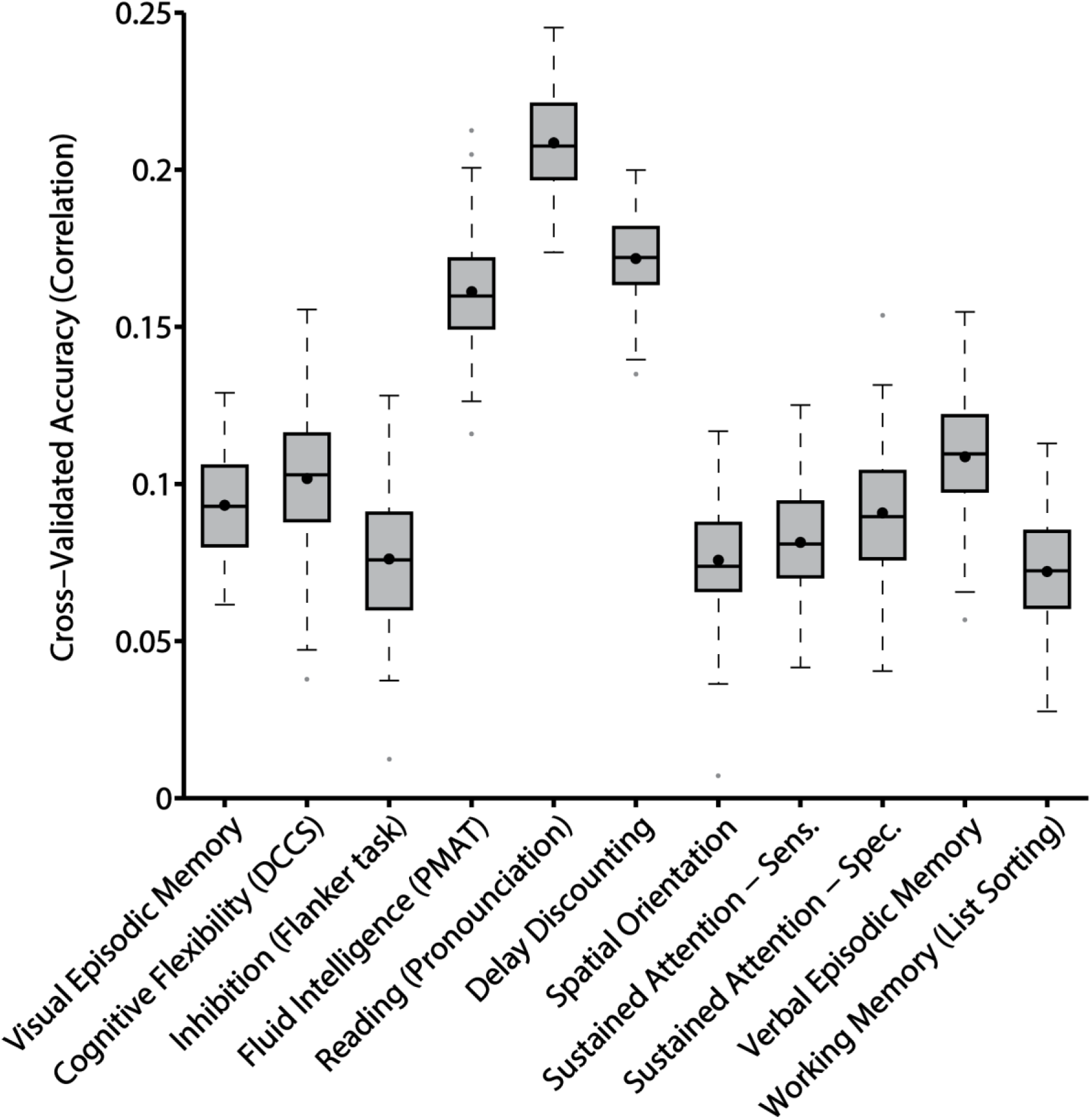
Prediction accuracy of 11 cognitive measures based on inter-subject differences in the spatial arrangement of task-relevant cortical networks. Boxplots utilized default Matlab parameters, i.e., box shows median and inter-quartile range (IQR). Whiskers indicate 1.5 IQR. Dot indicates mean. Average prediction accuracy based only on task-relevant networks was r = 0.1129 ± 0.0062 (mean ± std).

1 We note that with uniformly distributed ROIs, spatially extensive networks might be over-represented because they contribute disproportionately more ROIs to the computation of the connectivity profiles. However, regions within these spatially extensive networks do not necessarily have homogeneous connectivity patterns, so the uniformly distributed ROIs can capture these heterogeneous connectivity patterns. Furthermore, given that we do not know the networks a-priori, it would be challenging to define the ROIs a-priori, such that large patches of homogeneous regions would not be over-represented.

2 Conceptually, κ is estimated by averaging information across all vertices of all subjects. ∈_1:*L*_, σ_1:*L*_ and 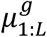 are estimated by averaging information across all vertices within each network across all subjects, 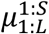 and 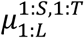 are estimated by averaging information across all vertices within each network for each subject, while *Θ*_1:*N*,1:*L*_ is estimated for each vertex by averaging information across all subjects. On the other hand, the spatial smoothness *V* (parameterized by c) serves to “clean up” individual-specific parcellations by removing isolated islands of vertices assigned to particular networks. Because these isolated islands constitute only a small fraction of the networks, excluding the spatial smoothness *V* (i.e., set *c* = 0) will not significantly affect the estimates of ∈_1:*L*_, σ_1:*L*_, *Θ*_1:*N*,1:*L*._

